# Study on potential differentially expressed genes in idiopathic pulmonary fibrosis by bioinformatics and next generation sequencing data analysis

**DOI:** 10.1101/2023.09.18.558229

**Authors:** Basavaraj Vastrad, Chanabasayya Vastrad

## Abstract

Idiopathic pulmonary fibrosis (IPF) is a chronic progressive lung disease with reduced quality of life and earlier mortality, but its pathogenesis and key genes are still unclear. In this investigation, bioinformatics was used to deeply analyze the pathogenesis of IPF and related key genes, so as to investigate the potential molecular pathogenesis of IPF and provide guidance for clinical treatment. Next generation sequencing dataset GSE213001 was obtained from Gene Expression Omnibus (GEO), the differentially expressed genes (DEGs) were identified between IPF and normal control group. The DEGs between IPF and normal control group were screened with the DESeq2 package of R language. The Gene Ontology (GO) and REACTOME pathway enrichment analyses of the DEGs were performed.. Using the g:Profiler, the function and pathway enrichment analysis of DEGs were performed. Then, a protein–protein interaction (PPI) network was constructed via the Integrated Interactions Database (IID) database. Cytoscape with Network Analyzer was used to identify the hub genes. miRNet and NetworkAnalyst database was used to construct the targeted microRNAs (miRNAs) and transcription factors (TFs) of the hub genes. Finally receiver operating characteristic (ROC) curve analysis was used to validates the hub genes. A total of 958 DEGs were screened out in this study, including 479 up regulated genes and 479 down regulated genes. Most of the DEGs were significantly enriched in response to stimulus, GPCR ligand binding, microtubule-based process and defective GALNT3 causes HFTC. In combination with the results of the PPI network, miRNA-hub gene regulatory network and TF-hub gene regulatory network, hub genes including LRRK2, BMI1, EBP, MNDA, KBTBD7, KRT15, OTX1, TEKT4, SPAG8 and EFHC2 were selected. Our findings will contribute to identification of potential biomarkers and novel strategies for the treatment of IPF, and provide a novel strategy for clinical therapy.

## Introduction

Lung fibrosis is a progressive, chronic, and irreversible fibrosing interstitial lung disease; it affects 2 - 9 peoples per 100,000 people’s worldwide [1]. It is also known as idiopathic pulmonary fibrosis (IPF) [2]. It is characterized by clinical symptoms of cough and dyspnea, declining pulmonary function with impaired gas exchange, and progressive lung scarring [3]. IPF increases the complications of developing pulmonary hypertension [4], lung cancer [5], diabetes mellitus [6], dermatomyositis [7], polymyositis [8], systemic sclerosis [9], mixed connective tissue disease [10], systemic lupus erythematosus [11], rheumatoid arthritis [12], sarcoidosis [13], scleroderma [14], pneumonia [15], heart failure [16], obesity [17], viral respiratory diseases [18], gastroesophageal reflux disease [19], chronic obstructive pulmonary disease [20] and airway inflammation [21]. Genetic, aging and environmental factors are thought to be the contributing factors to IPF [22]. Therefore, exploring the possible molecular mechanisms of IPF is of great importance.

The recent bioinformatics and next generation sequencing (NGS) data analysis of specimens from sufferers and normal individuals enables us to investigate numerous diseases at diverse levels from somatic mutations and copy number variations to genomic expressions at the transcriptomic level [23–24]. Defining the molecular targets for diagnosis and reexamination is crucial for therapeutic action and prognostic outcome of IPF patients. Several investigations has described that significant molecular biomarkers and signaling pathways in IPF were identified as prognostic, diagnostic and therapeutic factors, such as p53 [25], TINF2 [26], ELMOD2 [27], TERT [28], ABCA3 [29], TGF-β signaling pathway [30], Smad and STAT3 signaling pathways [31], p38 MAPK signaling pathway [32], Wnt/β-Catenin signaling pathway [33] and JAK-STAT signaling pathway [34]. These finding suggested the important roles of some function biomarkers and signaling pathways in IPF progression. However, the diagnostic value of many biomarkers has not been investigated in IPF.

In this investigation, we aimed to identify novel diagnostic biomarkers for IPF based on bioinformatics and machine learning. We analyzed Gene Expression Omnibus (GEO) [https://www.ncbi.nlm.nih.gov/geo/] [35] NGS dataset (GSE213001) to determine DEGs between IPF and healthy control. The NGS data related to IPF were used to generate the differentially expressed genes (DEGs) involved in IPF. Moreover, the gene ontology (GO) term enrichment analysis, REACTOME pathway enrichment analysis, and construction of protein-protein interaction (PPI) network and modules, miRNA-hub gene regulatory network and TF-hub gene regulatory network were all performed. Furthermore, hub genes were subject to receiver operating characteristic (ROC) curve analysis. Therefore, our investigation provides a deep understanding of susceptibility genes in IPF and might provide appropriate drug targets for IPF therapies.

## Materials and Methods

### Next generation sequencing data source

NGS data of human mRNA about IPF research (GSE213001) were obtained from the GEO database. There were 180 samples in GSE213001, including 41 normal control samples without IPF and 98 experimental samples with IPF. All samples were detected through the Illumina HiSeq 3000 (Homo sapiens) platform.

### Identification of DEGs

The DESeq2 package [36] of R language was utilized to screen DEGs. The false discovery rate (FDR) of Benjamini and Hochberg (BH) method was applied to adjust p-values for multiple comparisons [37]. The significant differentially expressed cut-off was set as |logFC| > 0.512 for up regulated genes, |logFC| < - 0.831 for down regulated genes and adjusted P < 0.05. Finally, a heatmap was drawn to observe the clustering of samples. ggplot2 and gplot packages in R software was subsequently performed to plot the volcano plot and heatmap of DEGs.

### GO and pathway enrichment analyses of DEGs

GO and REACTOME pathway enrichment analyses of DEGs were performed via g:Profiler (http://biit.cs.ut.ee/gprofiler/) [38]. The GO enrichment analysis (http://www.geneontology.org) [39] consists of biological processes (BP), cellular components (CC) and molecular functions (MF). REACTOME (https://reactome.org/) [40] is a pathway database resource for understanding high-level biological functions and utilities. Gene count >2 and p < 0.05 were set as the threshold.

### Construction of the PPI network and module analysis

To further analyze the impacts of DEGs on IPF, the PPI network was constructed among various DEGs. Also, the online software Integrated Interactions Database (IID) (http://iid.ophid.utoronto.ca/search_by_proteins/) [41] was used to analyze the interactions of proteins encoded by DEGs. Then the Cytoscape software (V3.10.0; http://cytoscape.org/) [42] was utilized to visualize the PPI network. Hub genes were identified using Network Analyzer, a plug-in of Cytoscape software. Finally, the degree [43], betweenness [44], stress [45] and closeness [46] of each DEG was obtained by analyzing the topological structure of the PPI network. Significant modules in the PPI network were identified by PEWCC [47], another plug-in of Cytoscape software.

### Construction of the miRNA-hub gene regulatory network

The hub genes in PPI were selected as the promising targets for searching miRNA through the miRNet database (https://www.mirnet.ca/) [48]. This database contains miRNA-hub gene regulatory network data from 14 disparate sources including TarBase, miRTarBase, miRecords, miRanda (S mansoni only), miR2Disease, HMDD, PhenomiR, SM2miR, PharmacomiR, EpimiR, starBase, TransmiR, ADmiRE and TAM 2. The results of this process were arranged such that each entry was a specific miRNA-hub gene interaction associated with its source link. The identified miRNA-hub gene regulatory network was visualized using the Cytoscape software [42].

### Construction of the TF-hub gene regulatory network

The hub genes in PPI were selected as the promising targets for searching TF through the NetworkAnalyst database (https://www.networkanalyst.ca/) [49]. This database contains TF-hub gene regulatory network data from JASPAR. The results of this process were arranged such that each entry was a specific TF-hub gene interaction associated with its source link. The identified TF-hub gene regulatory network was visualized using the Cytoscape software [42].

### Receiver operating characteristic curve (ROC) analysis

The multivariate modeling with combined selected hub genes were used to identify mole with high sensitivity and specificity for IPF diagnosis. The receiver operator characteristic curves were plotted and area under curve (AUC) was determined independently to assess the conduct of each model using the R packages “pROC” [50]. AUC > 0.9 marked that the model had a good fitting effect.

## Results

### Identification of DEGs

A total of 958 DEGs were screened between normal control and IPF groups with |logFC| > 0.512 for up regulated genes, |logFC| < −0.831 for down regulated genes and adjusted P < 0.05, including 479 up regulated DEGs and 479 down regulated DEGs (Table 1 and Fig. 1). The heatmap of the DEGs has been shown in Fig. 2.

**Fig. 1.**
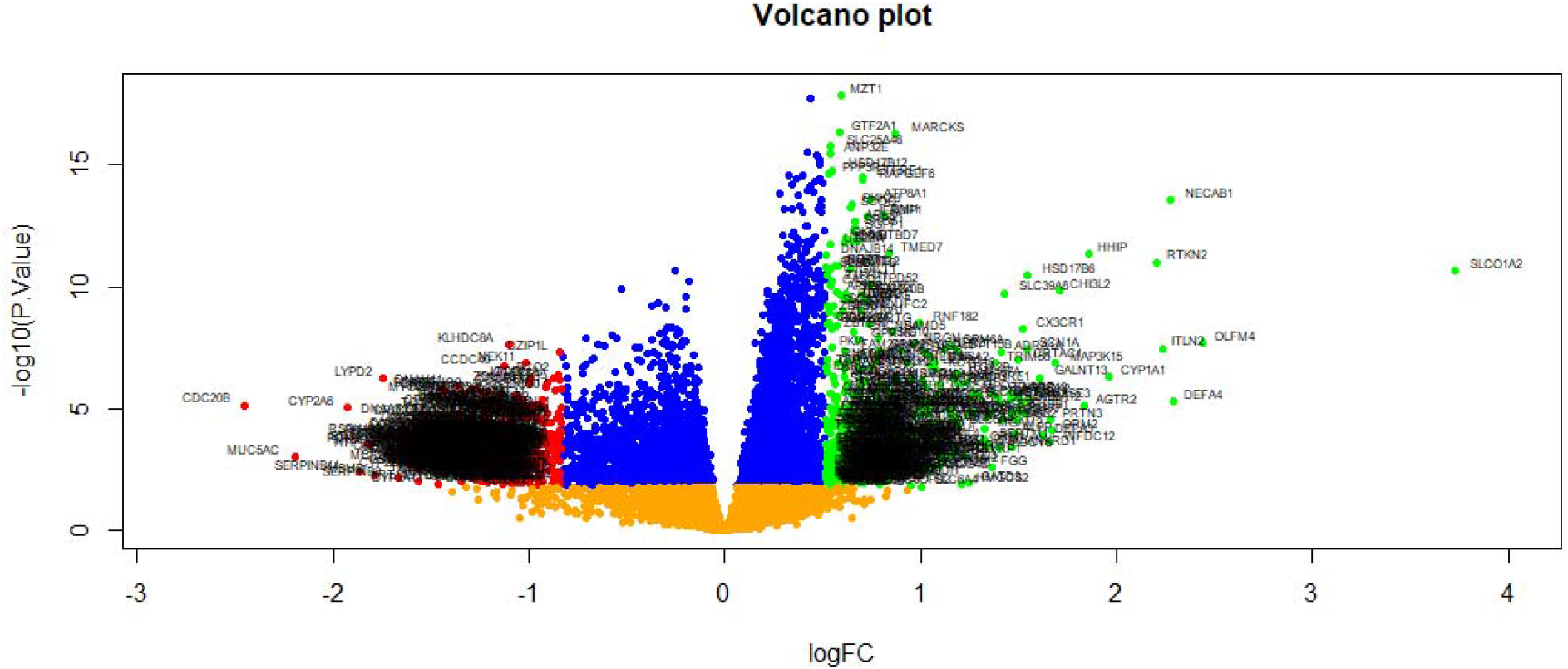
Volcano plot of differentially expressed genes. Genes with a significant change of more than two-fold were selected. Green dot represented up regulated significant genes and red dot represented down regulated significant genes.

**Fig. 2.**
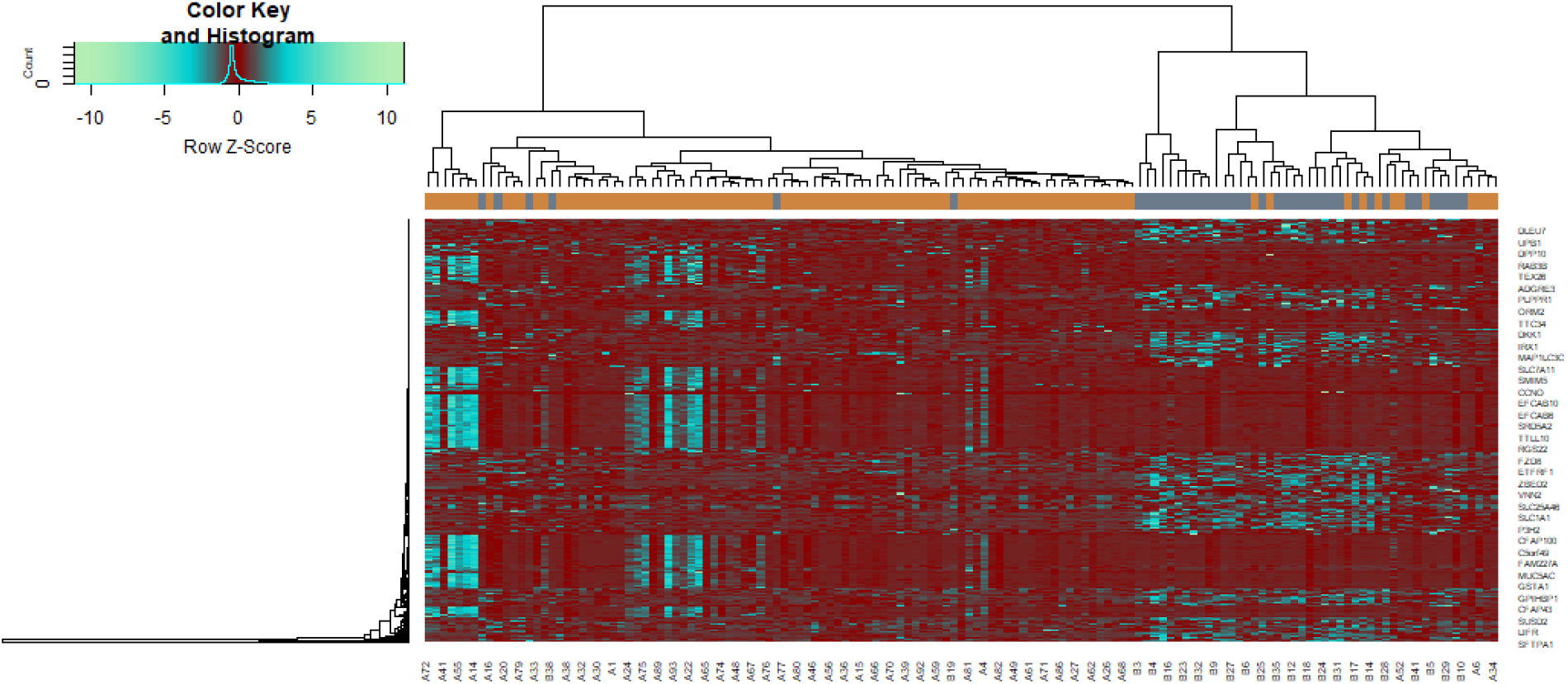
Heat map of differentially expressed genes. Legend on the top left indicate log fold change of genes. (A1 – A98 = IPF samples; B1 – B 41 = Normal control samples)

**Table 1.**
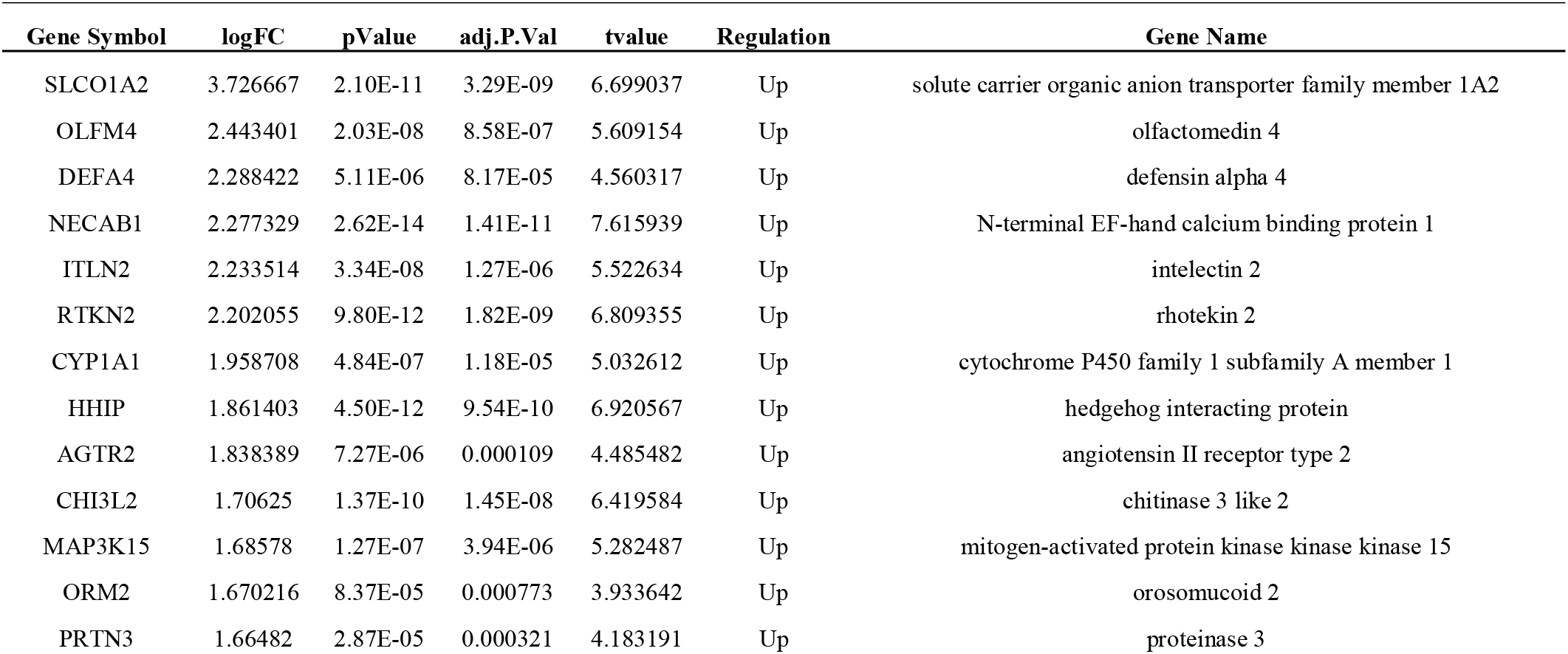
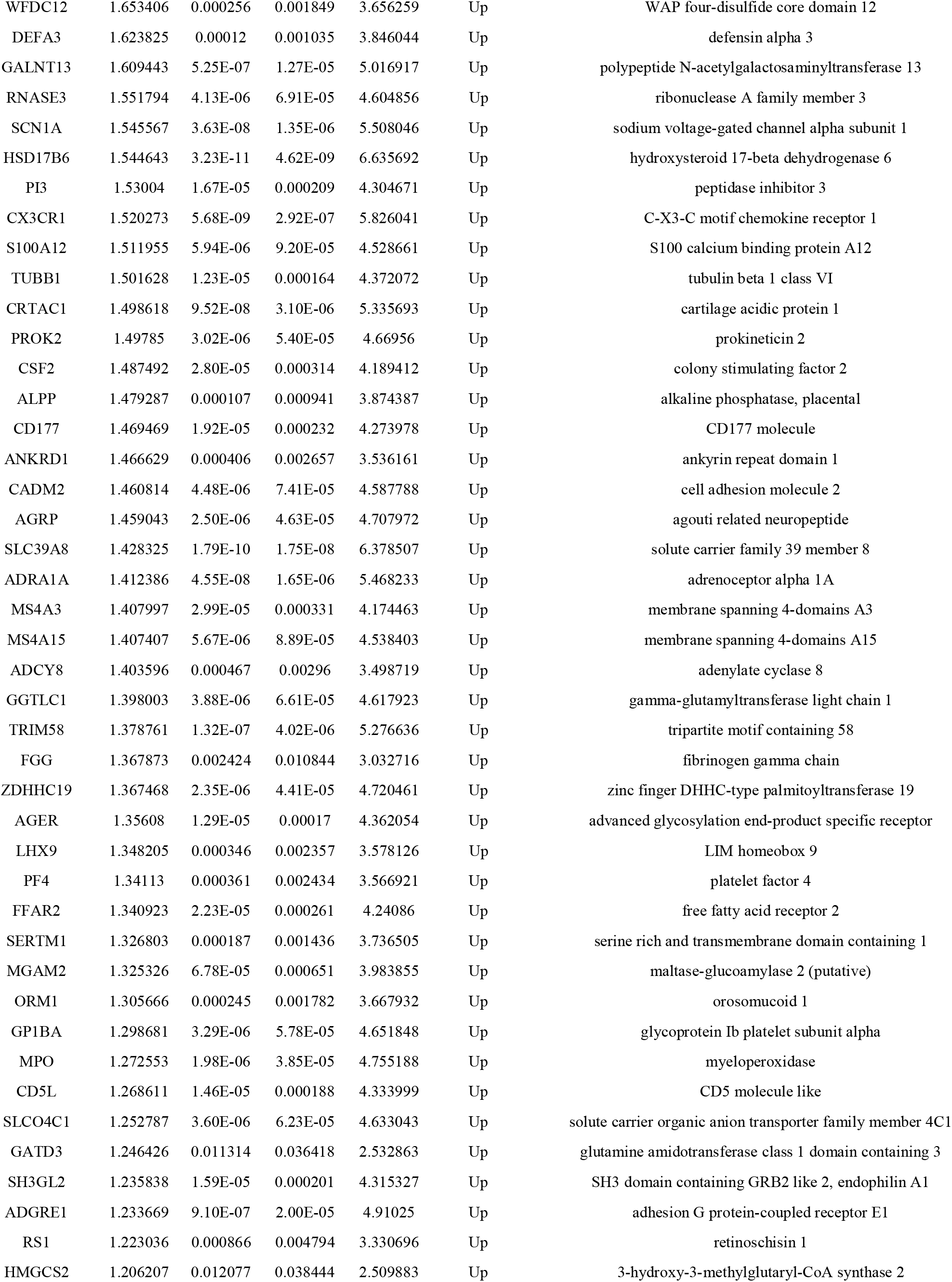
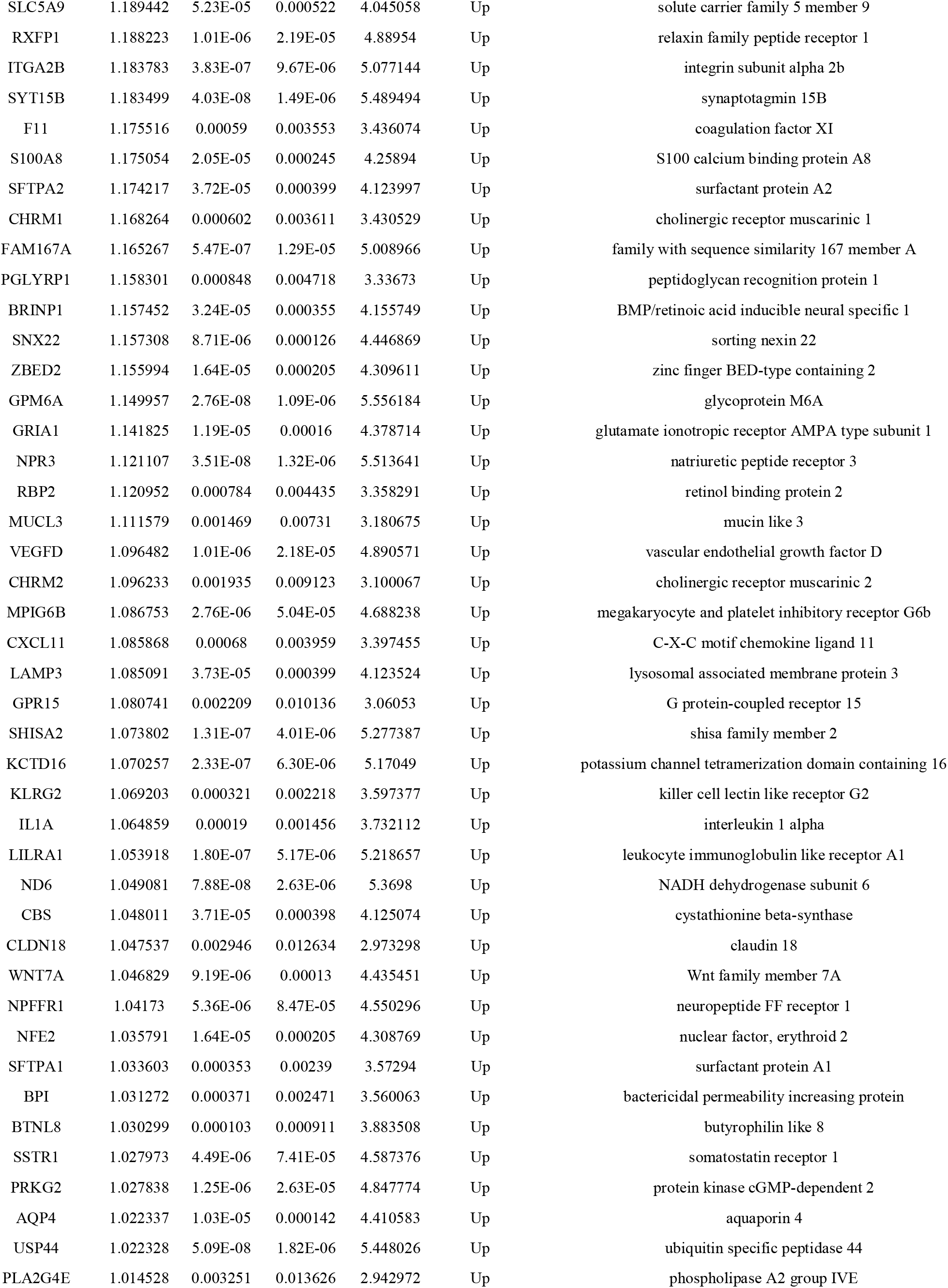
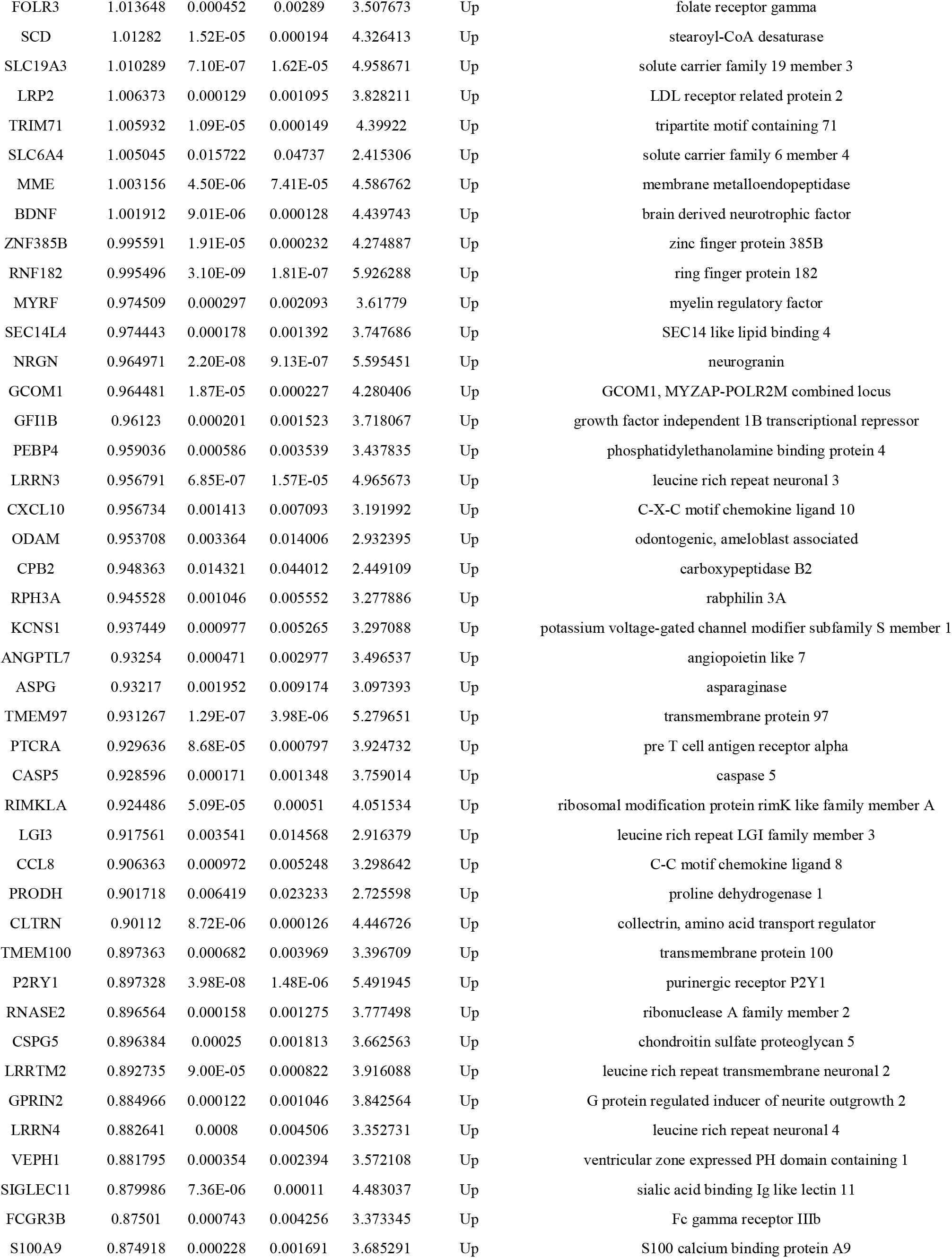
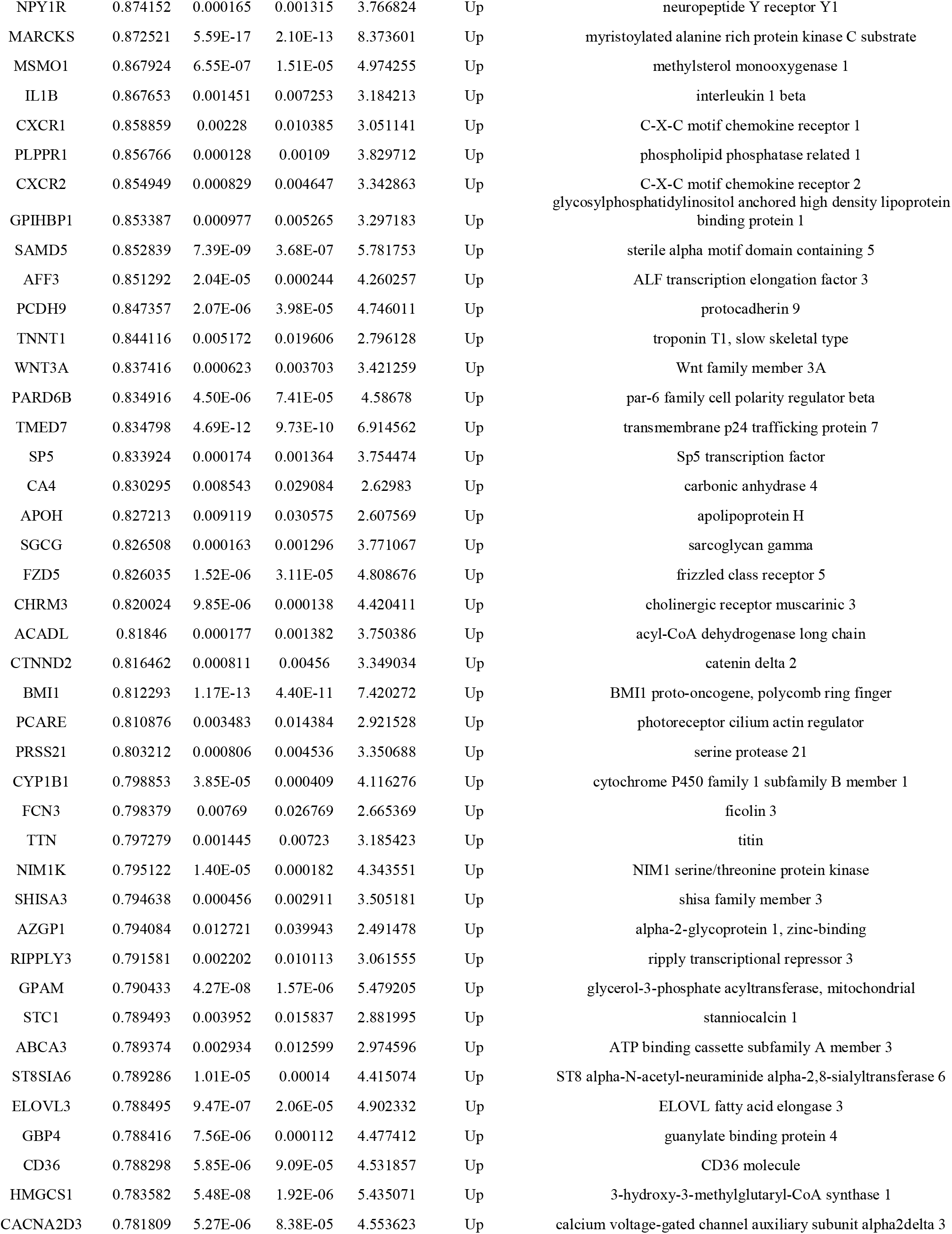
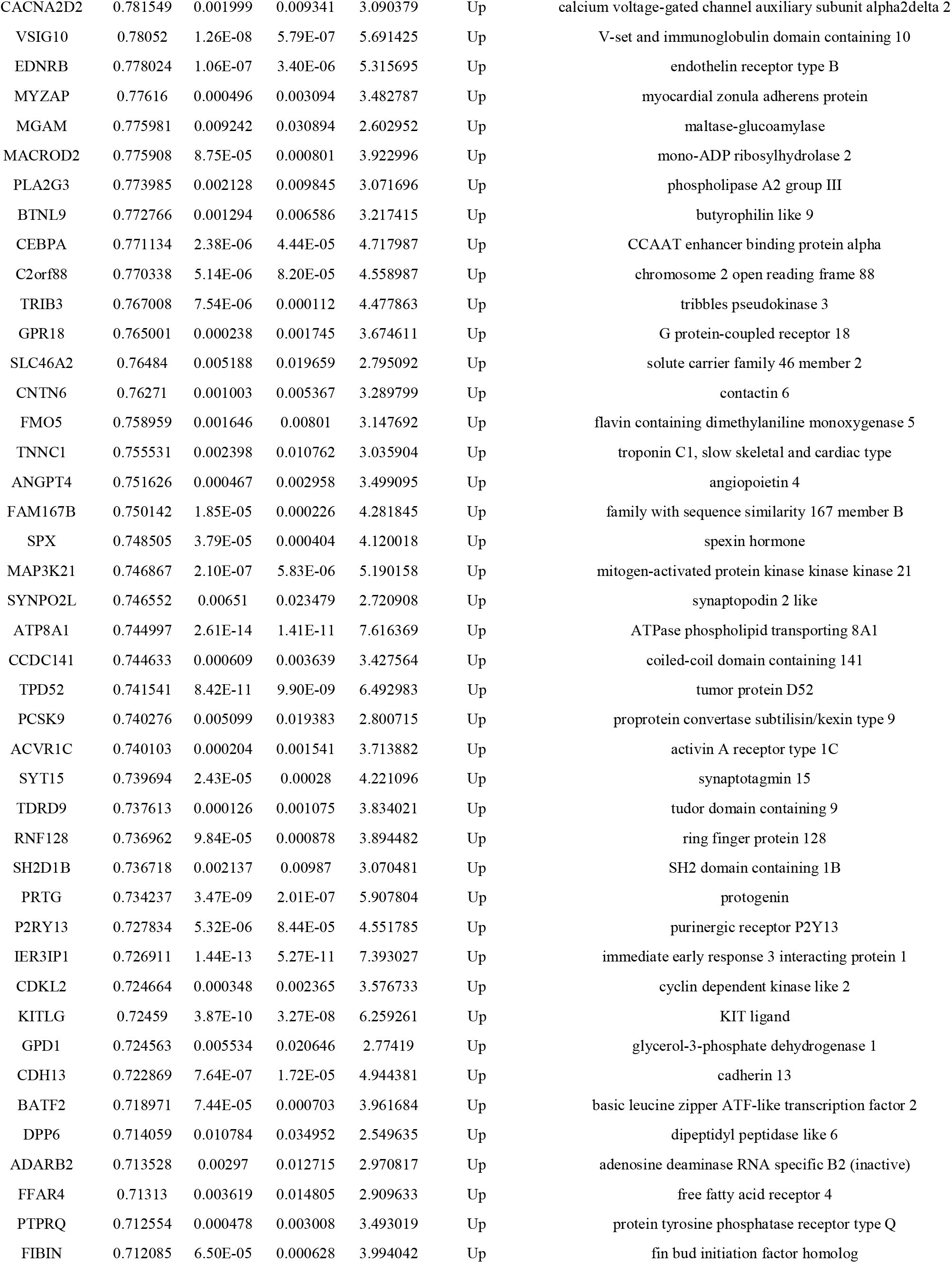
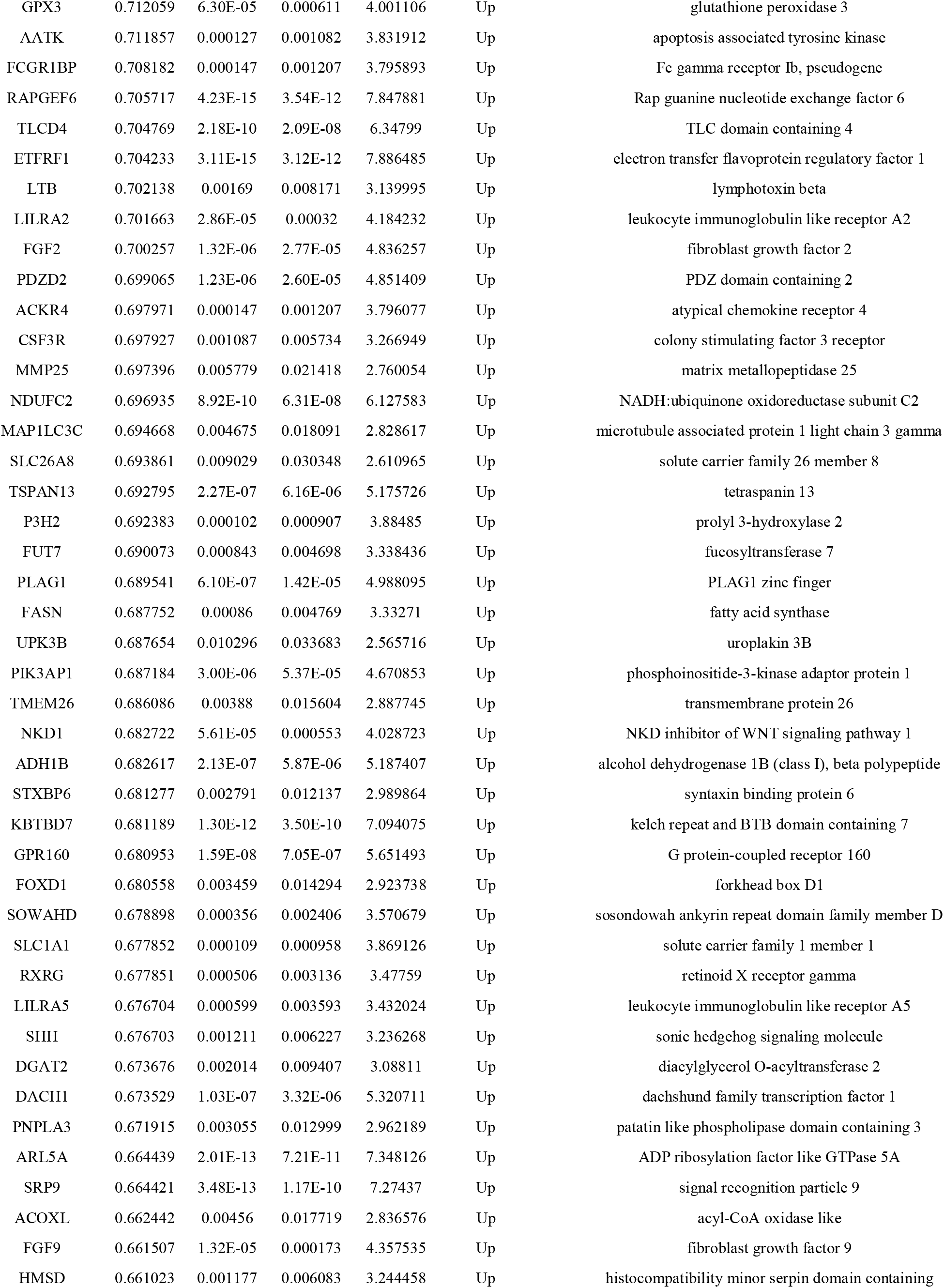
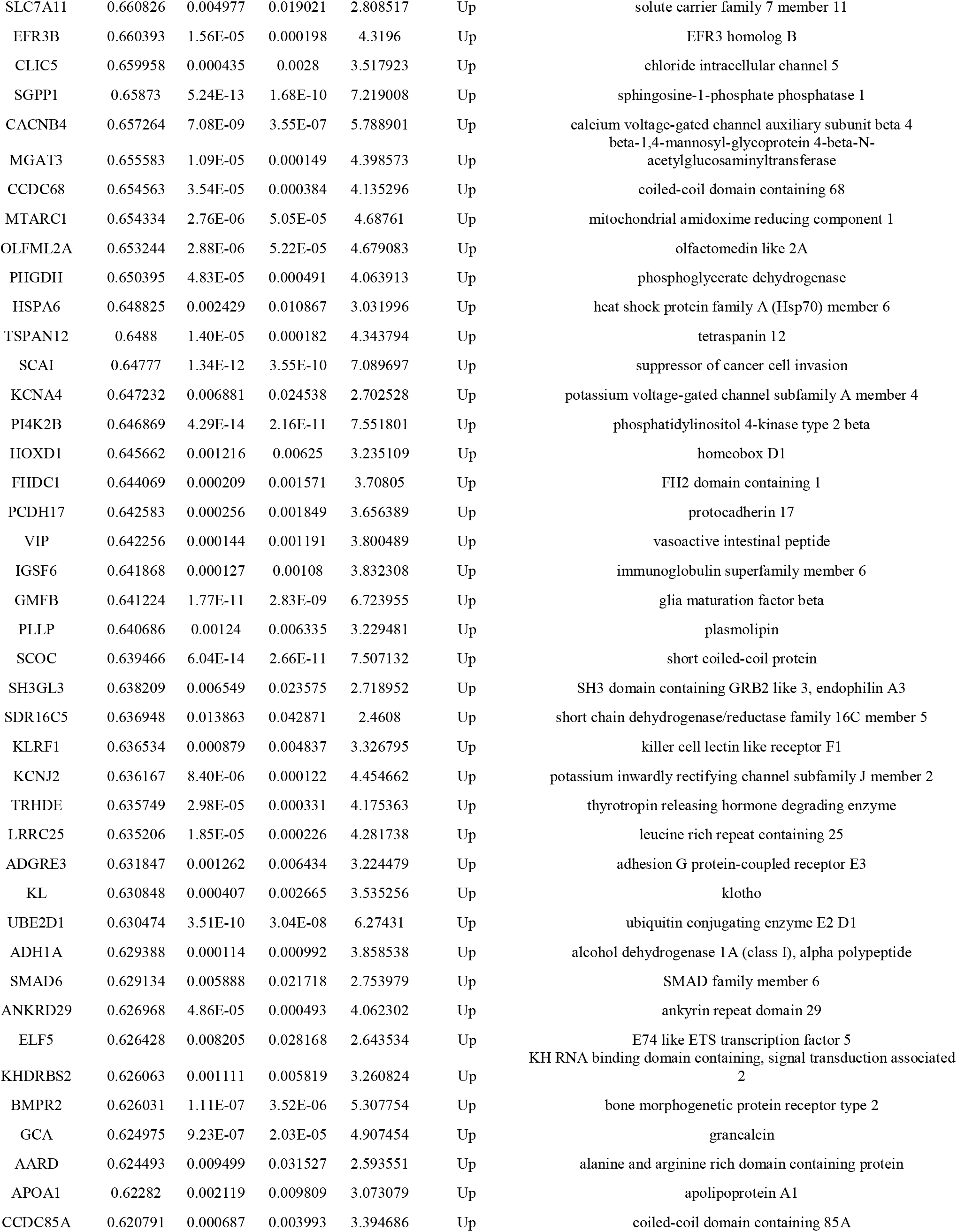
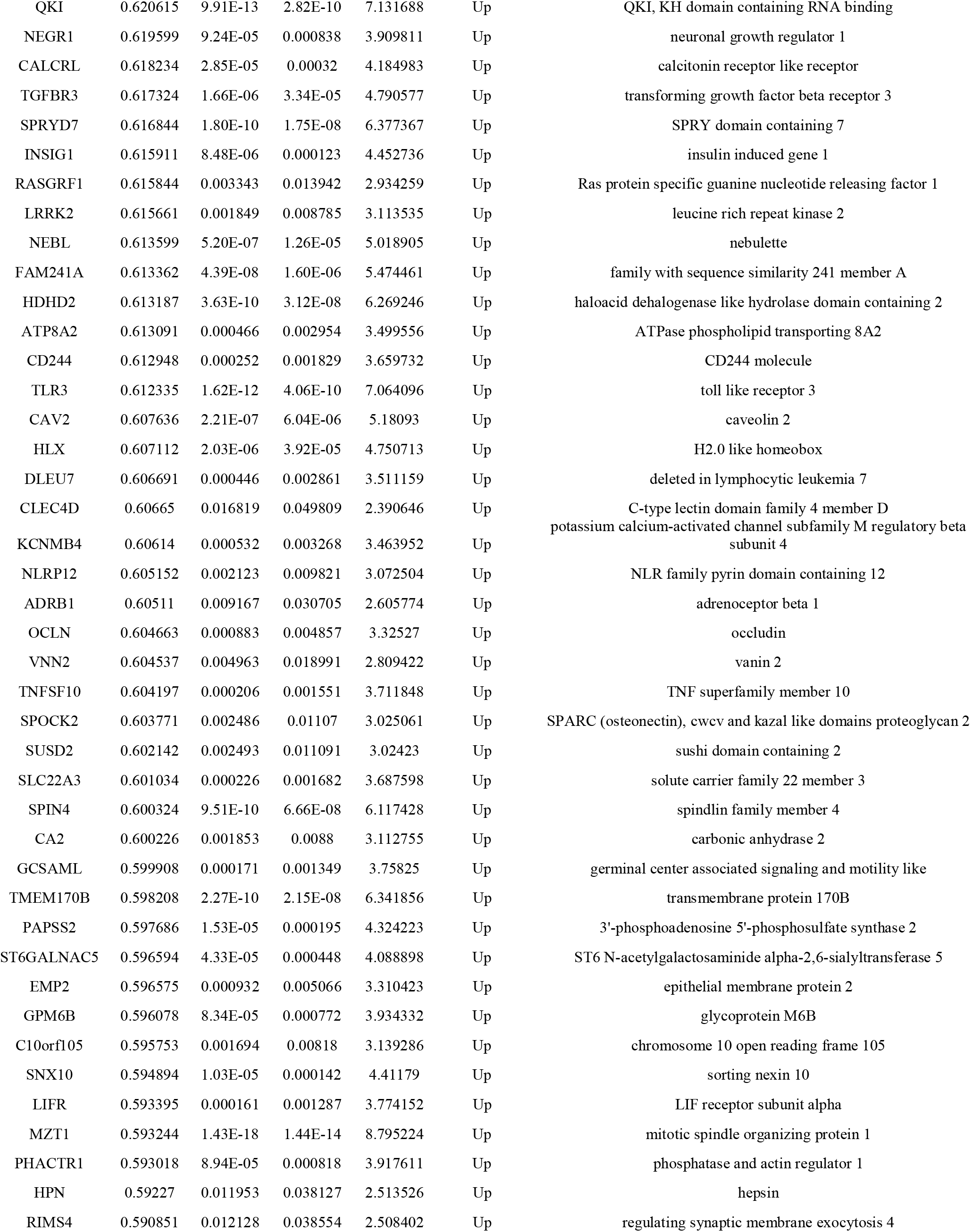
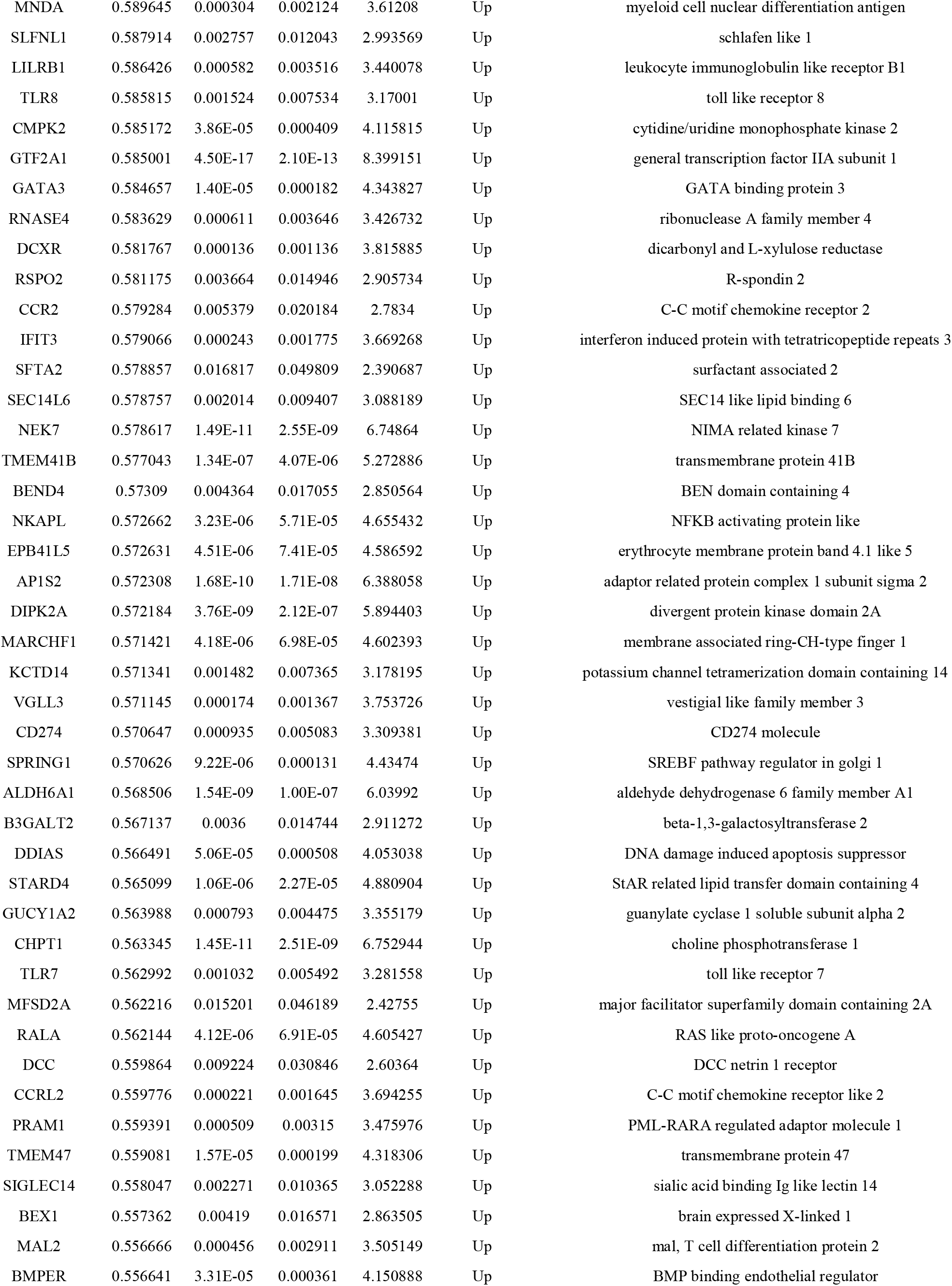
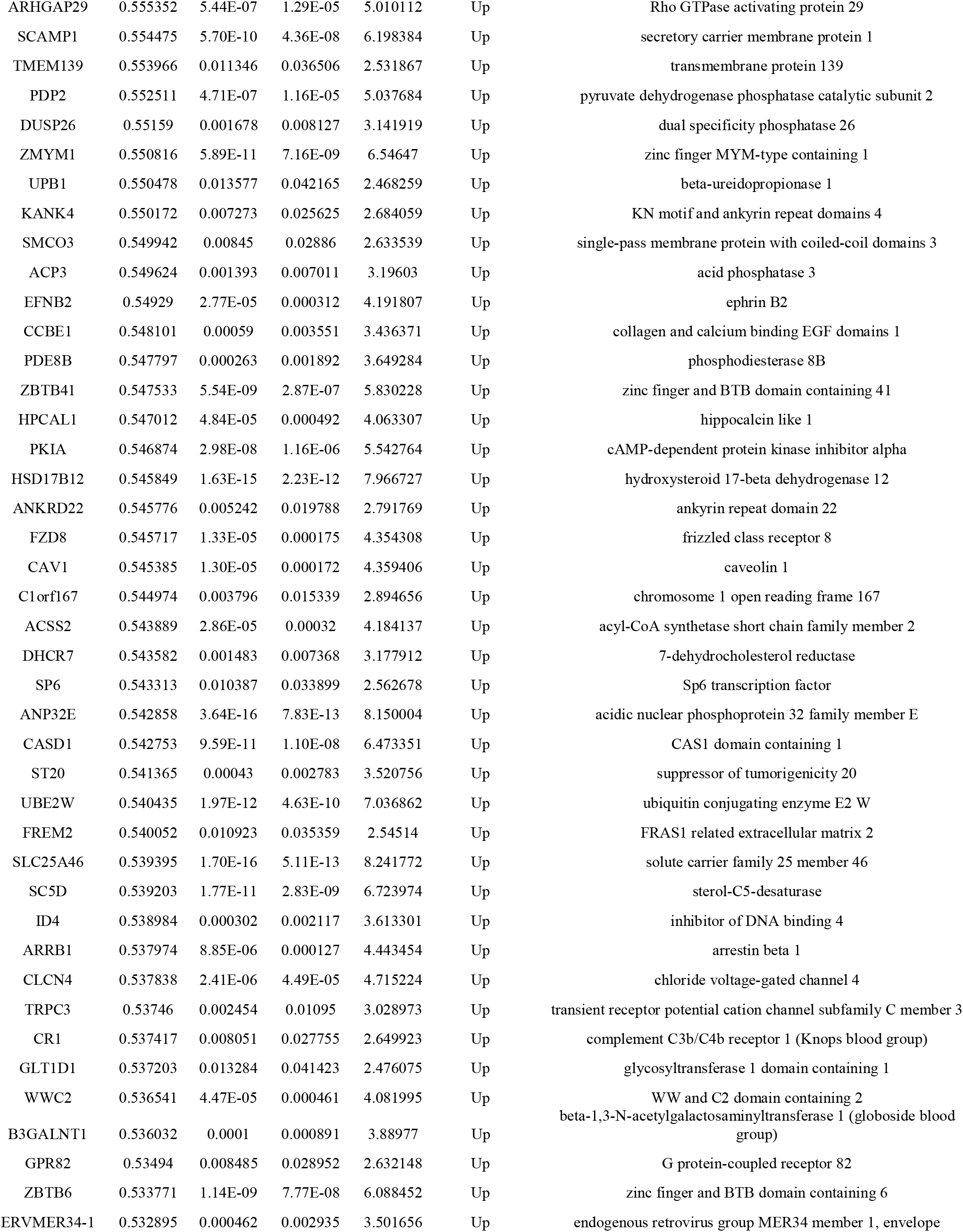
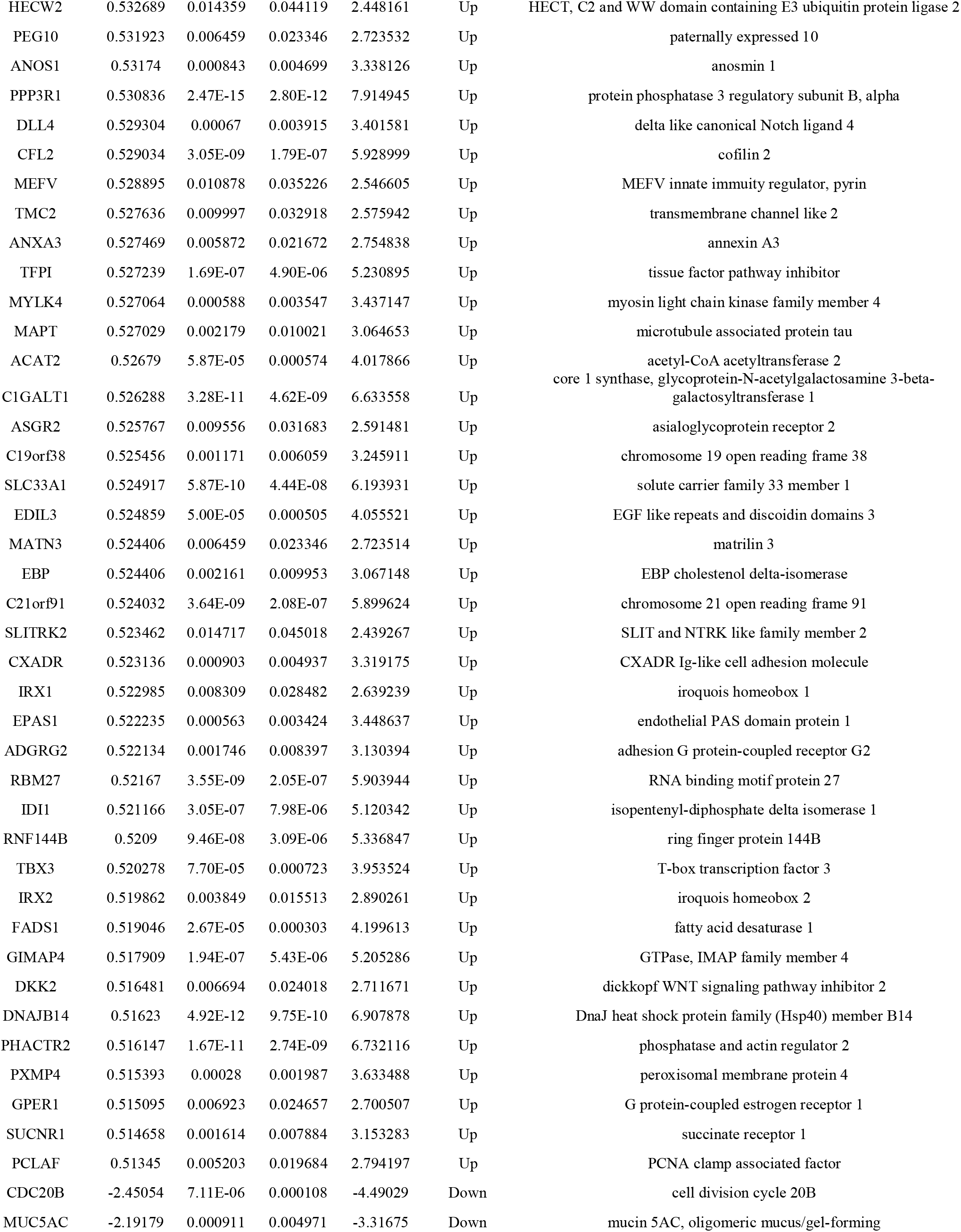
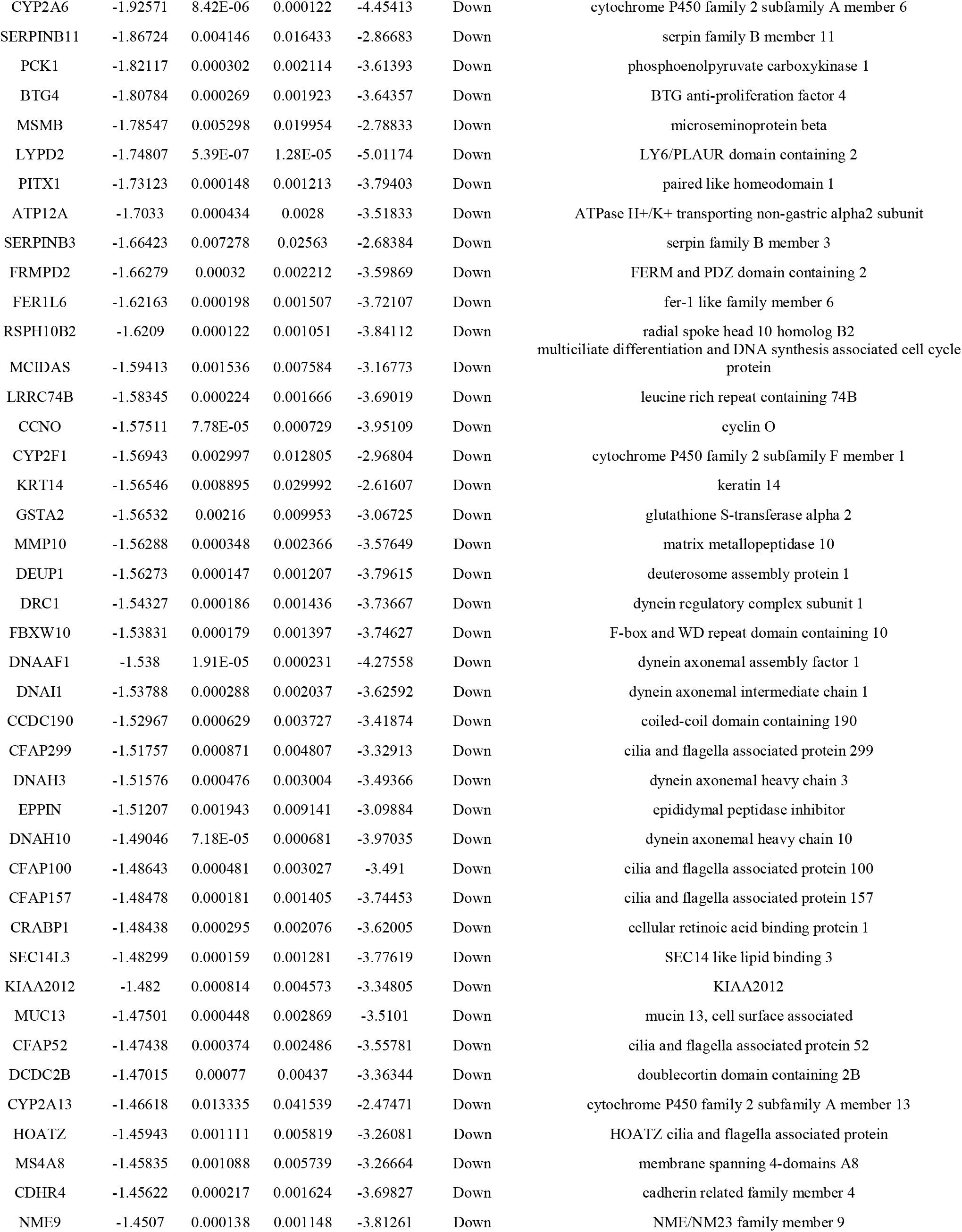
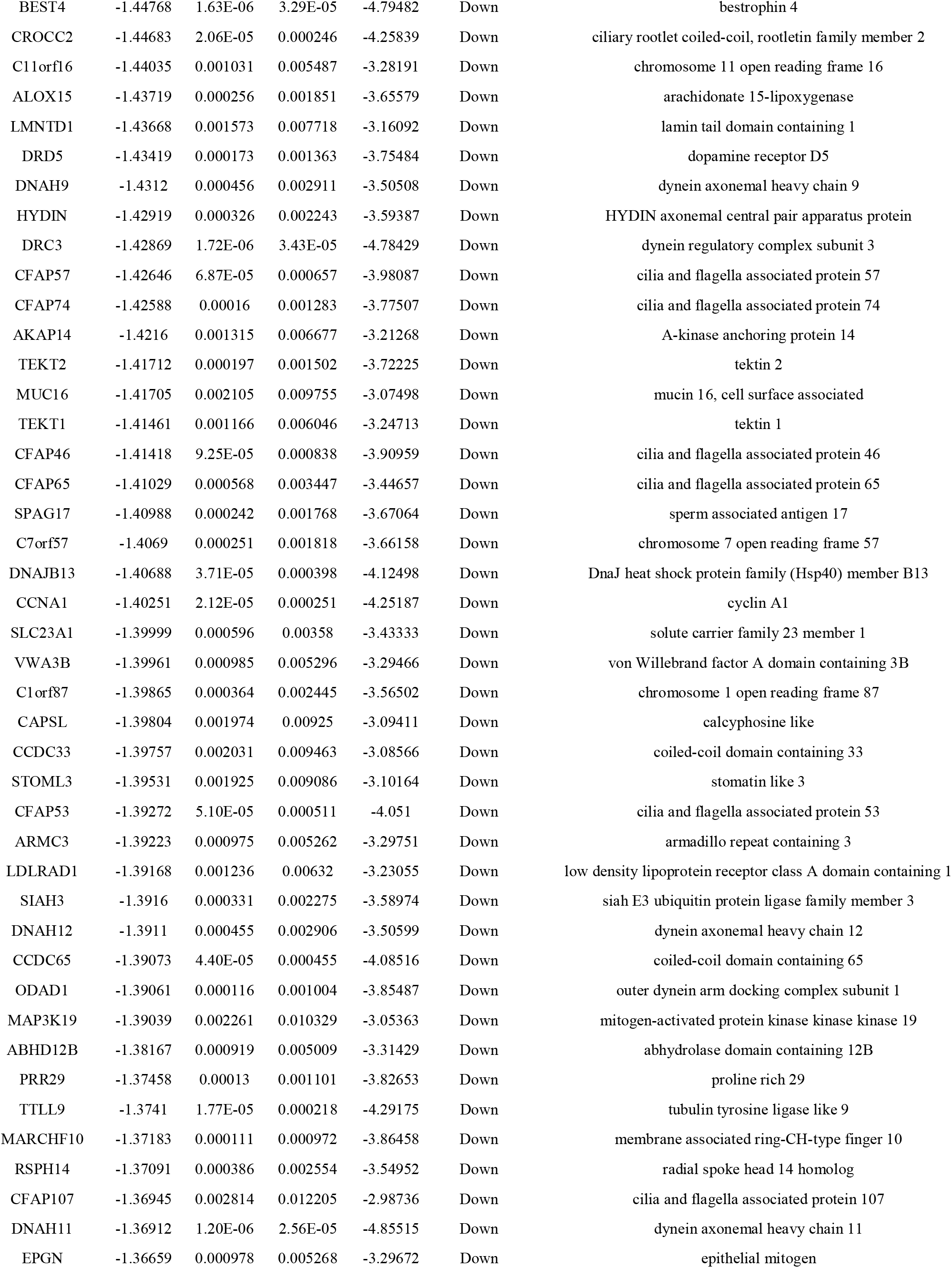
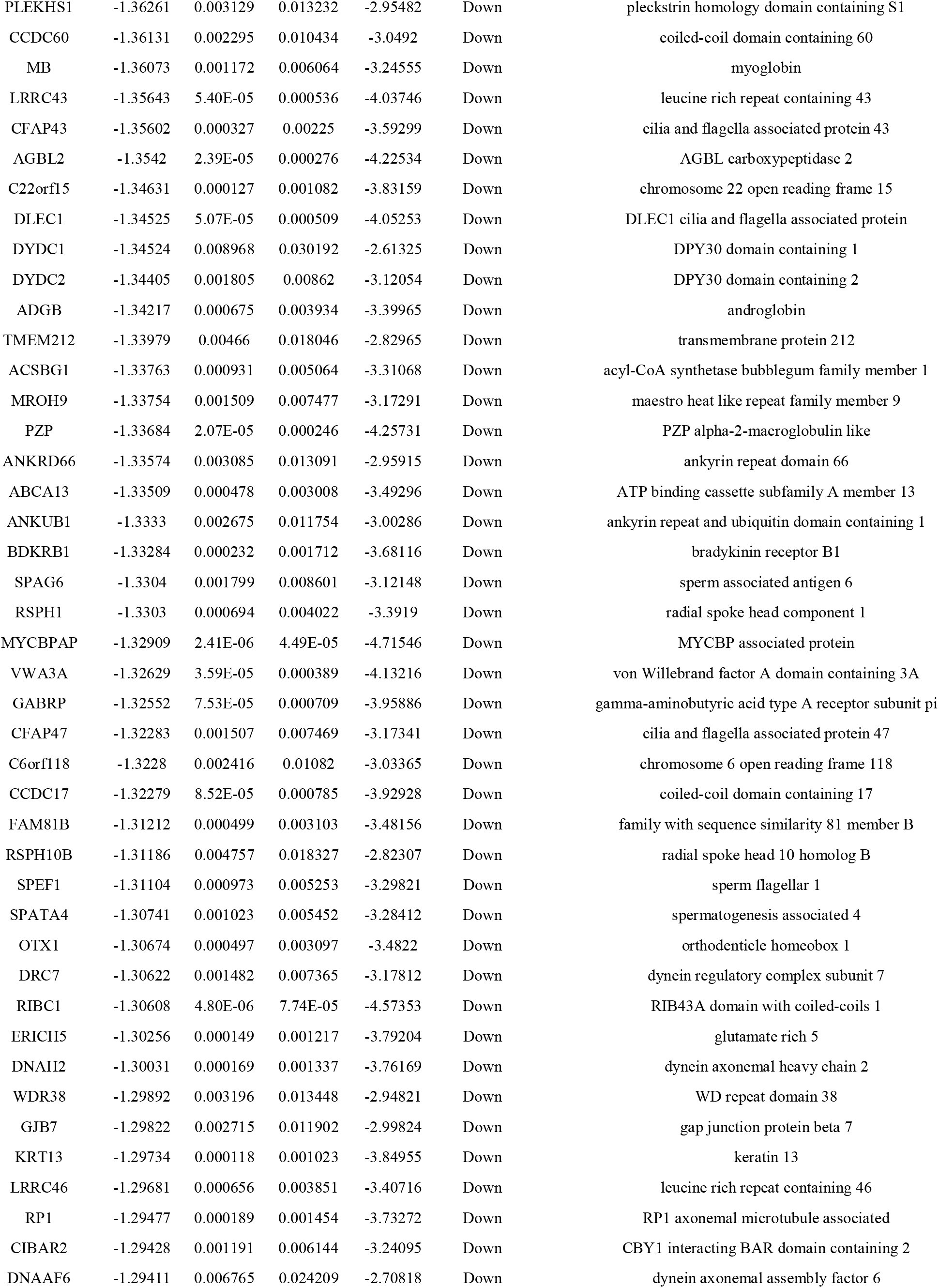
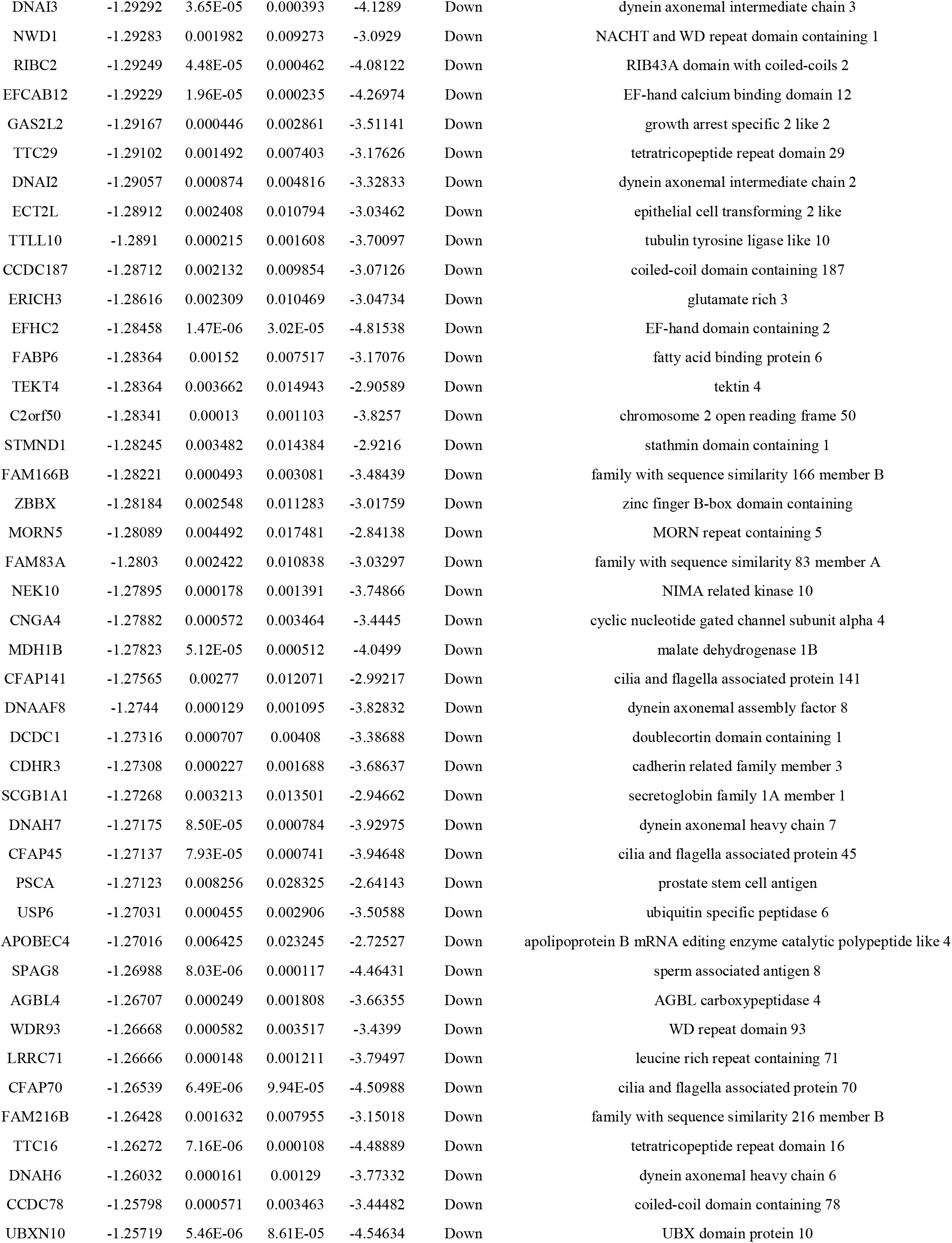
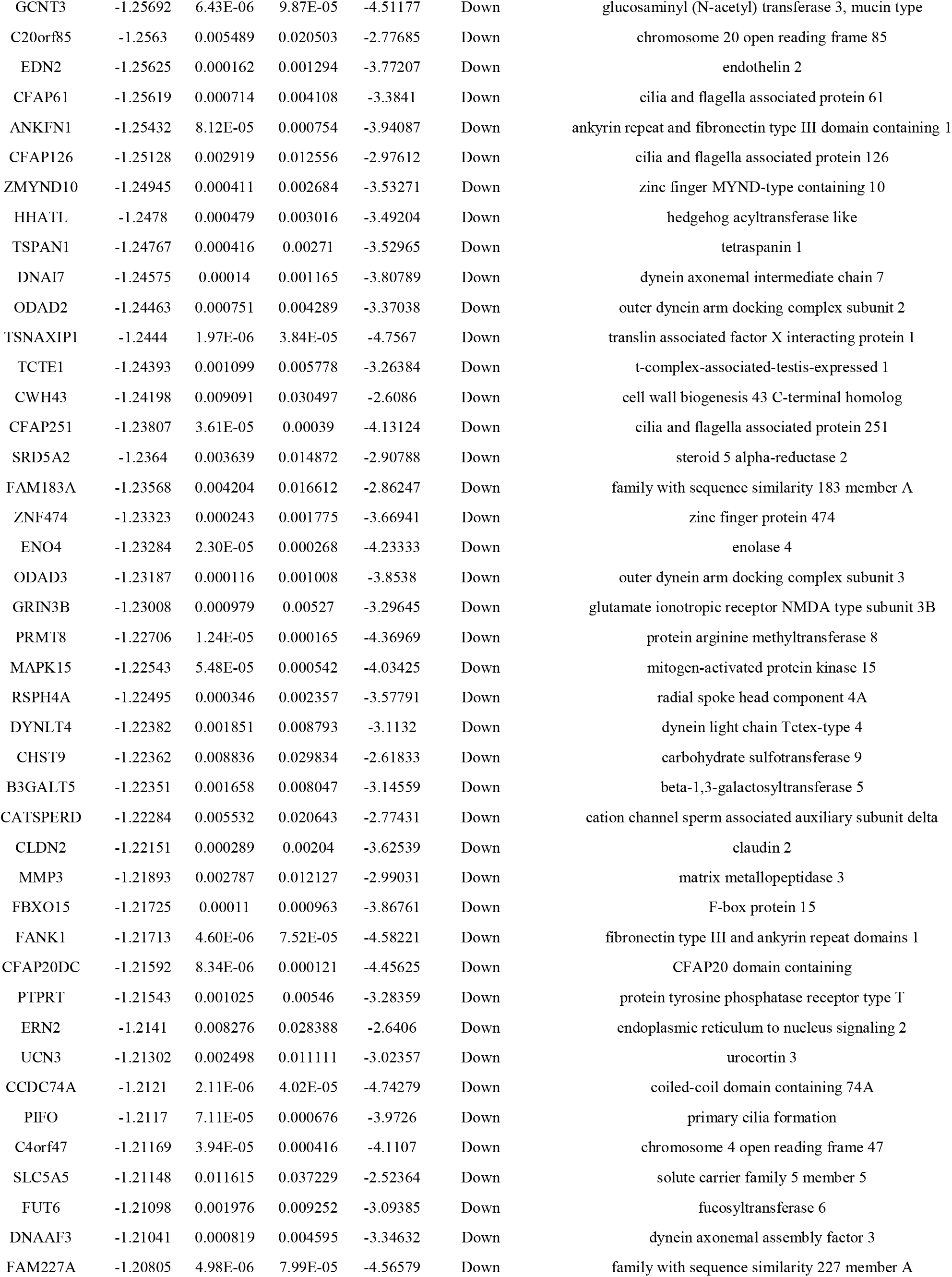
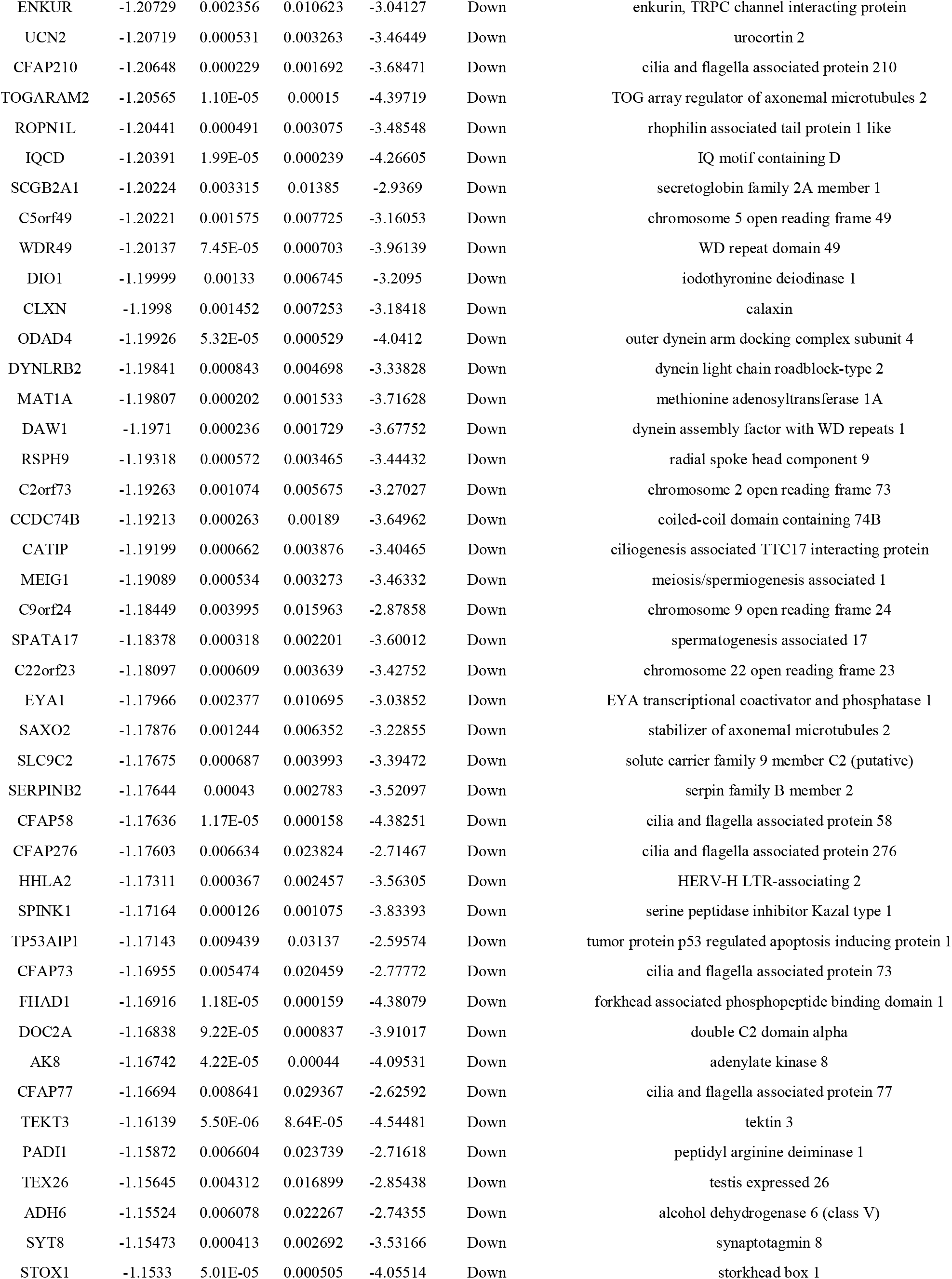
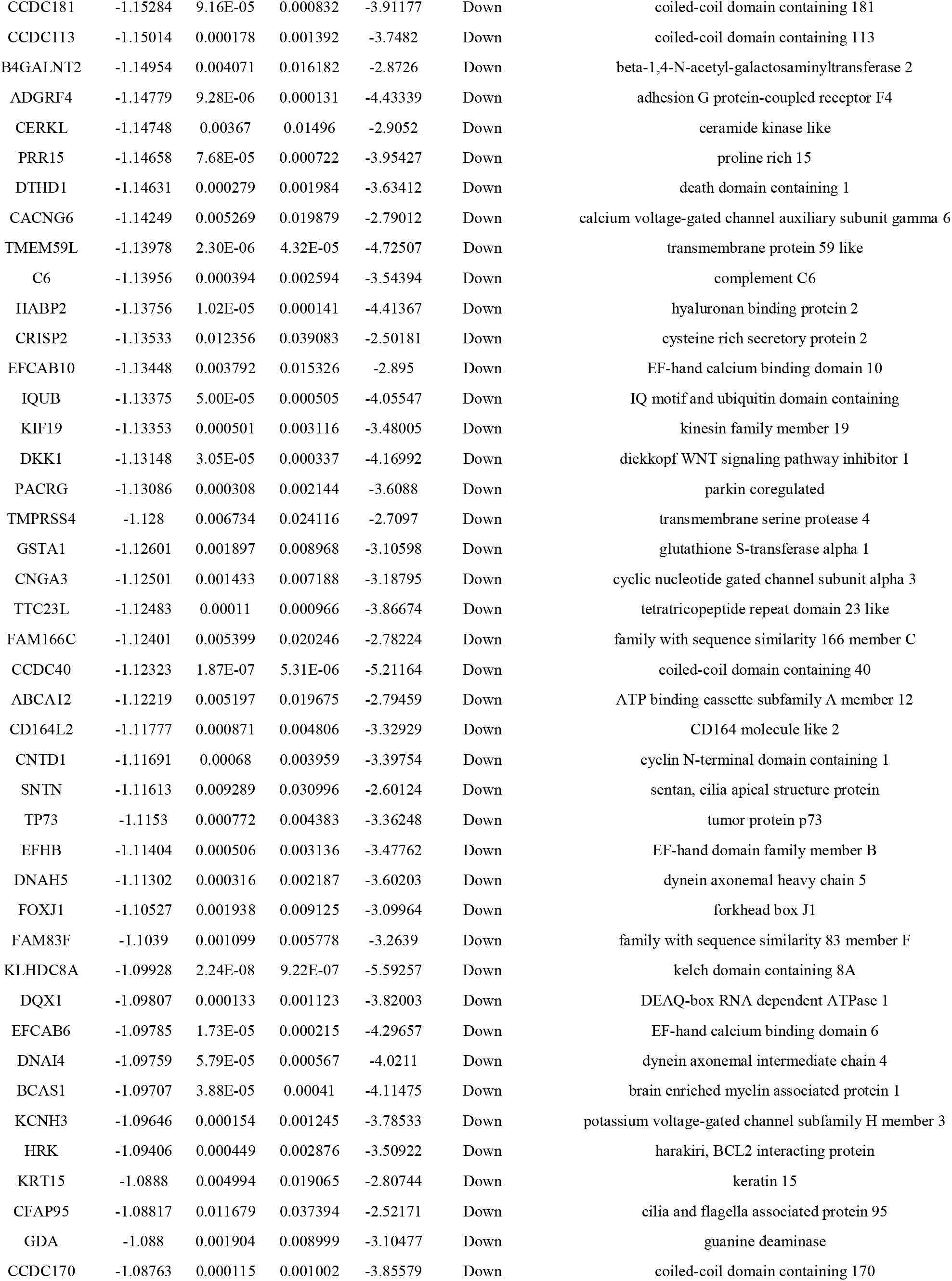
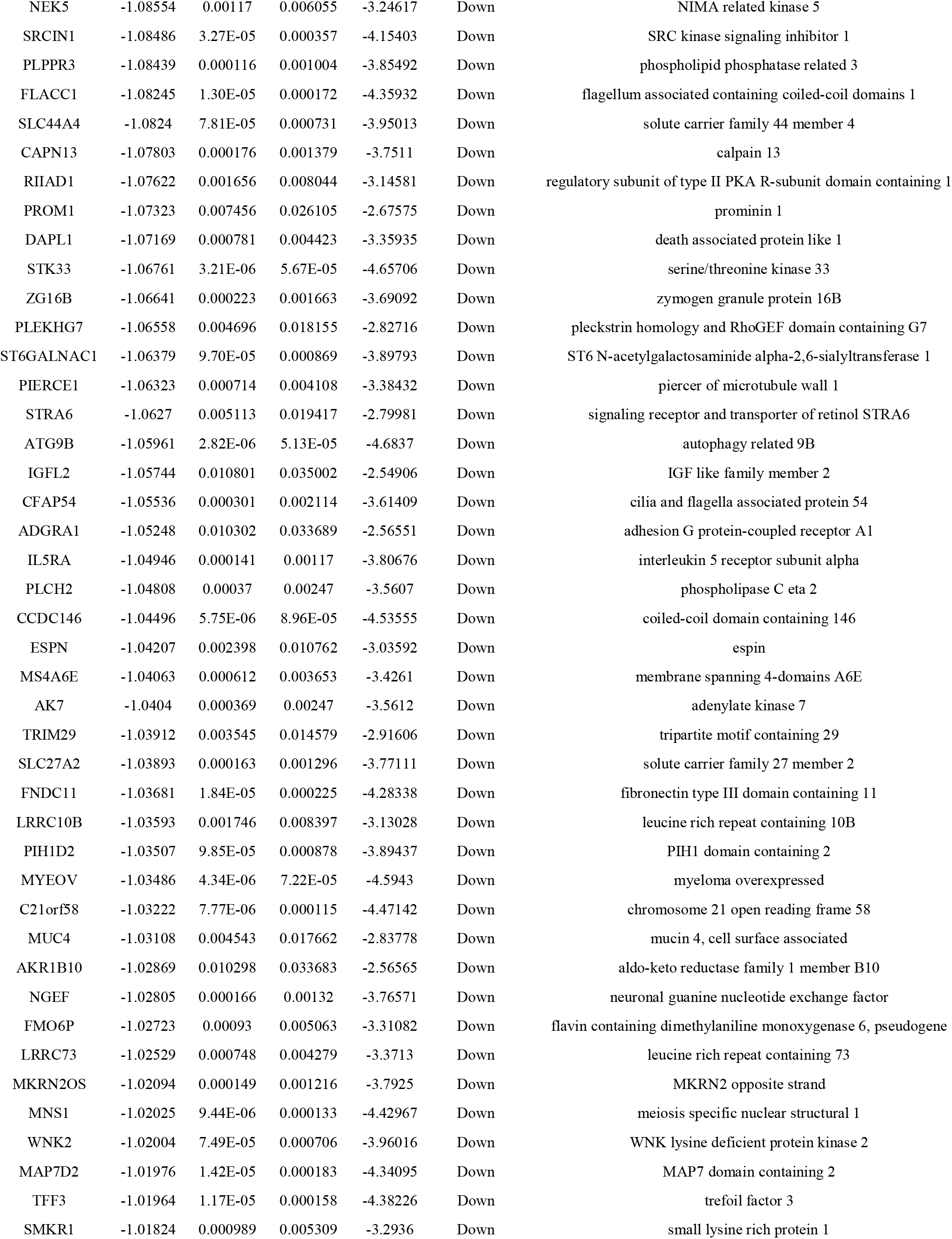
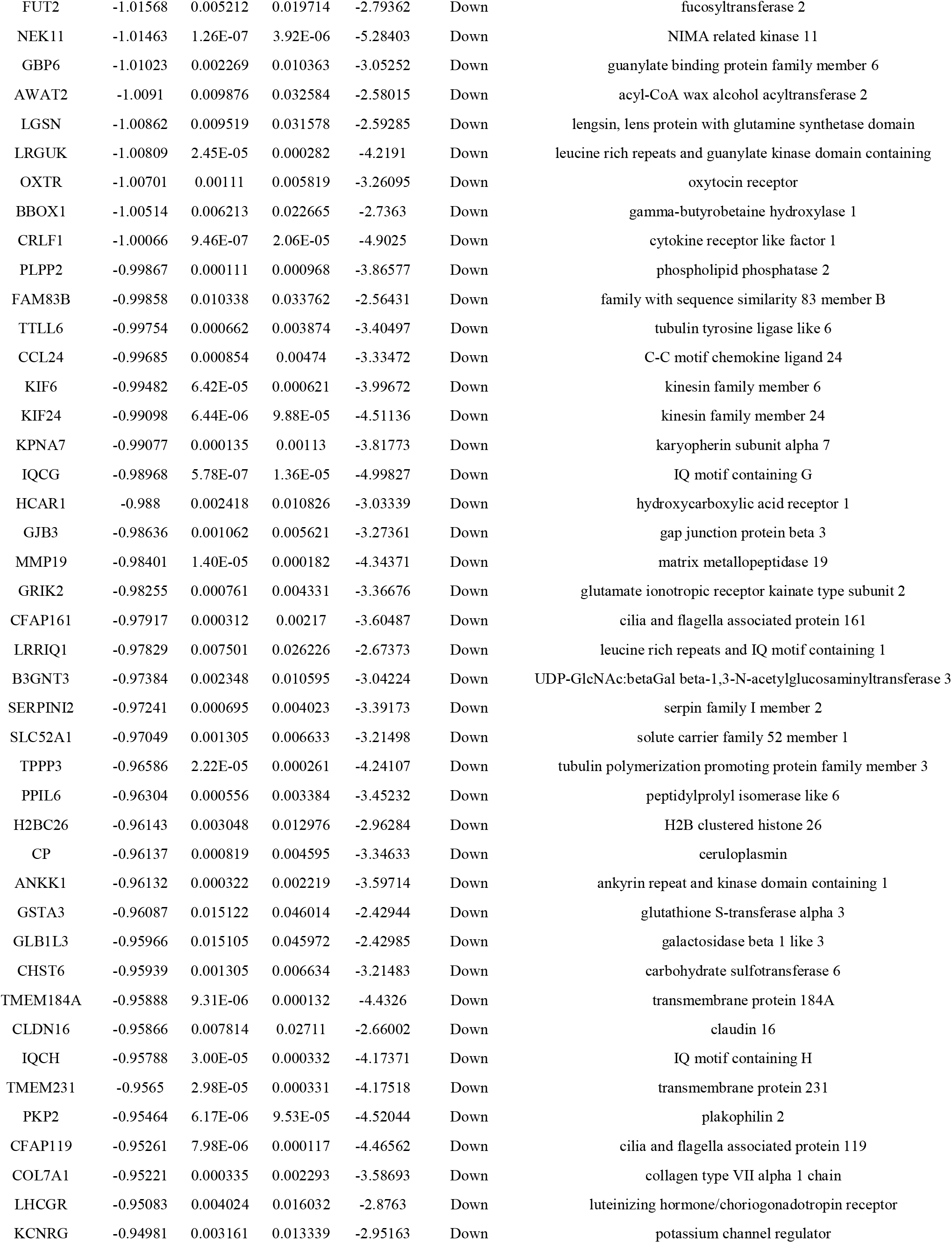
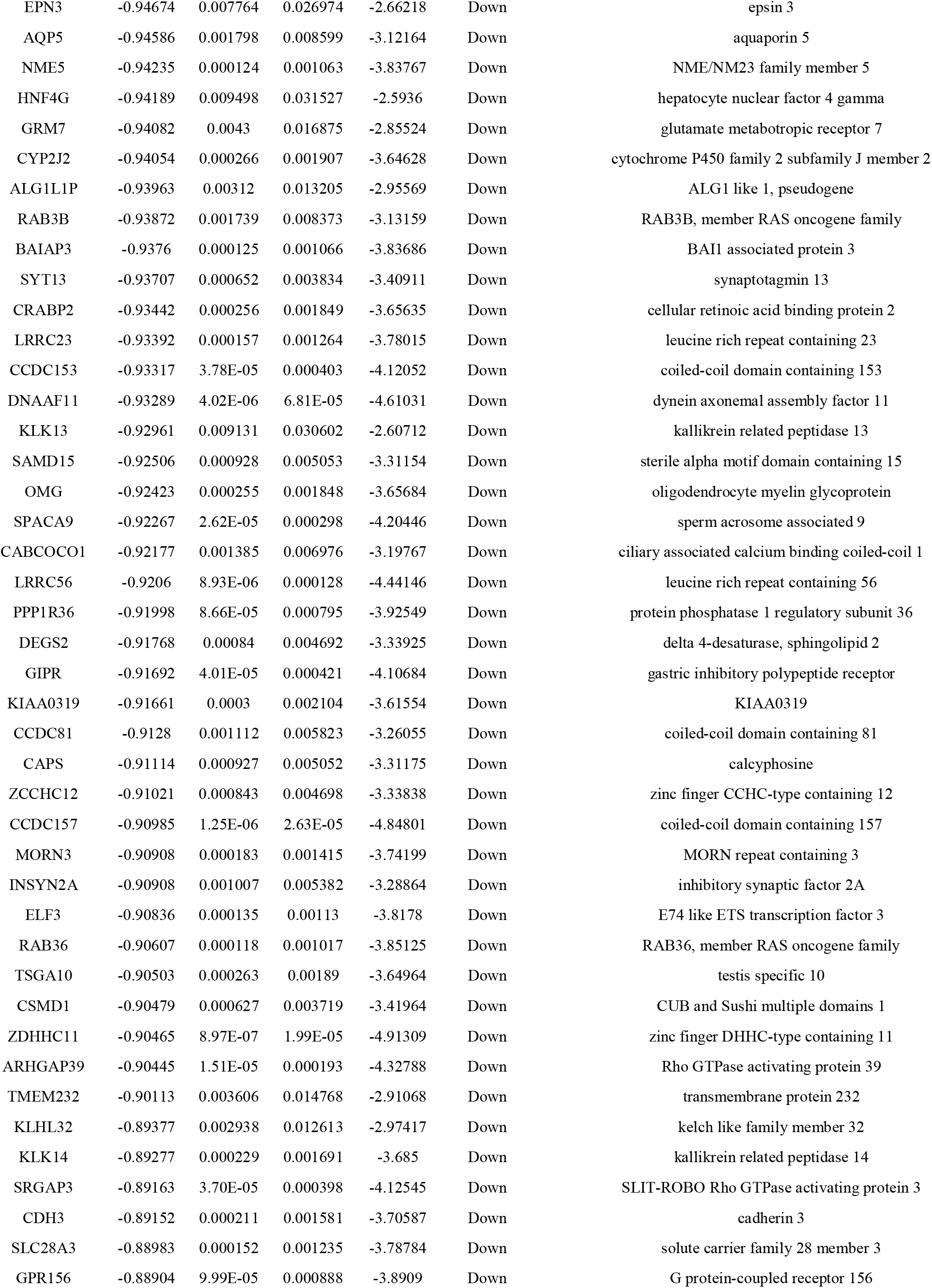
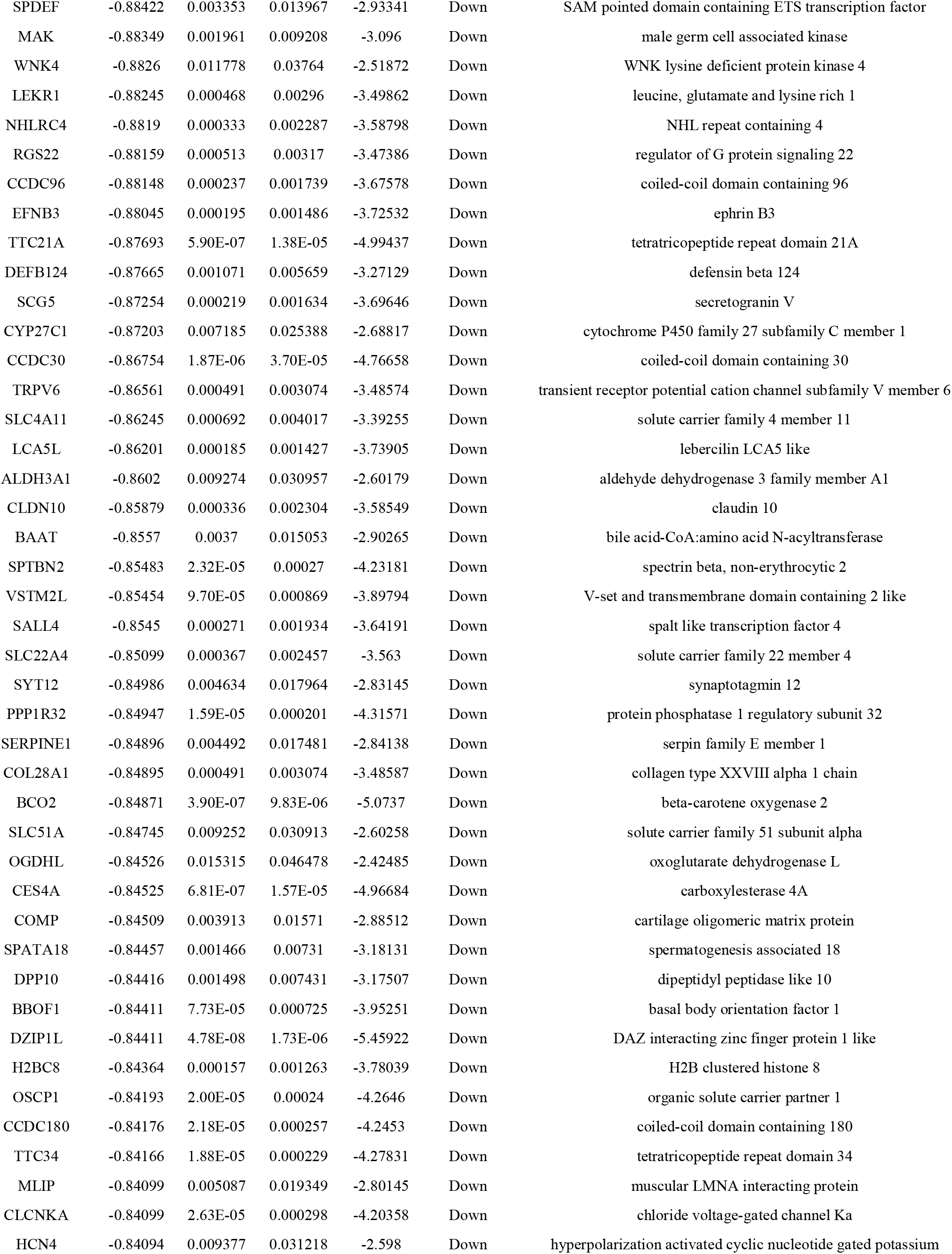
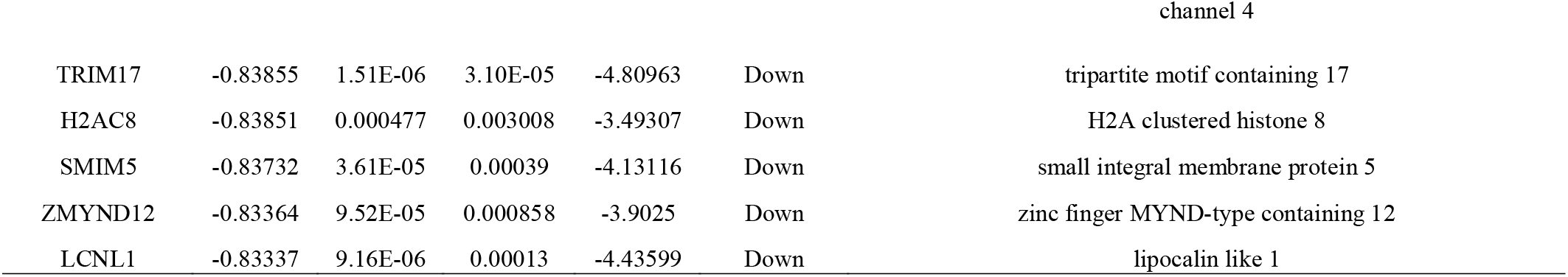
The statistical metrics for key differentially expressed genes (DEGs)

### GO and pathway enrichment analyses of DEGs

Functional enrichment analyses of the GO terms and REACTOME pathway were performed for both up regulated and down regulated DEGs. To gain insight into the BP, CC and MF of the DEGs products, we performed a GO enrichment analysis (Table 2). The GO analysis extracted from IPF patients and normal control subjects revealed that DEGs were significantly enriched in the following BP: response to stimulus, biological regulation, microtubule-based process and plasma membrane bounded cell projection organization. The CC analysis revealed that DEGs were predominantly located in the cell periphery, membrane, cell projection and cytoplasm. In the MF category, the DEGs were mainly enriched in signaling receptor binding, molecular transducer activity, tubulin binding and calcium ion binding. Meanwhile, REACTOME enrichment analysis revealed that pathways were significantly enriched including GPCR ligand binding, class A/1 (Rhodopsin-like receptors), defective GALNT3 causes HFTC and Lewis blood group biosynthesis (Table 3).

**Table 2.**
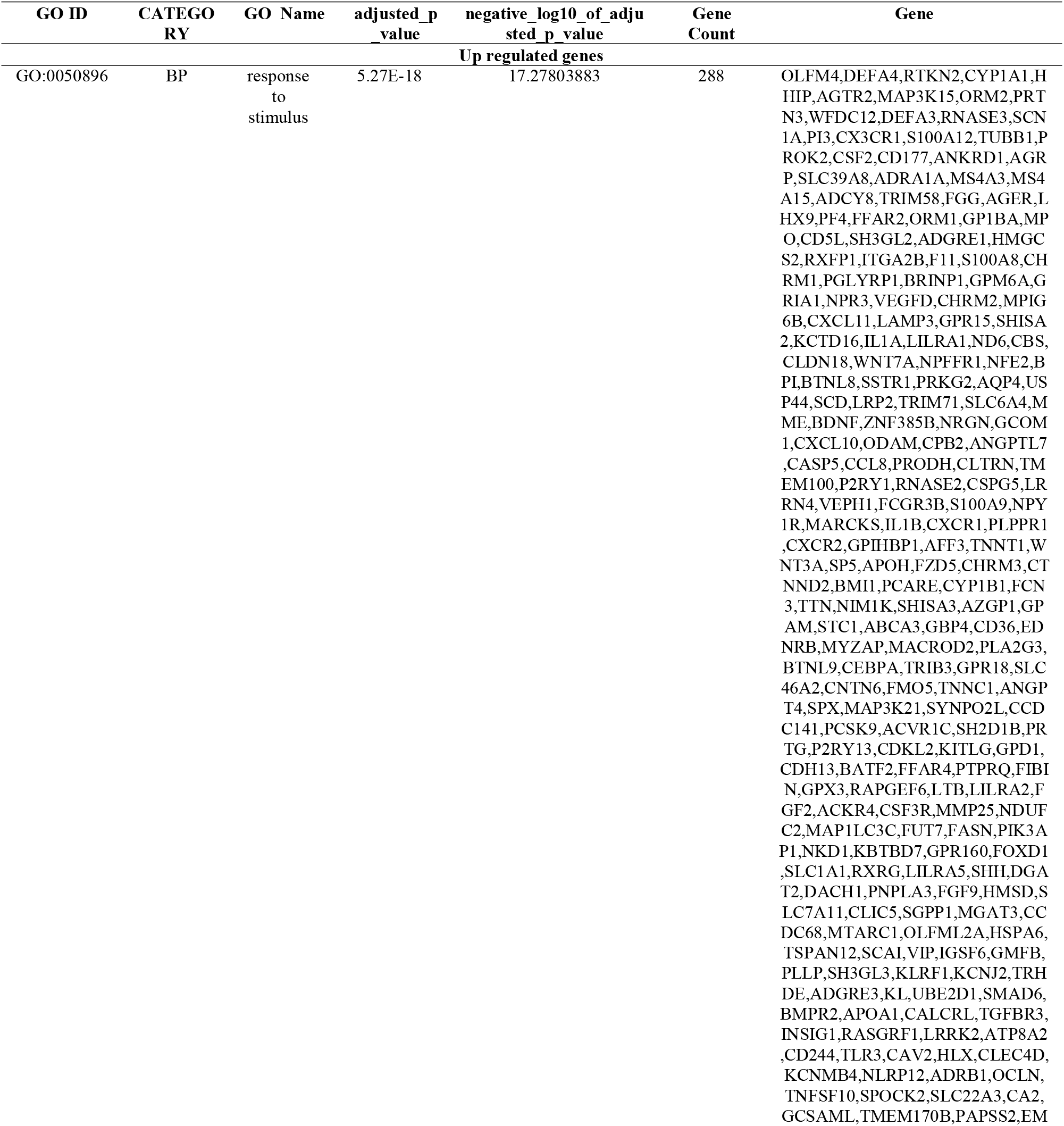
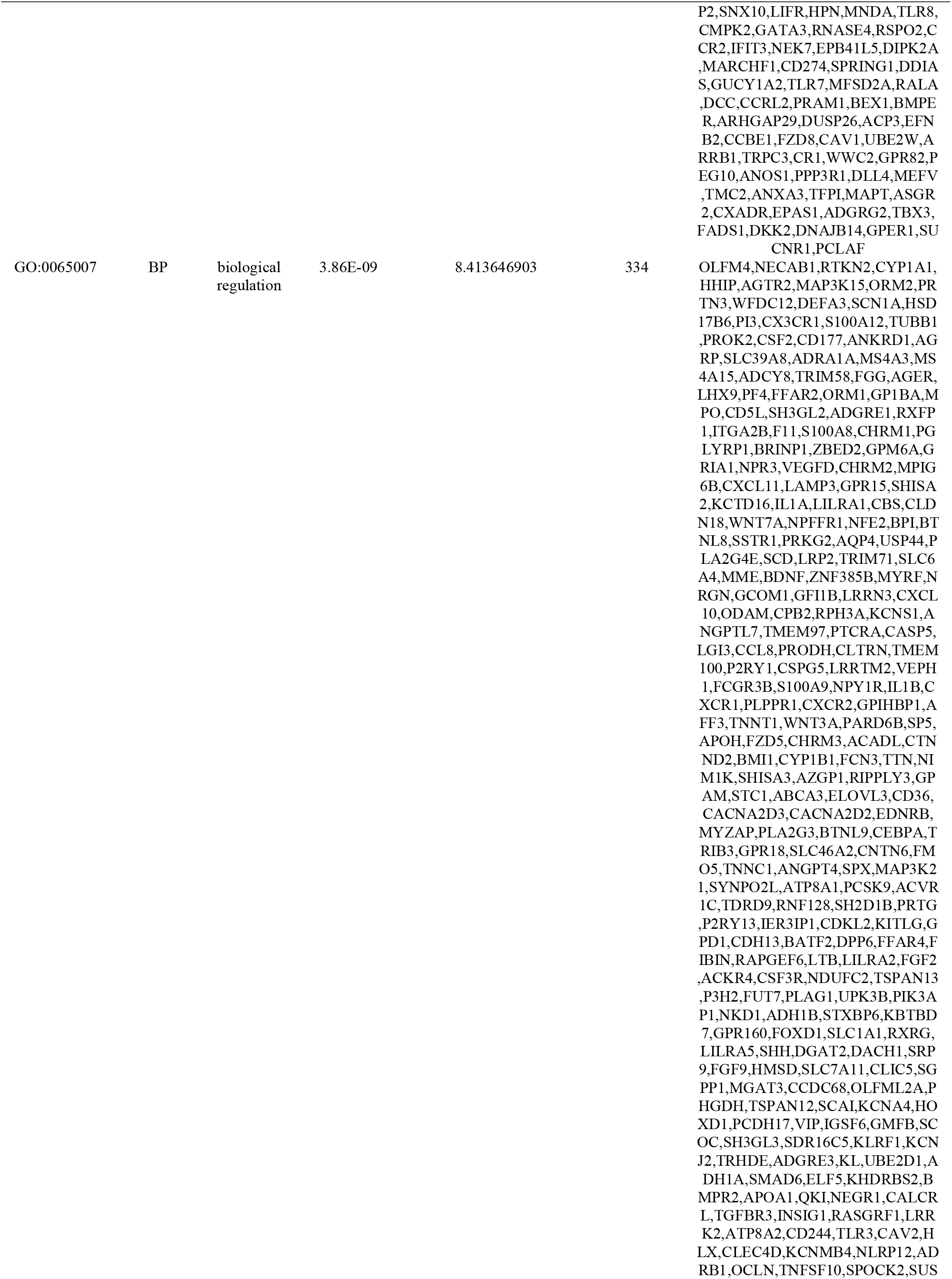
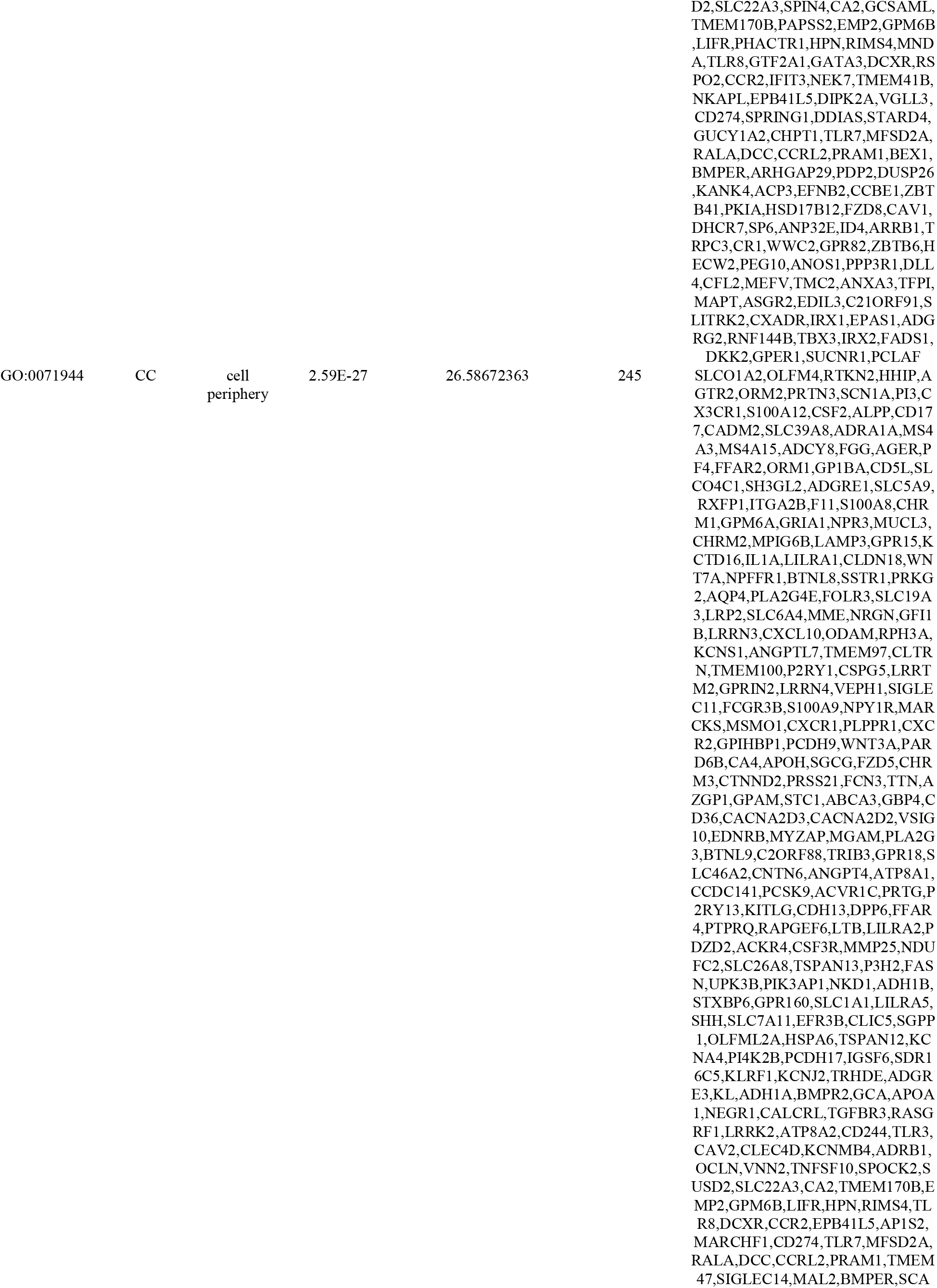
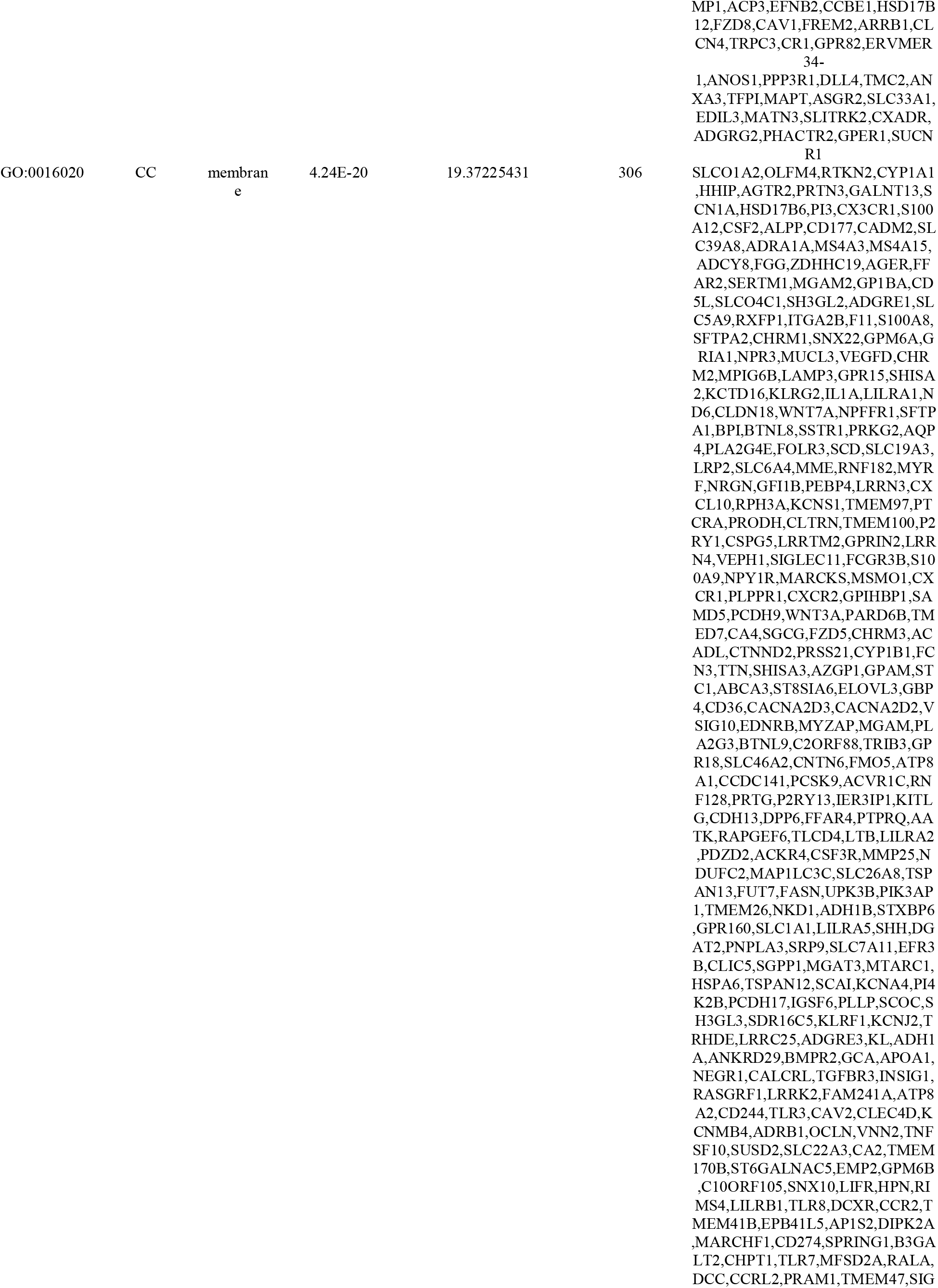
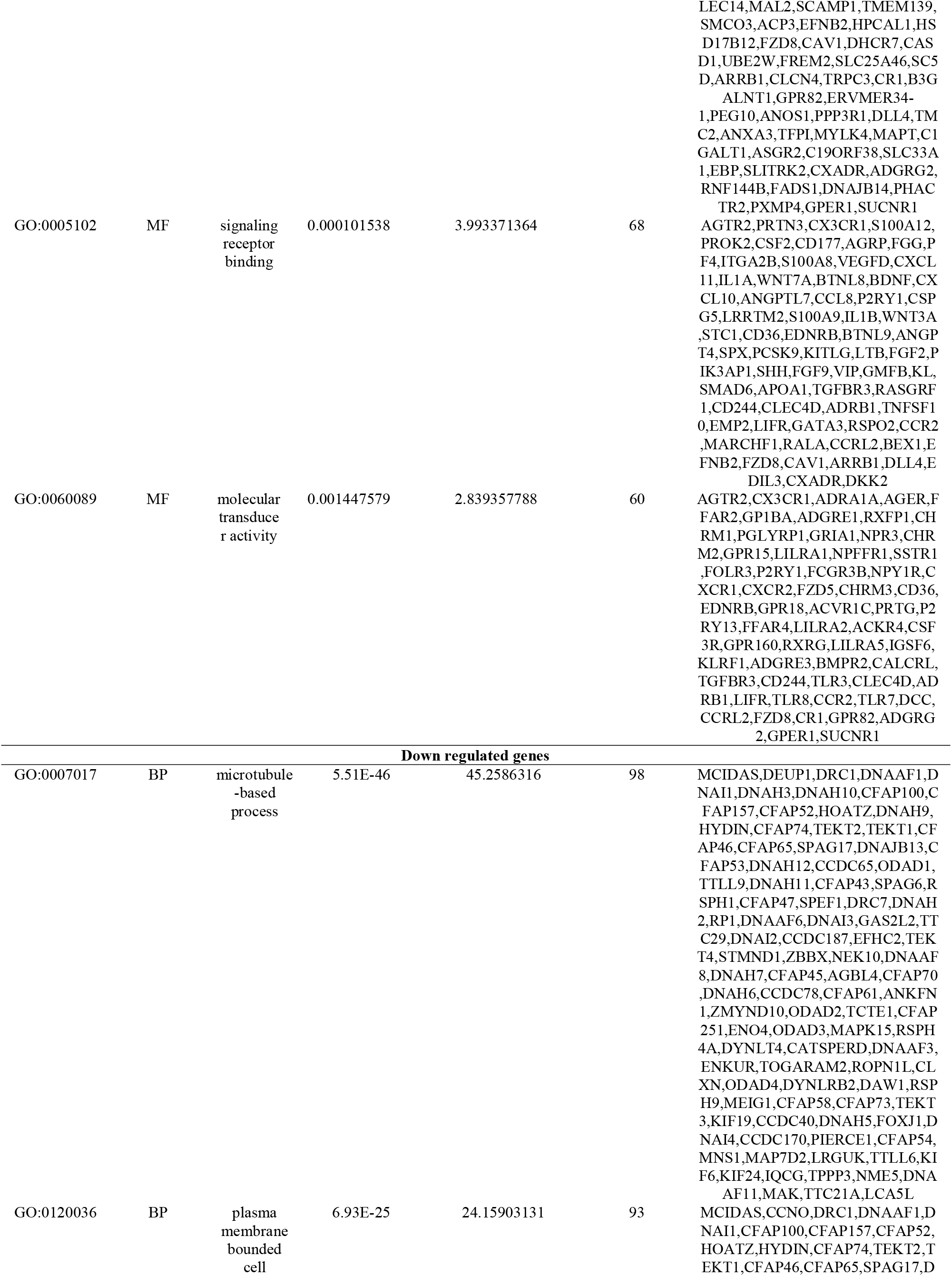
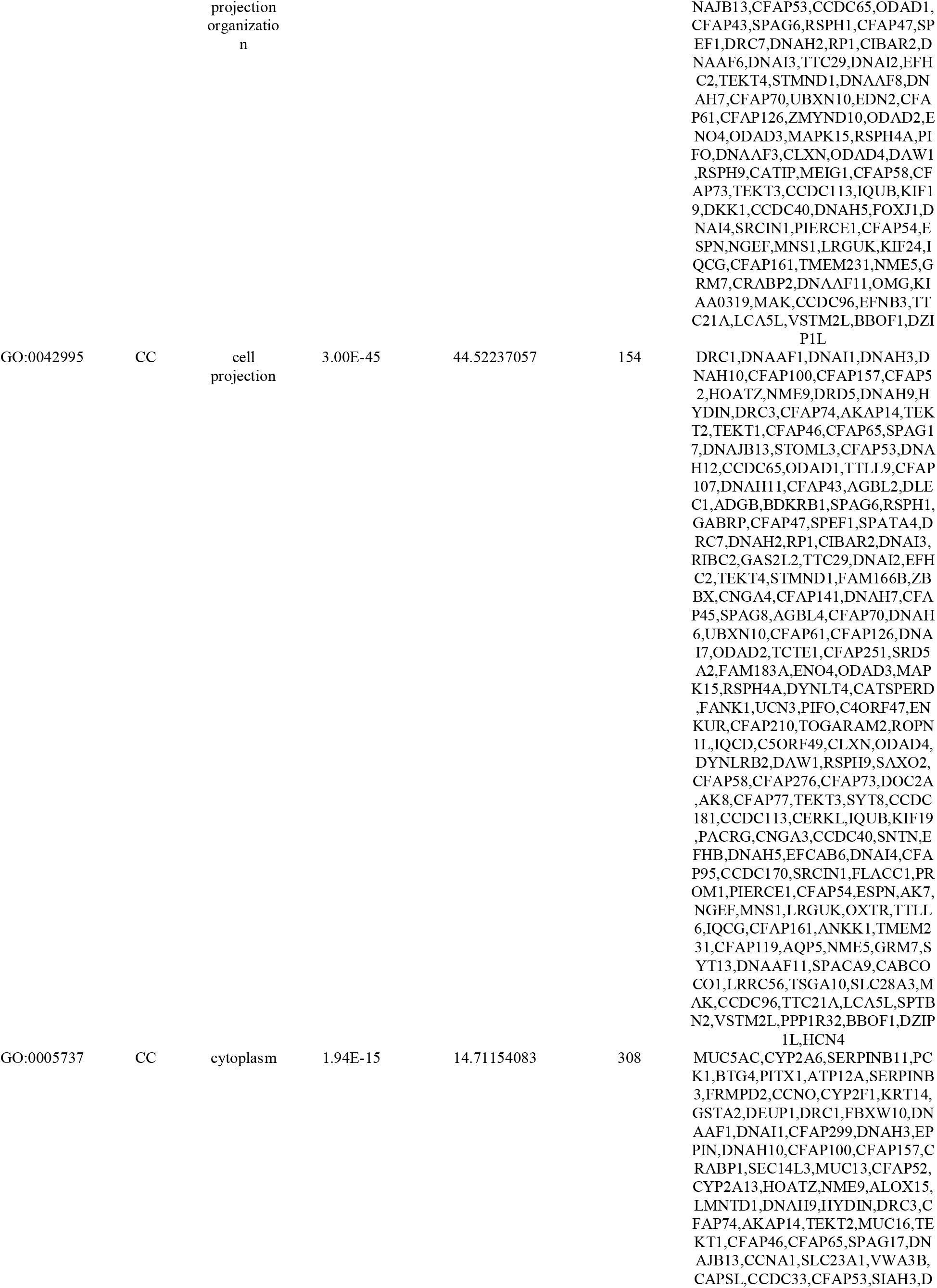
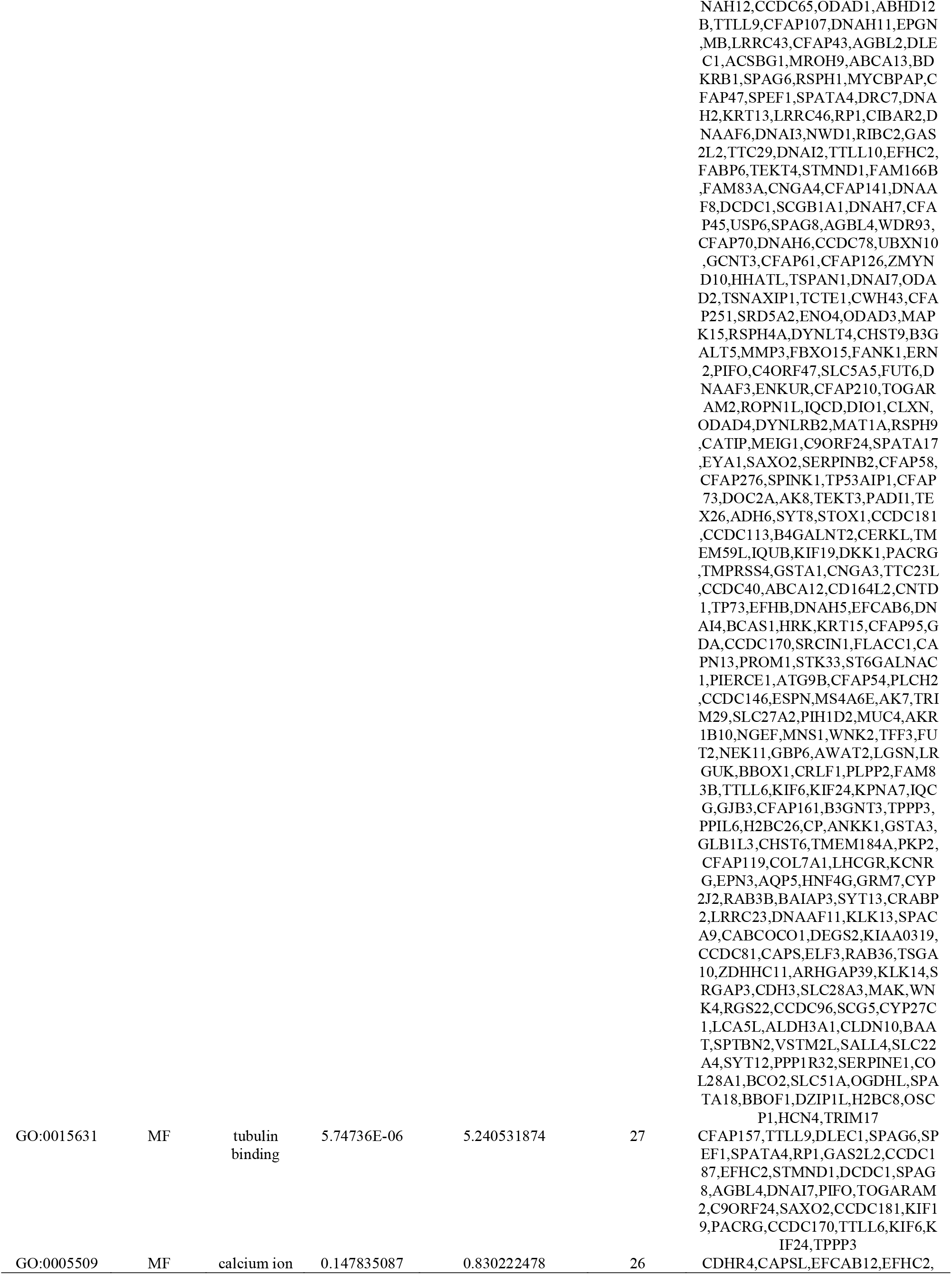
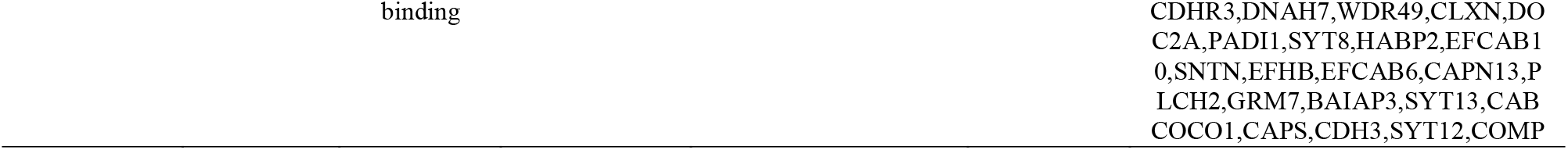
The enriched GO terms of the up and down regulated differentially expressed genes.

**Table 3.**
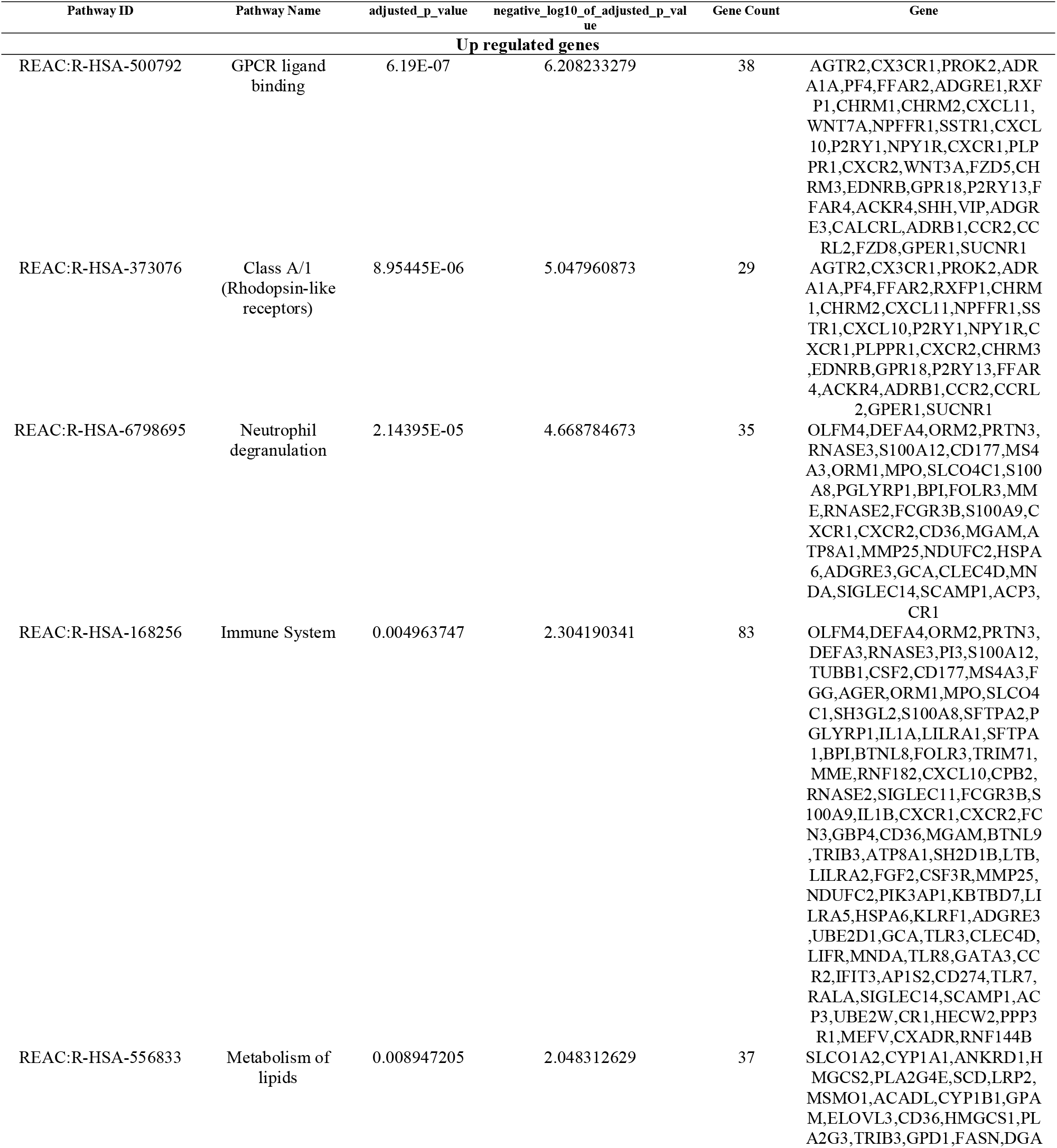
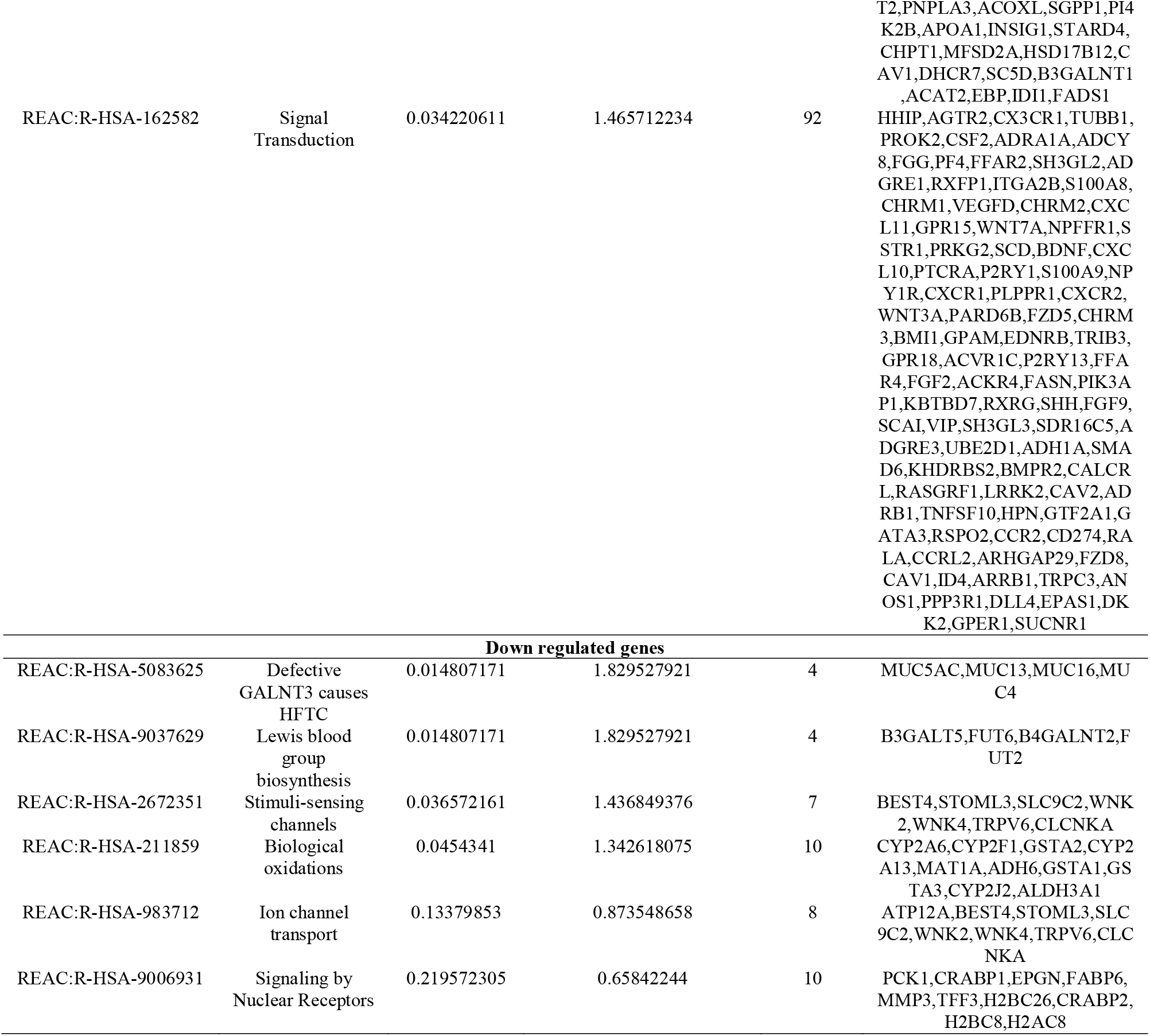
The enriched pathway terms of the up and down regulated differentially expressed genes.

### Construction of the PPI network and module analysis

The IID database was used to identify the PPI pairs. As revealed in Fig. 3, 5557 nodes (DEGs) and 9632 edges (interactions) were established in the constructed PPI network. According to the degree, betweenness, stress and closeness value, the top hub DEGs were determined. Ten hub genes (LRRK2, BMI1, EBP, MNDA, KBTBD7, KRT15, OTX1, TEKT4, SPAG8 and EFHC2) were identified through Cytoscape (Table 4). Module 1 and module 2 were significant modules in the PPI network. A total of 14 nodes and 26 edges were included in Module 1 (Fig. 4A), mainly involved in response to stimulus, metabolism of lipids and biological regulation, and a total of 9 nodes and 17 edges were included in Module 2 (Fig. 4B), mainly involved in cytoplasm and cell projection.

**Fig. 3.**
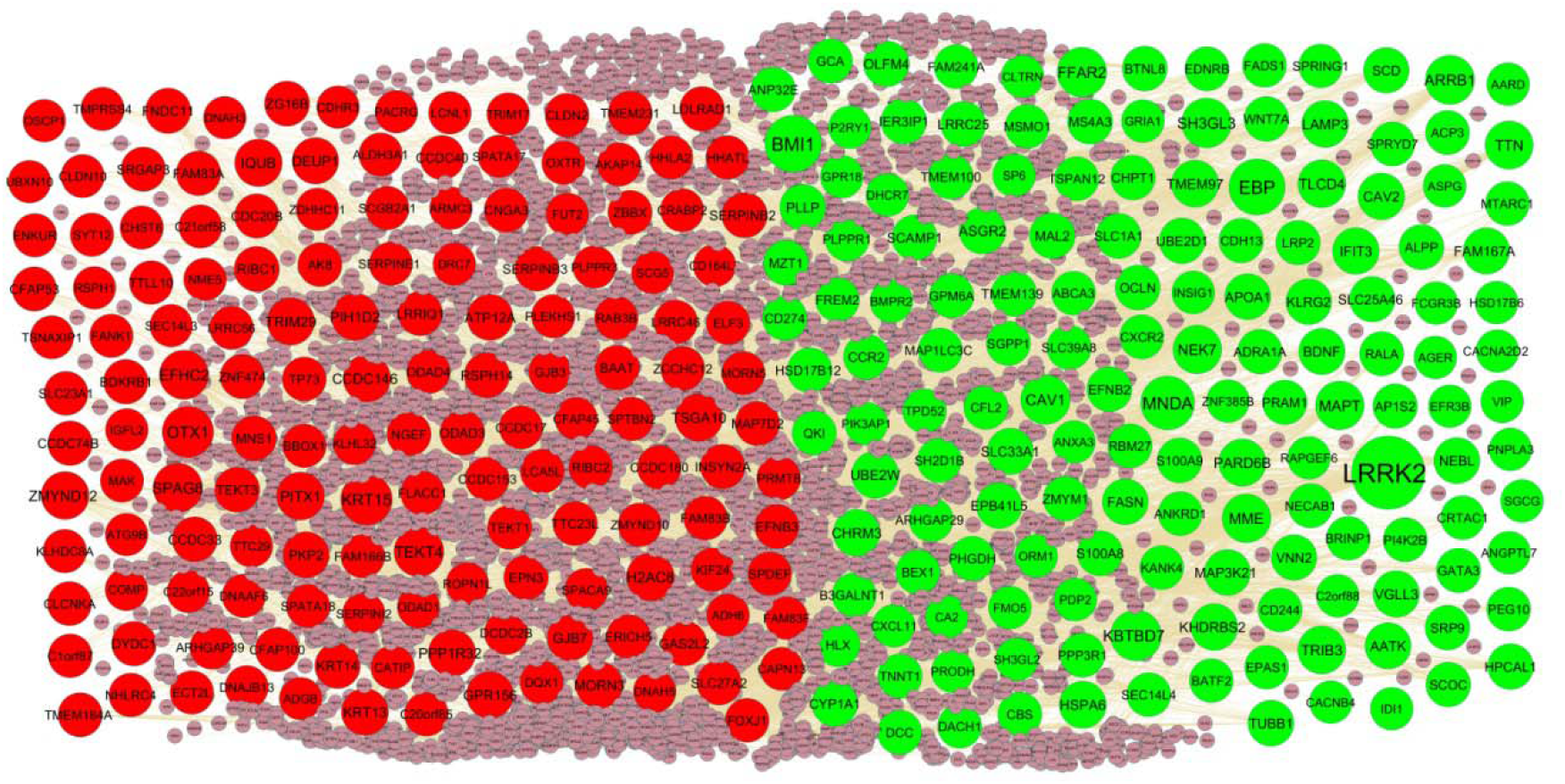
PPI network of DEGs. Up regulated genes are marked in parrot green; down regulated genes are marked in red

**Fig. 4.**
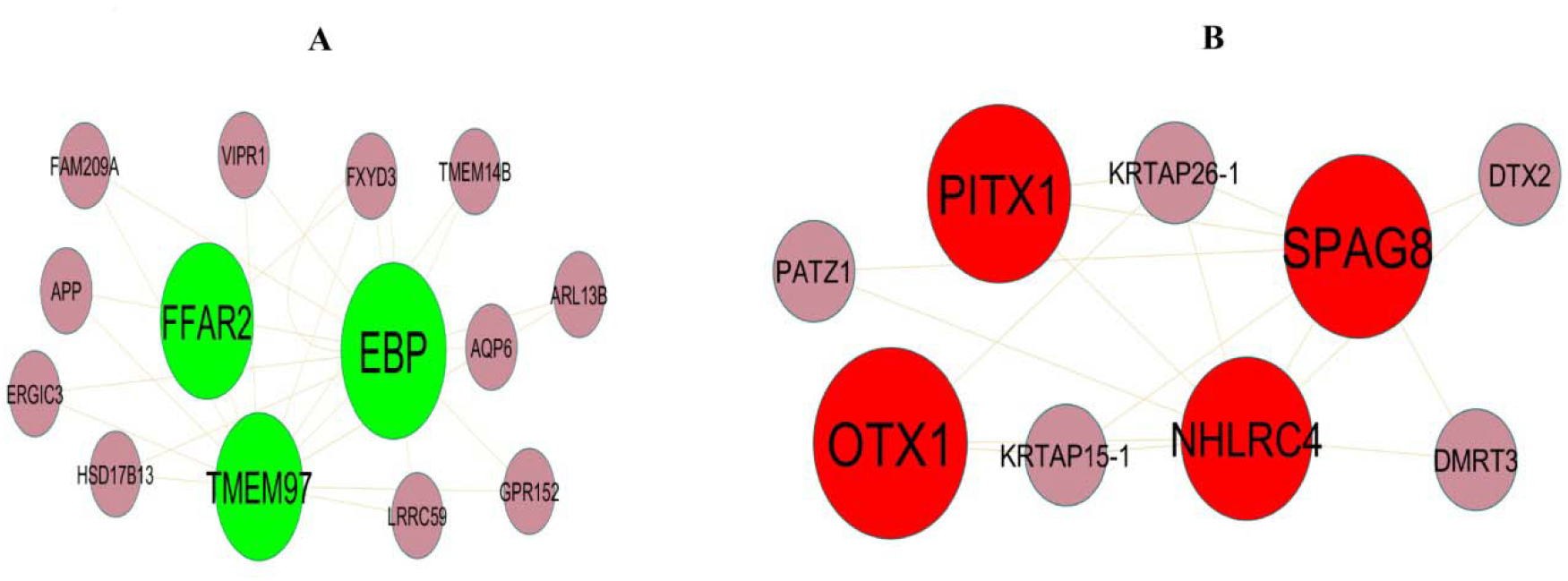
Modules selected from the PPI network. (A) The most significant module was obtained from PPI network with 14 nodes and 26 edges for up regulated genes (B) The most significant module was obtained from PPI network with 9 nodes and 17 edges for down regulated genes. Up regulated genes are marked in parrot green; down regulated genes are marked in red

**Table 4.**
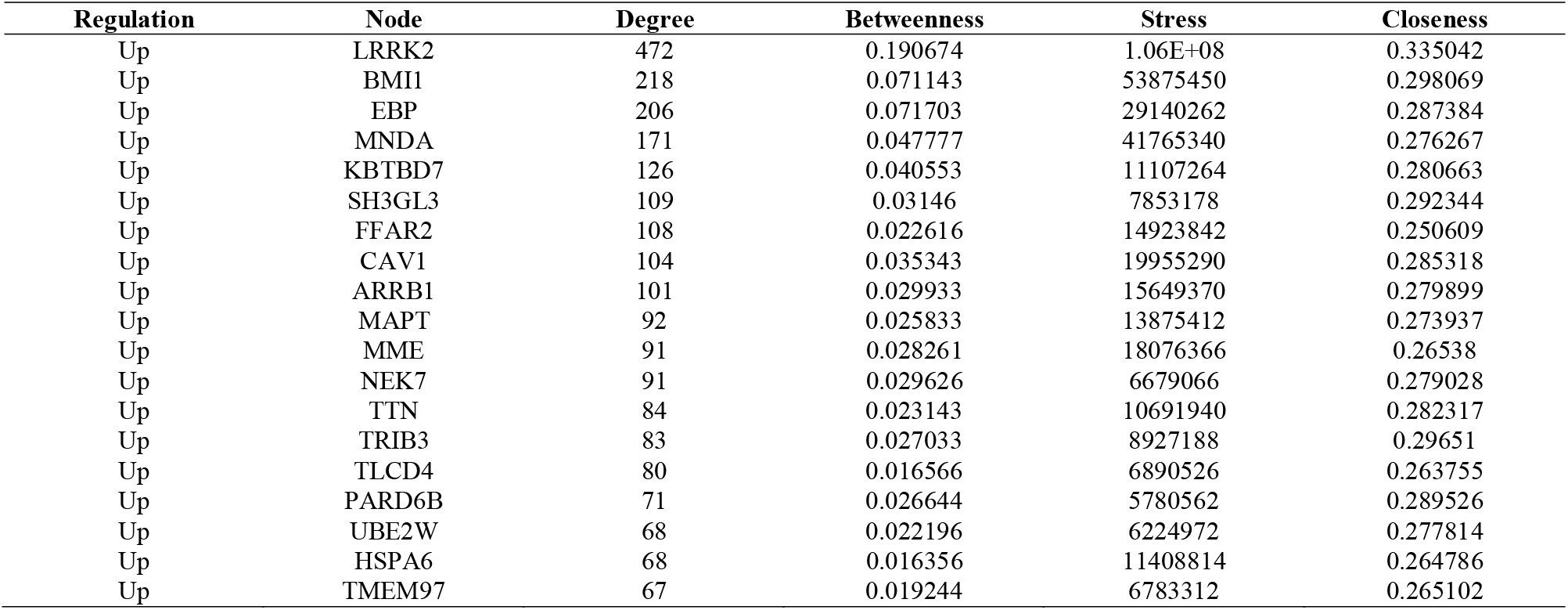
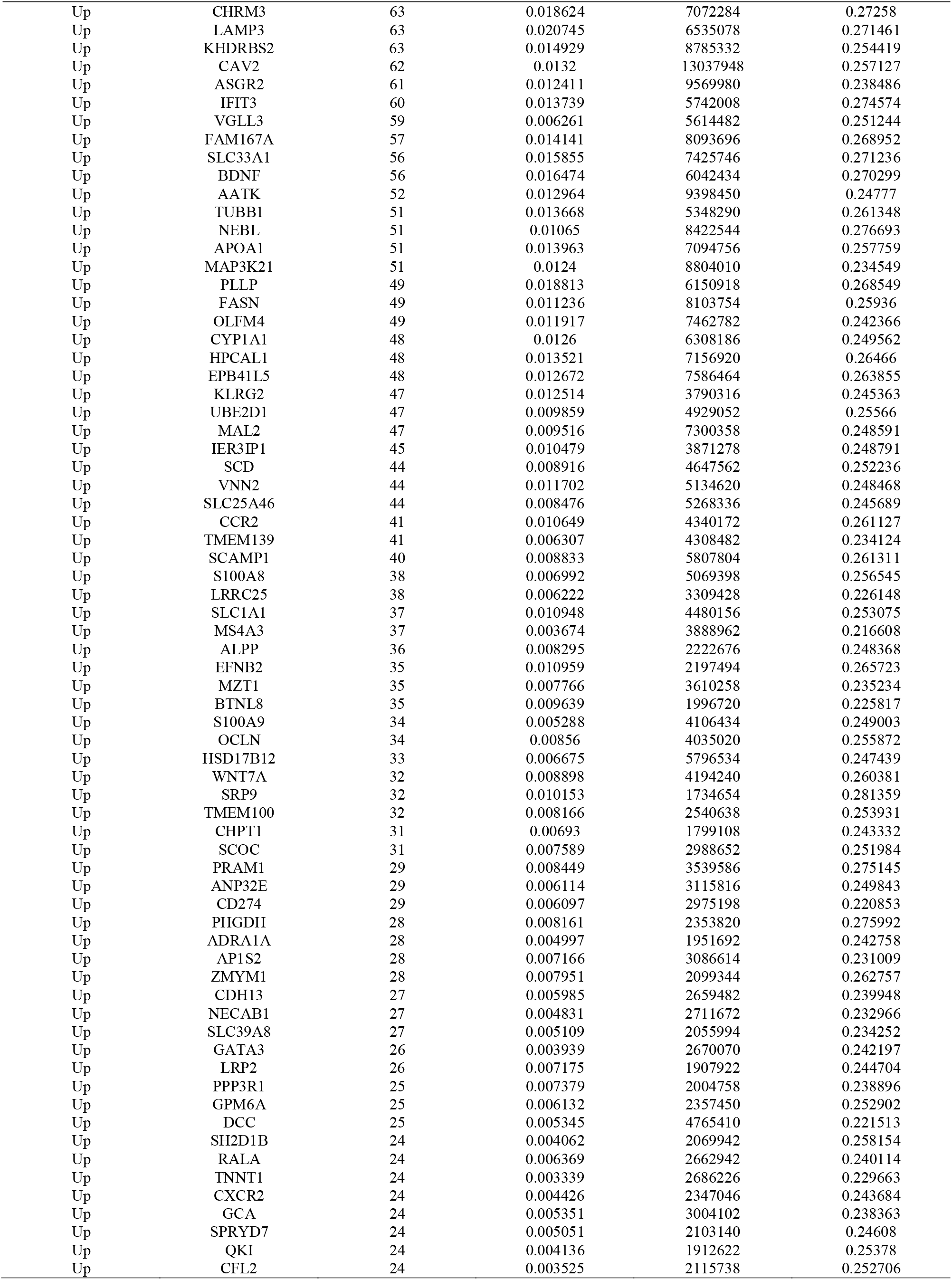
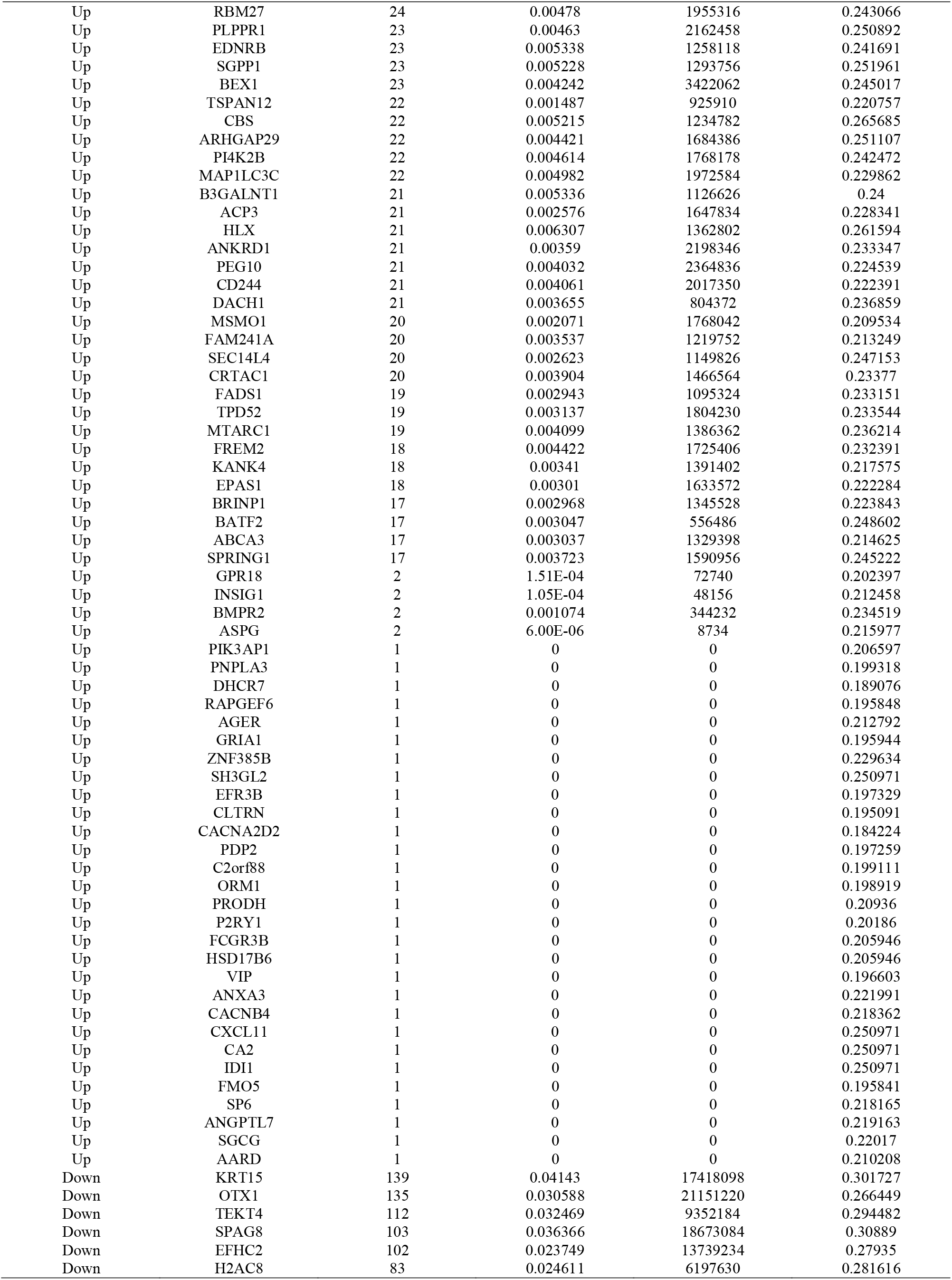
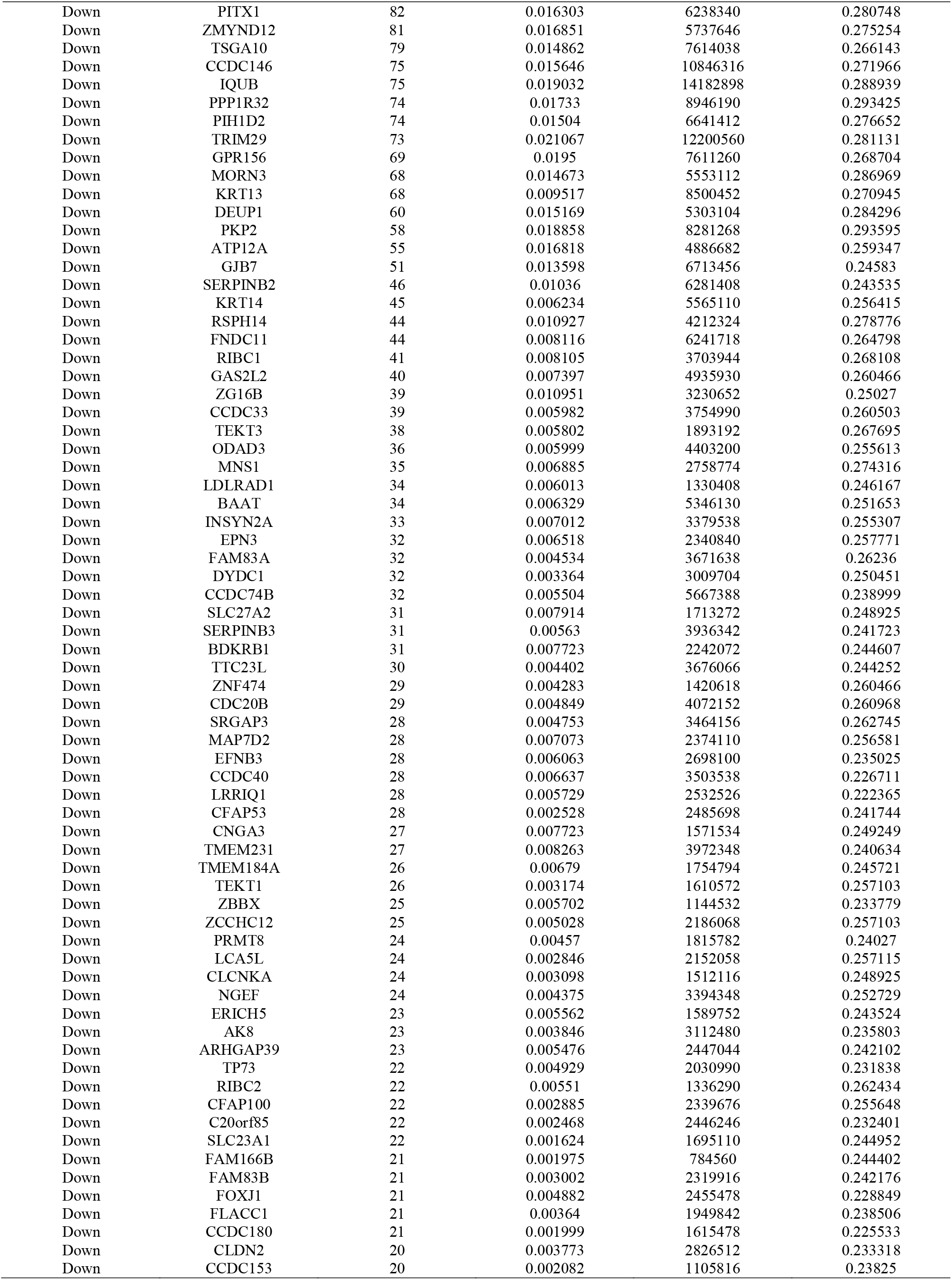
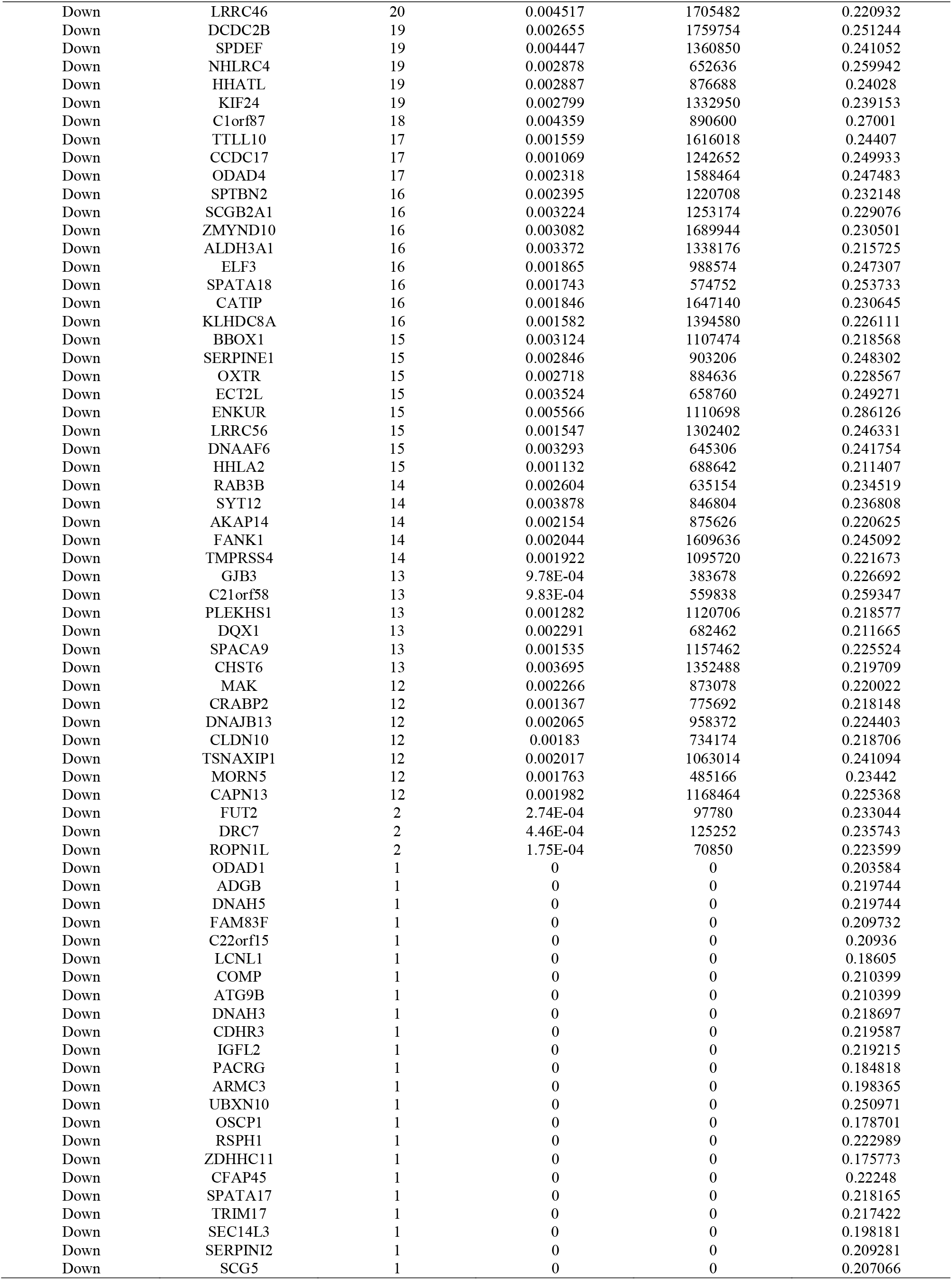
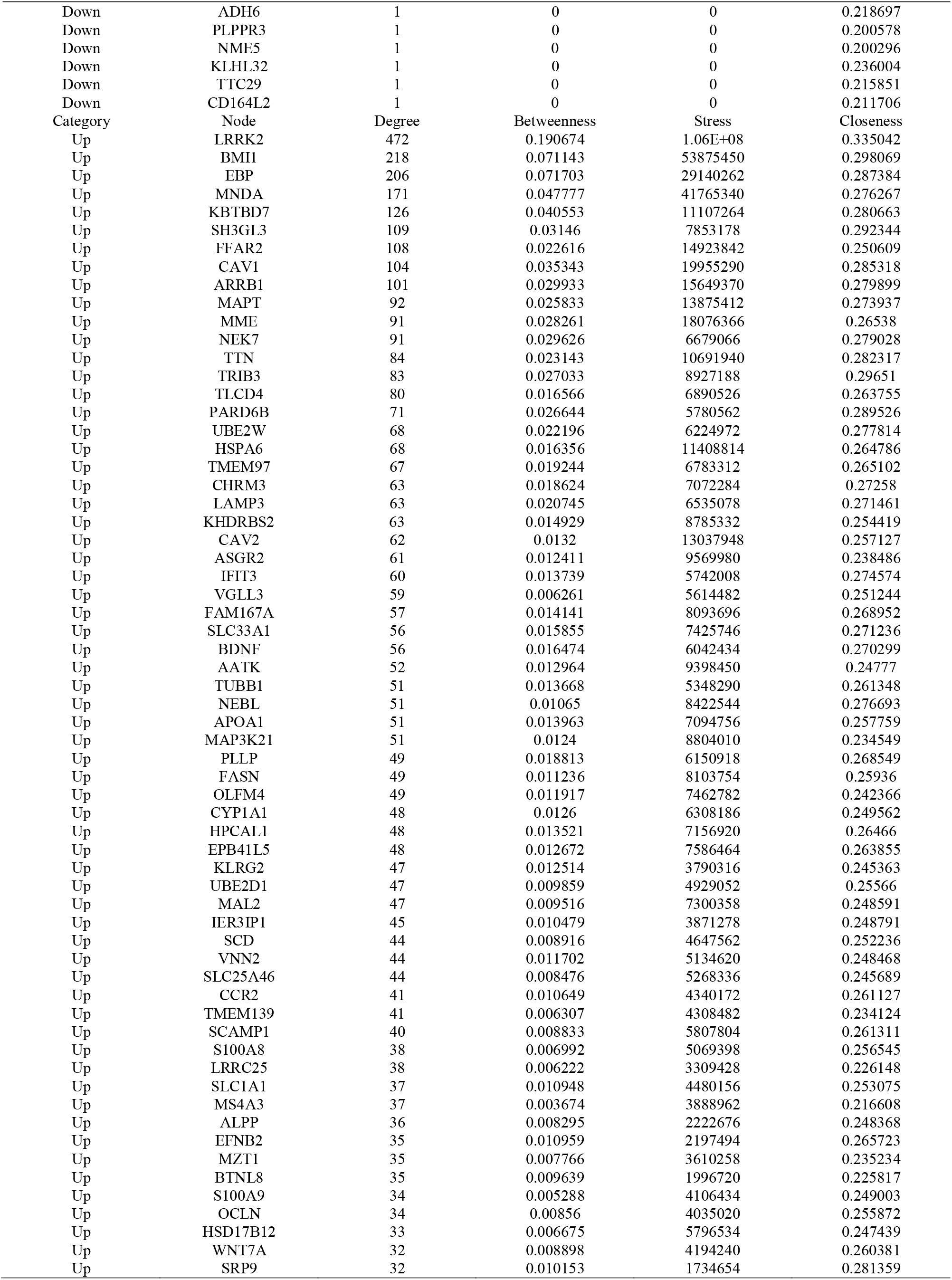
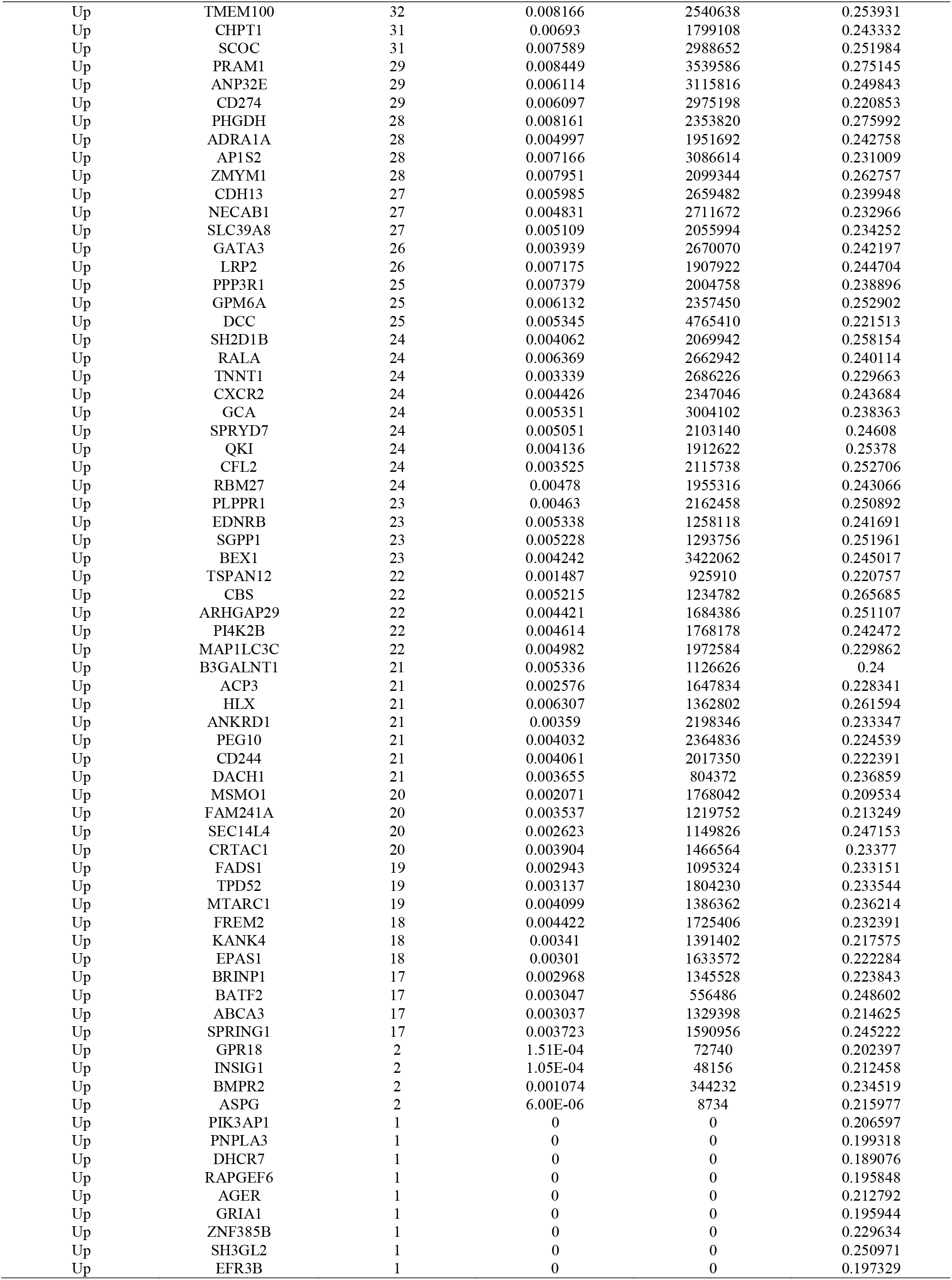
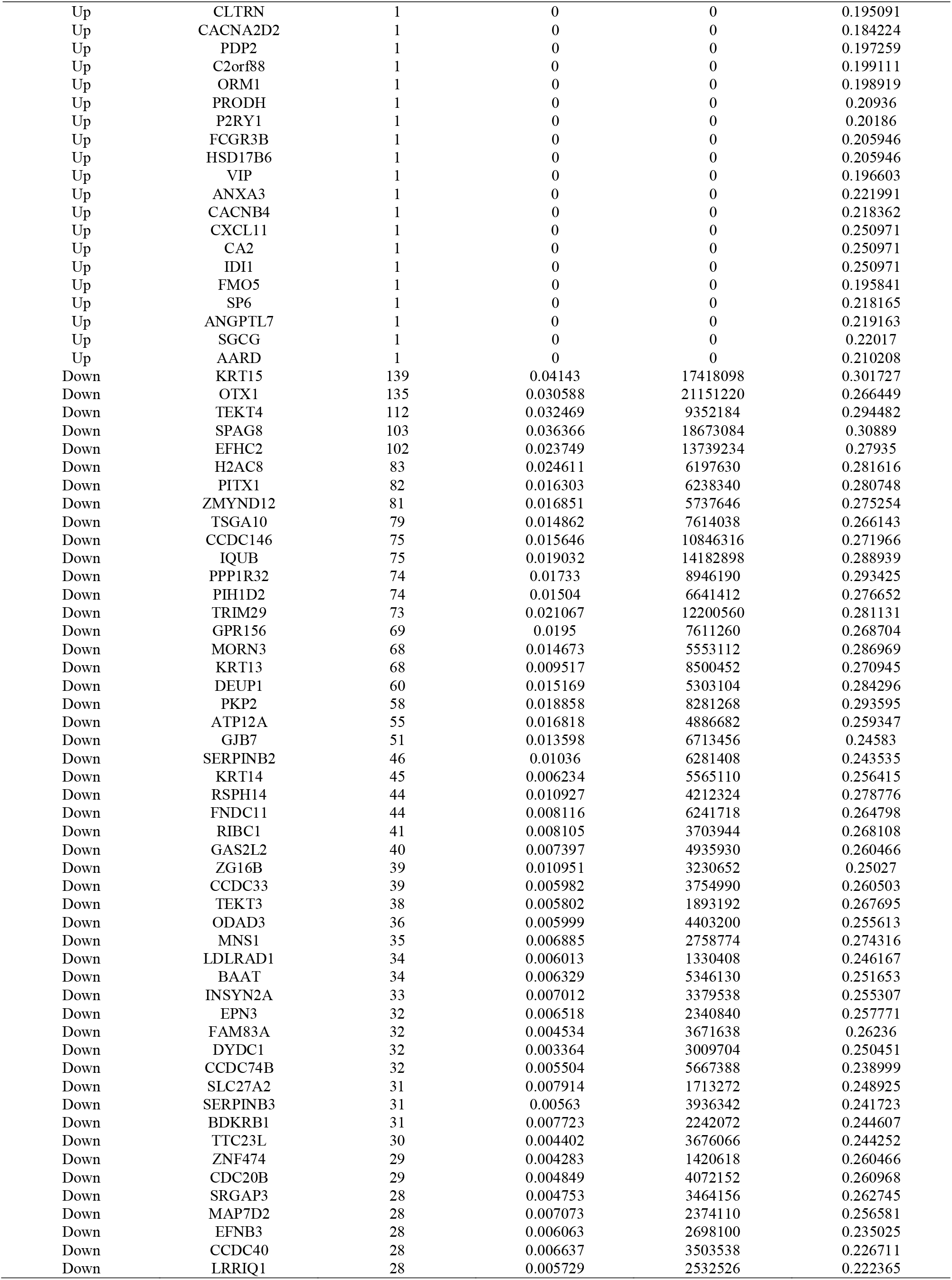
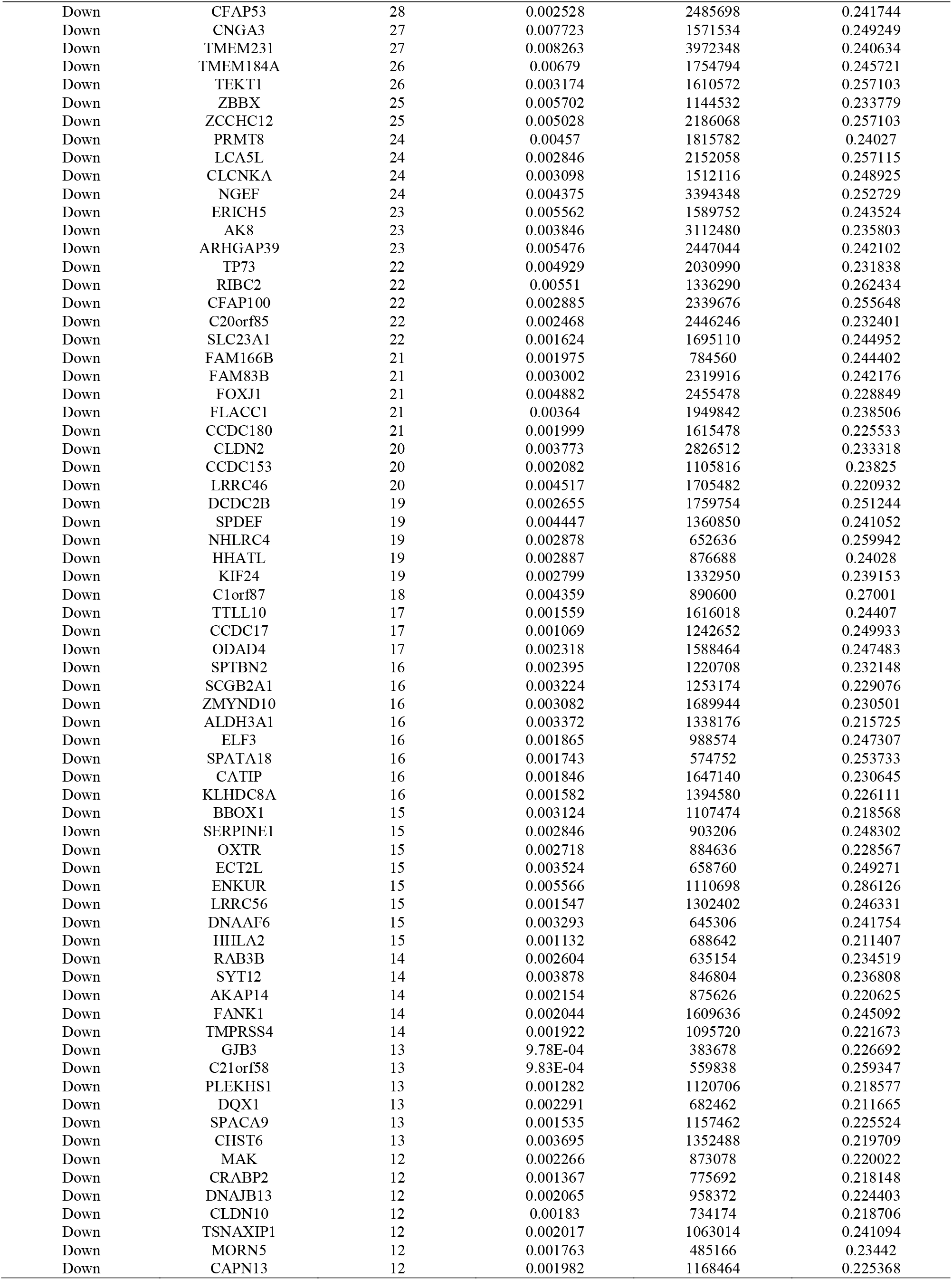
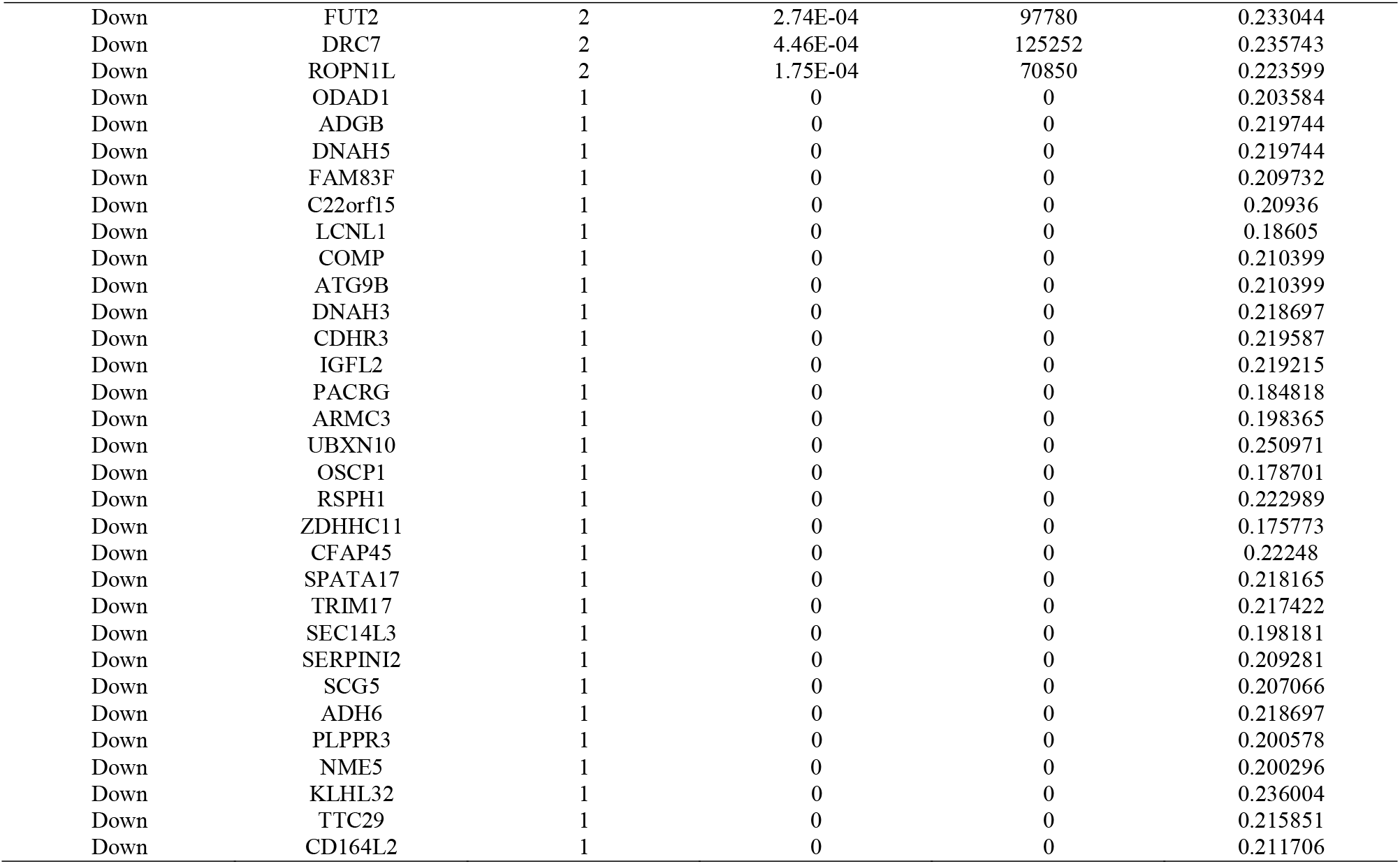
Topology table for up and down regulated genes.

### Construction of the miRNA-hub gene regulatory network

The database miRNet was adopted for predicting the target miRNA of the hub genes. The possible miRNA-hub gene network was constructed to accurately investigate the molecular mechanism, there were 2299 nodes (miRNA: 2023; hub gene: 276) and 10240 edges (Fig. 5). The 217 miRNAs (ex: hsa-mir-6830-5p) interacted with PARD6B; 156 miRNAs (ex: hsa-mir-362-3p) interacted with NEK7; 115 miRNAs (ex: hsa-mir-15b) interacted with BMI1; 115 miRNAs (ex: hsa-mir-766-5p) interacted with CAV1; 80 miRNAs (ex: hsa-mir-8057) interacted with TRIB3; 42 miRNAs (ex: hsa-mir-302a-3p) interacted with GPR156; 39 miRNAs (ex: hsa-mir-19b-3p) interacted with PITX1; 27 miRNAs (ex: hsa-mir-4524a-3p) interacted with TRIM29; 24 miRNAs (ex: hsa-mir-1537-5p) interacted with OTX1; 11 miRNAs (ex: hsa-mir-941) interacted with CCDC146 (Table 5).

**Fig. 5.**
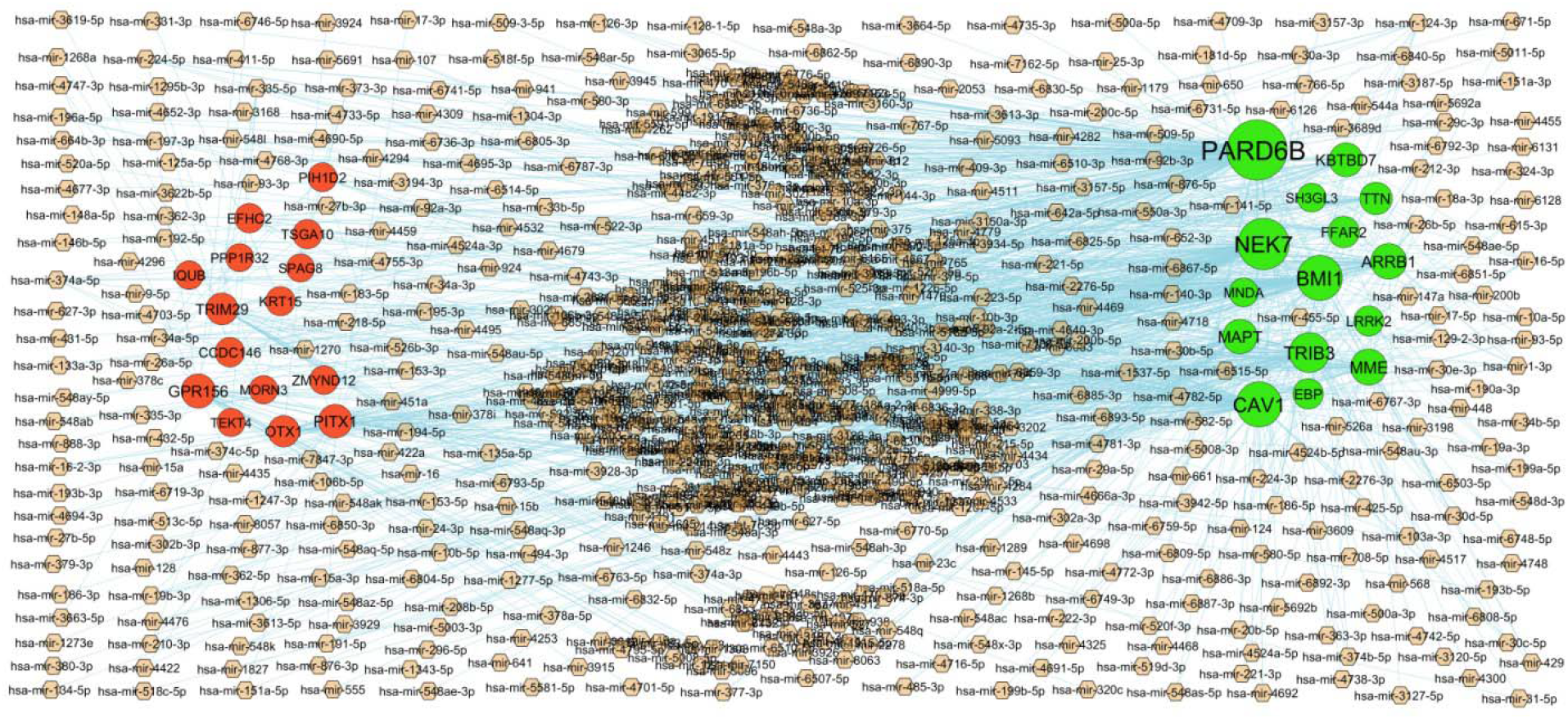
Hub gene - miRNA regulatory network. The chocolate color diamond nodes represent the key miRNAs; up regulated genes are marked in green; down regulated genes are marked in red.

**Table 5.**
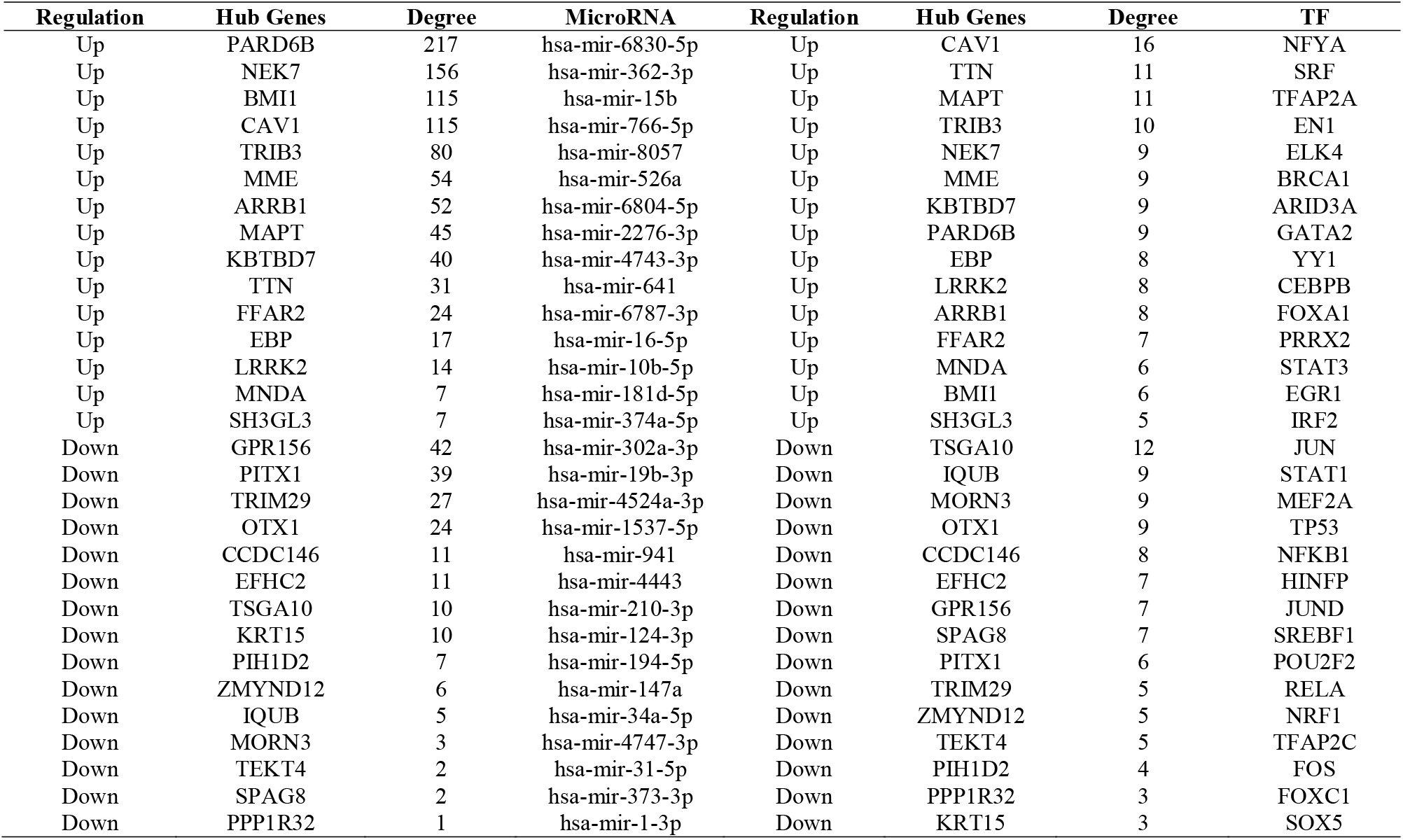
MiRNA - hub gene and TF – hub gene topology table.

### Construction of the TF-hub gene regulatory network

The database NetworkAnalyst was adopted for predicting the target TF of the hub genes. The possible TF-hub gene network was constructed to accurately investigate the molecular mechanism, there were 358 nodes (TF: 79; hub gene: 279) and 2169 edges (Fig. 6). The 16 TFs (ex: NFYA) interacted with CAV1; 11 TFs (ex: SRF) interacted with TTN; 11 TFs (ex: TFAP2A) interacted with MAPT; 10 TFs (ex: EN1) interacted with TRIB3; 9 TFs (ex: ELK4) interacted with NEK7; 12 TFs (ex: JUN) interacted with IQUB; 9 TFs (ex: STAT1) interacted with MORN3; 9 TFs (ex: MEF2A) interacted with OTX1; 9 TFs (ex: TP53) interacted with CCDC146; 8 TFs (ex: NFKB1) interacted with EFHC2 (Table 5).

**Fig. 6.**
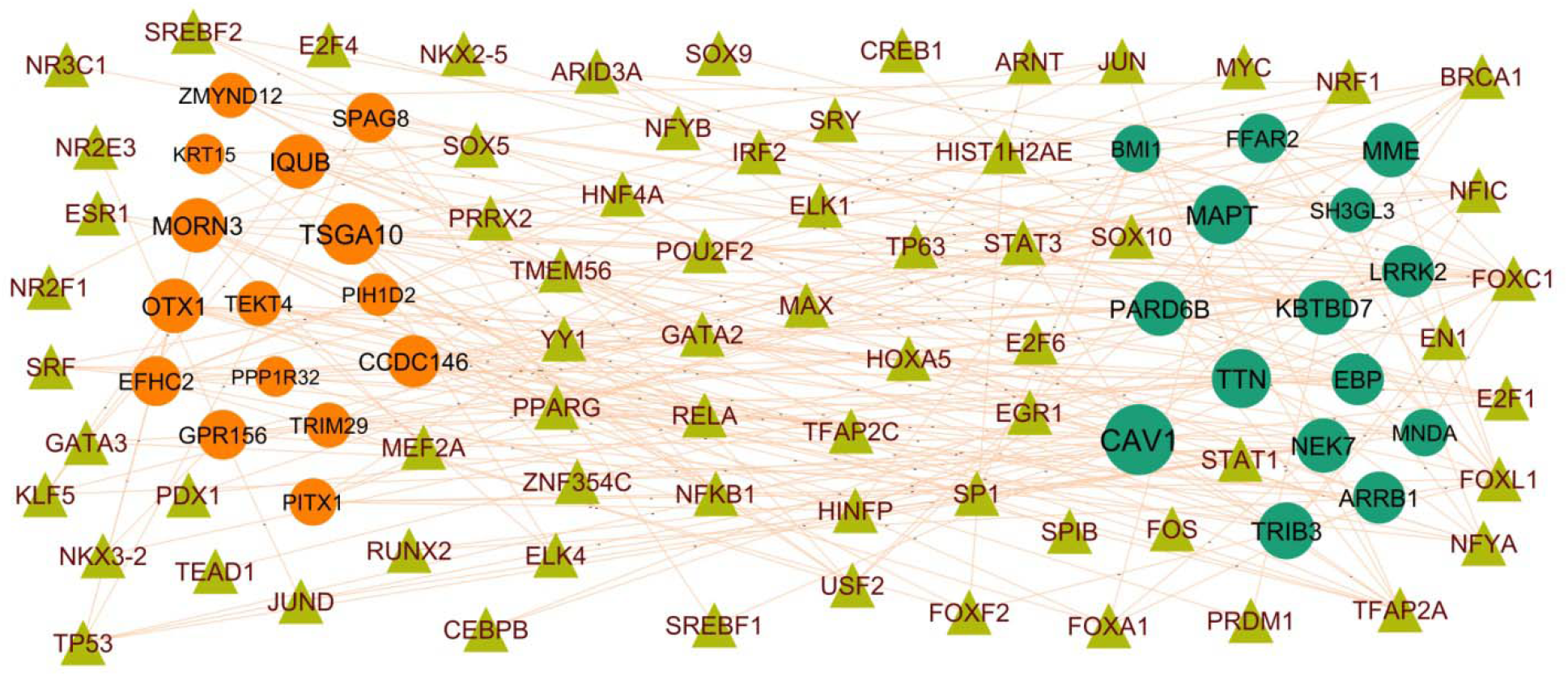
Hub gene - TF regulatory network. The olive color triangle nodes represent the key TFs; up regulated genes are marked in green; down regulated genes are marked in red.

### Receiver operating characteristic curve (ROC) analysis

Hence, the performance of hub genes in predicting disease samples was evaluated by plotting the ROC curves of the hub genes. There were hub genes related to IPF differences. ROC curves of each of the hub genes associated with IPF in normal control and IPF samples were constructed. The results revealed hub genes with predicted AUC > 0.8, namely LRRK2, BMI1, EBP, MNDA, KBTBD7, KRT15, OTX1, TEKT4, SPAG8 and EFHC2 (Fig. 7). The results show that these hub genes could successfully distinguish between IPF and normal control samples.

**Fig. 7.**
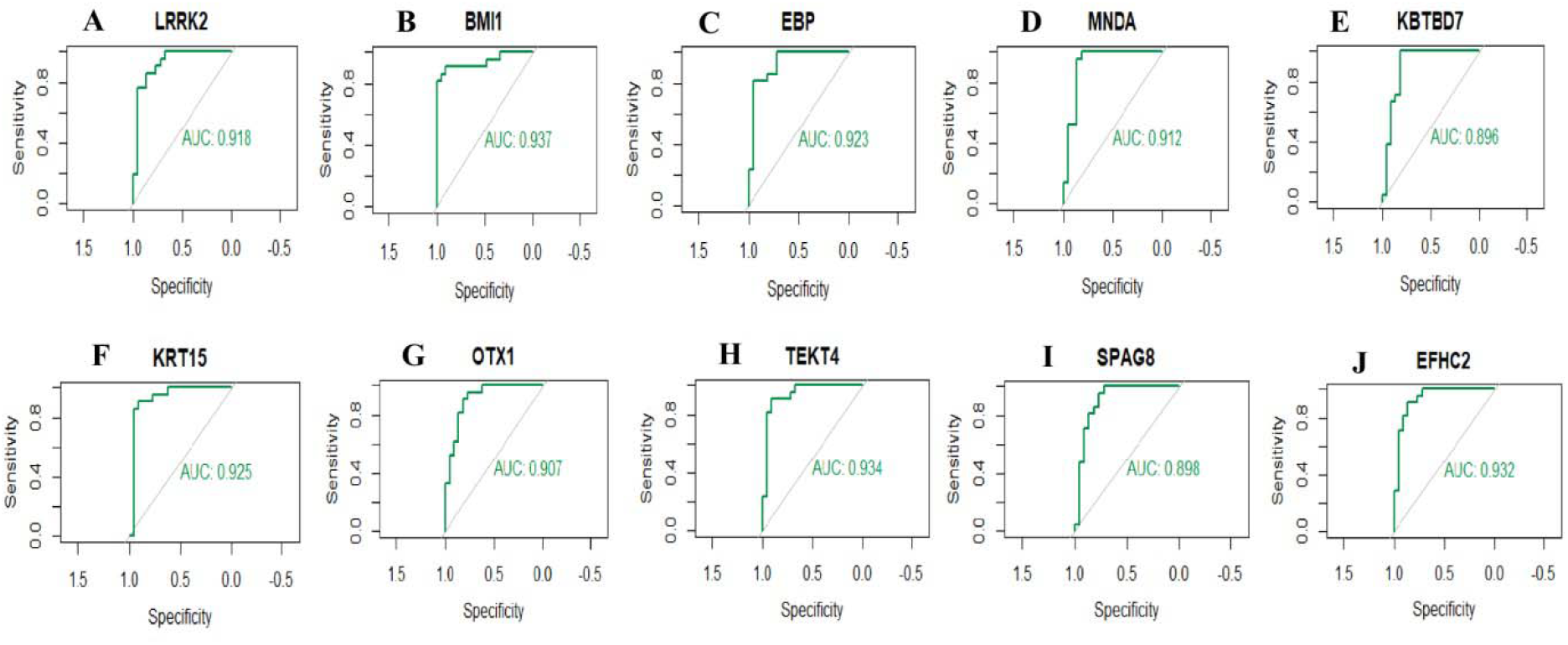
ROC curve analyses of hub genes. A) LRRK2 B) BMI1 C) EBP D) MNDA E) KBTBD7 F) KRT15 G) OTX1 H) TEKT4 I) SPAG8 J) EFHC2

## Discussion

Despite advances in adjunctive pharmacotherapy of IPF, it is still the leading threat to human health in lung diseases [51]. Thus, successful screening techniques and accurate diagnosis remain the great challenges for decreasing the incidence of IPF. In the current investigation, integrated bioinformatics and NGS data analysis was used to identify the potential hub genes related to IPF. By performing DEGs analysis, 479 up regulated and 479 down regulated DEGs were successfully identified (|logFC| > 0.512 for up regulated genes, |logFC| < −0.831 for down regulated genes and adjust P-value < .05), respectively. Involvement of SLCO1A2 [52], OLFM4 [53], RTKN2 [54], CYP1A1 [55] and MUC5AC [56] plays a key role in rheumatoid arthritis development. OLFM4 [57], RTKN2 [58], CYP1A1 [59], MUC5AC [60], CYP2A6 [61], PCK1 [62] and PITX1 [63] have been reported to encourage the development of lung cancer. Recent studies have demonstrated that the OLFM4 [64] is associated with obesity. OLFM4 [65] and AGTR2 [66] are important in the development of viral respiratory diseases. Recent studies have proposed that the CYP1A1 [67] and AGTR2 [68] are associated with systemic sclerosis. CYP1A1 [67] is a pathogenic gene for systemic lupus erythematosus. A previous study found that CYP1A1 [69] and MUC5AC [70] are positively correlated with the severity of pneumonia. Studies had shown that CYP1A1 [71] and AGTR2 [72] were associated with heart failure. CYP1A1 [73], HHIP (hedgehog interacting protein) [74], MUC5AC [75] and CYP2A6 [76] might be a potential therapeutic targets for chronic obstructive pulmonary disease treatment. Previous studies have reported that the CYP1A1 [77], MUC5AC [78] and ATP12A [79] are a key regulator of airway inflammation. Studies have found that HHIP (hedgehog interacting protein) [80], MUC5AC [81], CYP2A6 [82] and PCK1 [83] are altered expressed in obesity. AGTR2 [68], MUC5AC [70] and ATP12A [84] are a potential targets for IPF therapy. CYP1A1 [85], HHIP (hedgehog interacting protein) [86], AGTR2 [87], CYP2A6 [88] and PCK1 [83] expression had been confirmed in diabetes mellitus. In the light of important roles in cells, these DEGs in IPF might represent potential prognostic, diagnostics biomarkers and/or therapeutic targets for IPF.

In the current investigations, bioinformatics methods are promising methods to analyze the critical genes in GO terms and pathways, which might provide novel clues for diagnosis, therapy, and prognosis of IPF. Signaling pathways include GPCR ligand binding [89], neutrophil degranulation [90], immune system [91], metabolism of lipids [92] and signal transduction [93] made great contribution to the development of IPF. MAP3K15 [94], PRTN3 [95], CX3CR1 [96], AGRP (agouti related neuropeptide) [97], MPO (myeloperoxidase) [98], CD5L [99], S100A8 [100], NPR3 [101], VEGFD (vascular endothelial growth factor D) [102], CXCL11 [103], IL1A [104], CBS (cystathionine beta-synthase) [105], WNT7A [106], SCD (stearoyl-CoA desaturase) [107], LRP2 [108], SLC6A4 [109], BDNF (brain derived neurotrophic factor) [110], CXCL10 [111], ANGPTL7 [112], S100A9 [113], NPY1R [114], IL1B [115], GPIHBP1 [116], CYP1B1 [117], CD36 [118], MACROD2 [119], TRIB3 [120], SPX (spexin hormone) [121], PCSK9 [122], GPD1 [123], CDH13 [124], FFAR4 [125], FGF2 [126], FASN (fatty acid synthase) [127], DGAT2 [128], DACH1 [129], PNPLA3 [130], FGF9 [131], SLC7A11 [132], CLIC5 [133], VIP (vasoactive intestinal peptide) [134], SMAD6 [135], BMPR2 [136], APOA1 [137], INSIG1 [138], TLR3 [139], NLRP12 [140], ADRB1 [141], TLR8 [142], GATA3 [143], CCR2 [144], TLR7 [145], CCRL2 [146], BMPER (BMP binding endothelial regulator) [147], CAV1 [148], TFPI (tissue factor pathway inhibitor) [149], FADS1 [150], SUCNR1 [151], CADM2 [152], SLC19A3 [153], SGCG (sarcoglycan gamma) [154], ADH1B [155], NEGR1 [156], HSD17B12 [157], OXTR (oxytocin receptor) [158] and ANKK1 [159] were frequently altered in obesity. MAP3K15 [94], CX3CR1 [160], S100A12 [161], PF4 [162], FFAR2 [163], MPO (myeloperoxidase) [98], HMGCS2 [164], F11 [165], S100A8 [100], GRIA1 [166], NPR3 [167], CXCL11 [103], CBS (cystathionine beta-synthase) [105], WNT7A [106], AQP4 [168], SCD (stearoyl-CoA desaturase) [169], SLC6A4 [170], CXCL10 [171], S100A9 [172], NPY1R [173], IL1B [174], CXCR1 [175], CXCR2 [176], GPIHBP1 [177], WNT3A [178], APOH (apolipoprotein H) [179], CHRM3 [180], CD36 [181], TRIB3 [182], PCSK9 [183], ACVR1C [184], GPD1 [123], FFAR4 [185], GPX3 [186], FGF2 [187], FASN (fatty acid synthase) [188], DGAT2 [189], DACH1 [190], PNPLA3 [191], FGF9 [192], SLC7A11 [193], VIP (vasoactive intestinal peptide) [194], KL (klotho) [195], UBE2D1 [196], APOA1 [197], RASGRF1 [198], LRRK2 [199], TLR3 [200], OCLN (occludin) [201], SLC22A3 [202], LIFR (LIF receptor subunit alpha) [203], TLR8 [142], GATA3 [204], CCR2 [205], NEK7 [206], CD274 [207], TLR7 [208], CCRL2 [146], EFNB2 [209], CAV1 [210], TRPC3 [211], DLL4 [212], ANXA3 [213], TFPI (tissue factor pathway inhibitor) [214], FADS1 [215], GPER1 [216], SUCNR1 [217], CADM2 [218], SLC19A3 [219], ADH1B [220], NEGR1 [156], HSD17B12 [157], KIF6 [221], UCN3 [222], ANKK1 [223], AQP5 [224] and HCN4 [225] have been reported to be associated with diabetes mellitus. PRTN3 [226], CX3CR1 [227], S100A12 [228], CSF2 [229], FGG (fibrinogen gamma chain) [230], LHX9 [231], MPO (myeloperoxidase) [232], F11 [233], S100A8 [234], CXCL11 [235], BPI (bactericidal permeability increasing protein) [236], BDNF (brain derived neurotrophic factor) [237], CXCL10 [238], S100A9 [239], IL1B [240], CXCR1 [241], CXCR2 [242], CYP1B1 [243], EDNRB (endothelin receptor type B) [244], CEBPA (CCAAT enhancer binding protein alpha) [245], CDH13 [246], GPX3 [247], FGF2 [248], SHH (sonic hedgehog signaling molecule) [249], VIP (vasoactive intestinal peptide) [250], KL (klotho) [251], SMAD6 [252], BMPR2 [253], APOA1 [254], TLR3 [255], GATA3 [256], CCR2 [257], CAV1 [258], TRPC3 [259], EPAS1 [260], SIGLEC14 [261], MAPK15 [262], DNAH5 [263] and AQP5 [264] were identified to be associated with chronic obstructive pulmonary disease. DEFA3 [265], CX3CR1 [266], S100A12 [161], TUBB1 [267], ANKRD1 [268], ADRA1A [269], FGG (fibrinogen gamma chain) [270], AGER (advanced glycosylation end-product specific receptor) [271], PF4 [272], FFAR2 [273], MPO (myeloperoxidase) [274], CD5L [275], HMGCS2 [164], RXFP1 [276], F11 [277], S100A8 [278], PGLYRP1 [279], VEGFD (vascular endothelial growth factor D) [280], CHRM2 [281], CBS (cystathionine beta-synthase) [282], BPI (bactericidal permeability increasing protein) [283], LRP2 [284], BDNF (brain derived neurotrophic factor) [285], GCOM1 [286], CXCL10 [287], ANGPTL7 [288], PRODH (proline dehydrogenase 1) [289], P2RY1 [290], LRRN4 [291], S100A9 [292] CXCR1 [293], CXCR2 [293], GPIHBP1 [294], TNNT1 [295], WNT3A [296], BMI1 [297], CYP1B1 [298], FCN3 [299], TTN (titin) [300], STC1 [301], CD36 [302], MYZAP (myocardial zonula adherens protein) [303], TRIB3 [304], GPR18 [305], TNNC1 [306], SPX (spexin hormone) [121], SYNPO2L [307], PCSK9 [308], GPD1 [309], FFAR4 [310], GPX3 [311], FGF2 [187], ACKR4 [312], NDUFC2 [313], KBTBD7 [314], SHH (sonic hedgehog signaling molecule) [315], DACH1 [316], PNPLA3 [317], FGF9 [192], SLC7A11 [193], SGPP1 [318], VIP (vasoactive intestinal peptide) [319], KCNJ2 [320], KL (klotho) [321], SMAD6 [135], BMPR2 [322], APOA1 [323], CALCRL (calcitonin receptor like receptor) [324], INSIG1 [325], RASGRF1 [198], LRRK2 [326], TLR3 [327], ADRB1 [328], SLC22A3 [329], CA2 [330], SNX10 [331], LIFR (LIF receptor subunit alpha) [332], TLR8 [333], CMPK2 [334], GATA3 [335], RSPO2 [336], CCR2 [205], NEK7 [337], TLR7 [338], BEX1 [339], EFNB2 [340], CAV1 [341], ARRB1 [342], TRPC3 [343], CR1 [344], PEG10 [345], DLL4 [346], MEFV (MEFV innate immuity regulator, pyrin) [347], TFPI (tissue factor pathway inhibitor) [348], EPAS1 [349], FADS1 [215], DKK2 [350], CACNA2D2 [351], DPP6 [352], KCNA4 [353], PCDH17 [354], SUSD2 [355], PHACTR2 [356], DNAH9 [357], DNAH11 [358], CFAP45 [359], DNAH5 [360], FOXJ1 [361], MNS1 [299], KIF6 [221], DRD5 [362], UCN3 [363], OXTR (oxytocin receptor) [364], ANKK1 [365] and HCN4 [366] have been shown to be a biomarker of heart failure. Study demonstrated that PI3 [367], CX3CR1 [368], S100A12 [369], MPO (myeloperoxidase) [370], CD5L [371], S100A8 [372], CXCL11 [373], BPI (bactericidal permeability increasing protein) [374], AQP4 [375], BDNF (brain derived neurotrophic factor) [376], CXCL10 [377], CCL8 [378], S100A9 [379], IL1B [240], CXCR1 [380], CXCR2 [381], ABCA3 [382], GPR18 [383], VIP (vasoactive intestinal peptide) [384], KL (klotho) [385], TLR3 [386], NLRP12 [387], GATA3 [388], CCR2 [389], TLR7 [390], CAV1 [391], CR1 [392], DLL4 [393] and AQP5 [394] were essential for the induction of airway inflammation. CX3CR1 [395], S100A12 [396], MPO (myeloperoxidase) [397], RXFP1 [398], S100A8 [399], CXCL11 [373], CBS (cystathionine beta-synthase) [400], WNT7A [401], BDNF (brain derived neurotrophic factor) [402], CXCL10 [403], CCL8 [404], FCGR3B [405], S100A9 [406], IL1B [407], CXCR2 [408], WNT3A [409], BMI1 [410], STC1 [411], ABCA3 [412], CD36 [413], TRIB3 [414], GPX3 [415], FGF2 [416], FASN (fatty acid synthase) [417], SHH (sonic hedgehog signaling molecule) [418], DACH1 [419], FGF9 [420], SLC7A11 [421], VIP (vasoactive intestinal peptide) [422], KL (klotho) [423], BMPR2 [424], APOA1 [425], LRRK2 [426], TLR3 [427], GATA3 [428], RSPO2 [429], CCR2 [430], NEK7 [431], BMPER (BMP binding endothelial regulator) [432], CAV1 [433], CR1 [434], TFPI (tissue factor pathway inhibitor) [435], AP1S2 [436], FOXJ1 [437], AQP5 [438], MUC16 [439] and MUC4 [440] could be used as a therapeutic target for IPF. CX3CR1 [441], S100A12 [442], PF4 [443], MPO (myeloperoxidase) [444], WNT7A [445], SLC6A4 [446], BDNF (brain derived neurotrophic factor) [447], CXCL10 [448], NEK7 [449], CYP1B1 [450], ABCA3 [451], TRIB3 [452], PCSK9 [453], FGF2 [454], ACKR4 [455], FASN (fatty acid synthase) [456], VIP (vasoactive intestinal peptide) [457], KL (klotho) [458], BMPR2 [459], APOA1 [323], TLR3 [460], CCR2 [461], TLR7 [462], CAV1 [463], WWC2 [464], TFPI (tissue factor pathway inhibitor) [465], EPAS1 [466] and CCDC40 [467] could serve as a potential therapeutic for pulmonary hypertension treatment. A previous study reported that the genes include CX3CR1 [468], CSF2 [469], CLDN18 [470], TRIM58 [471], PF4 [472], FFAR2 [473], MPO (myeloperoxidase) [474], CD5L [475], SH3GL2 [476], ITGA2B [477], S100A8 [478], VEGFD (vascular endothelial growth factor D) [479], CXCL11 [480], IL1A [481], WNT7A [482], SSTR1 [483], AQP4 [484], SCD (stearoyl-CoA desaturase) [485], SLC6A4 [486], BDNF (brain derived neurotrophic factor) [487], CXCL10 [488], ODAM (odontogenic, ameloblast associated) [489], CASP5 [490], CCL8 [491], TMEM100 [492], S100A9 [493], IL1B [494], CXCR1 [495], CXCR2 [496], WNT3A [497], BMI1 [498], CYP1B1 [499], FCN3 [500], TTN (titin) [501], SHISA3 [502], AZGP1 [503], ABCA3 [504], CD36 [505], EDNRB (endothelin receptor type B) [506], BTNL9 507], CEBPA (CCAAT enhancer binding protein alpha) [508], TRIB3 [509], TNNC1 [510], PCSK9 [511], P2RY13 [512], KITLG (KIT ligand) [513], CDH13 [514], GPX3 [515], FGF2 [416], FUT7 [516], FASN (fatty acid synthase) [517], NKD1 [518], FOXD1 [519], SLC1A1 [520], SHH (sonic hedgehog signaling molecule) [521], DACH1 [522], FGF9 [523], SLC7A11 [524], CLIC5 [525], MGAT3 [526], HSPA6 [527], TSPAN12 [528], SCAI (suppressor of cancer cell invasion) [529], VIP (vasoactive intestinal peptide) [530], SH3GL3 [531], KCNJ2 [532], KL (klotho) [533], UBE2D1 [534], SMAD6 [535], BMPR2 [536], APOA1 [537], TGFBR3 [538], RASGRF1 [539], LRRK2 [540], ATP8A2 [541], TLR3 [542], OCLN (occludin) [543], EMP2 [544], MNDA (myeloid cell nuclear differentiation antigen) [545], TLR8 [546], GATA3 [547], RSPO2 [548], CCR2 [549], EPB41L5 [550], CD274 [551], DDIAS (DNA damage induced apoptosis suppressor) [552], TLR7 [553], CCRL2 [554], BMPER (BMP binding endothelial regulator) [555], DUSP26 [556], CCBE1 [557], FZD8 [558], CAV1 [559], ARRB1 [560], CR1 [561], WWC2 [562], DLL4 [563], ANXA3 [564], EPAS1 [565], FADS1 [566], DKK2 [567], GPER1 [568], CADM2 [569], PARD6B [570], CACNA2D2 [571], ATP8A1 [572], PDZD2 [573], STXBP6 [574], ADH1A [575], GCA (grancalcin) [576], SUSD2 [577], EDIL3 [578], PHACTR2 [579], DNAH10 [580], CCDC65 [581], SPAG6 [582], MAPK15 [583], ENKUR (enkurin, TRPC channel interacting protein) [584], DNAH5 [585], PIERCE1 [586], TPPP3 [587], TTC21A [588], DLEC1 [589], SRCIN1 [590], PROM1 [591], AQP5 [592], SYT13 [593], TTC21A [594], SPTBN2 [595], MUC13 [596], MUC16 [597] and MUC4 [598] pathogenic genes were associated with the lung cancer. The altered expression of CX3CR1 [599], S100A12 [600], PF4 [601], MPO (myeloperoxidase) [602], RXFP1 [603], S100A8 [604], VEGFD (vascular endothelial growth factor D) [605], CXCL11 [606], IL1A [607], BPI (bactericidal permeability increasing protein) [608], SLC6A4 [609], CXCL10 [610], FCGR3B [611], S100A9 [612], IL1B [613], CXCR2 [614], CTNND2 [615], CD36 [616], PCSK9 [617], FGF2 [618], SHH (sonic hedgehog signaling molecule) [619], KL (klotho) [620], BMPR2 [621], TLR8 [622], GATA3 [623], CCR2 [624], TLR7 [625], CAV1 [626], TFPI (tissue factor pathway inhibitor) [627] and SPAG17 [628] have been identified in systemic sclerosis. A previous study found that CX3CR1 [629], S100A12 [630], MPO (myeloperoxidase) [631], CD5L [632], F11 [633], S100A8 [634], BPI (bactericidal permeability increasing protein) [635], AQP4 [636], MME (membrane metalloendopeptidase) [637], BDNF (brain derived neurotrophic factor) [638], CXCL10 [639], RNASE2 [640], FCGR3B [641], S100A9 [642], GPIHBP1 [643], AFF3 [644], APOH (apolipoprotein H) [645], FCN3 [646], PCSK9 [308], LILRA2 [647], APOA1 [648], LRRK2 [649], CD244 [650], TLR3 [651], TLR8 [652], GATA3 [653], CCR2 [654], IFIT3 [655], NEK7 [656], TLR7 [657], CAV1 [658], CR1 [659], TFPI (tissue factor pathway inhibitor) [660], GPER1 [661], SIGLEC14 [662], FOXJ1 [663], GABRP (gamma-aminobutyric acid type A receptor subunit pi) [664] and TSGA10 [665] are positively correlated with the severity of systemic lupus erythematosus, suggesting its potential as a biomarker for systemic lupus erythematosus. Altered expression of genes include CX3CR1 [666], S100A12 [667], CSF2 [668], MPO (myeloperoxidase) [669], CD5L [670], F11 [671], S100A8 [672], PGLYRP1 [673], VEGFD (vascular endothelial growth factor D) [674], CXCL11 [373], BPI (bactericidal permeability increasing protein) [675], CXCL10 [676], S100A9 [677], CXCR1 [678], CXCR2 [679], ABCA3 [680], CD36 [681], SHH (sonic hedgehog signaling molecule) [682], TLR3 [683], CLEC4D [684], CCR2 [685], NEK7 [686], TLR7 [687], CCRL2 [688] and CAV1 [689] accelerates pneumonia progression. CX3CR1 [666], CD177 [690], PF4 [691], FFAR2 [692], MPO (myeloperoxidase) [693], F11 [694], S100A8 [695], VEGFD (vascular endothelial growth factor D) [674], IL1A [696], BPI (bactericidal permeability increasing protein) [697], AQP4 [698], BDNF (brain derived neurotrophic factor) [699], CXCL10 [700], RNASE2 [701], FCGR3B [702], S100A9 [703], IL1B [704], CXCR2 [705], GPIHBP1 [294], CD36 [706], TRIB3 [707], PCSK9 [708], FGF2 [709], FASN (fatty acid synthase) [710], PNPLA3 [711], HSPA6 [712], VIP (vasoactive intestinal peptide) [713], TLR3 [683], ADRB1 [328], SPOCK2 [714], TLR8 [715], CCR2 [716], IFIT3 [717], NEK7 [718], TLR7 [687], EFNB2 [719], CAV1 [720], CR1 [721] and AQP5 [722] plays essential roles in viral respiratory diseases. Previous study confirmed that CX3CR1 [723], S100A12 [724], CD177 [725], PF4 [726], MPO (myeloperoxidase) [727], CD5L [728], F11 [729], S100A8 [730], PGLYRP1 [731], GPR15 [732], BPI (bactericidal permeability increasing protein) [733], AQP4 [734], BDNF (brain derived neurotrophic factor) [735], CXCL10 [736], FCGR3B [737], S100A9 [738], IL1B [739], CXCR1 [740], CXCR2 [741], AFF3 [742], WNT3A [743], FCN3 [744], AZGP1 [745], CD36 [746], PCSK9 [747], GPX3 [748], FGF2 [749], SHH (sonic hedgehog signaling molecule) [750], SLC7A11 [751], VIP (vasoactive intestinal peptide) [752], KL (klotho) [753], APOA1 [754], RASGRF1 [755], CD244 [756], TLR3 [757], NLRP12 [758], SNX10 [759], TLR8 [760], GATA3 [761], CCR2 [762], TLR7 [763], CCRL2 [764], EFNB2 [765], FZD8 [766], CAV1 [767], CR1 [768], MEFV (MEFV innate immuity regulator, pyrin) [769], SUCNR1 [770], GCA (grancalcin) [771] and FOXJ1 [663] are strongly associated with rheumatoid arthritis. S100A12 [772], MPO (myeloperoxidase) [773], S100A8 [774], CXCL10 [775], S100A9 [774], CD244 [775] and TLR7 [777] have been linked to dermatomyositis. MPO (myeloperoxidase) [778] have been proposed as novel biomarkers for mixed connective tissue disease progression. MPO (myeloperoxidase) [779], CXCL11 [780], IL1A [781], CXCL10 [782], FCGR3B [783], IL1B [784], TTN (titin) [300], BATF2 [785], VIP (vasoactive intestinal peptide) [786], TLR3 [787], CCR2 [788], TLR7 [789] and CR1 [790] are involved in the progression of sarcoidosis. A study indicates that MPO (myeloperoxidase) [791], RXFP1 [792], CXCL11 [793], CTNND2 [615], BMPR2 [794], TLR3 [795], CCR2 [796] and CAV1 [797] are positively correlated with scleroderma. CXCL11 [798] and CD244 [776] are involved in growth and development of polymyositis. Study demonstrated that IL1B [799], CXCR1 [800], FFAR4 [801], VIP (vasoactive intestinal peptide) [802] and GATA3 [803] were essential for the induction of gastroesophageal reflux disease. Therefore, it is necessary to perform GO term and pathway enrichment analysis in order to understand the interactions between DEGs and the associated biological processes. This discovery will help us better understand the pathogenesis of IPF and its complications and provide effective and novel treatment strategies for IPF.

Hub genes in the molecular pathogenesis of IPF were identified by using the IID software of Cytoscape. Studies have shown that LRRK2 [199] and FFAR2 [163] played a key role in diabetes mellitus development. LRRK2 [326], BMI1 [297], KBTBD7 [314] and FFAR2 [273] plays an important role in the development of heart failure. LRRK2 [426] and BMI1 [410] plays a vital role in the pathogenesis of IPF. LRRK2 [540], BMI1 [398], MNDA (myeloid cell nuclear differentiation antigen) [545], OTX1 [804], FFAR2 [473] and PITX1 [63] playing a pathogenic role in lung cancer. LRRK2 [649] was shown to be altered expressed in systemic lupus erythematosus and involved in the pathogenesis of systemic lupus erythematosus. A study has shown that FFAR2 [692] was involved in the **viral respiratory diseases.** Hub genes include EBP (EBP cholestenol delta-isomerase), KRT15, TEKT4, SPAG8, EFHC2, TMEM97 and NHLRC4 might participate in the pathophysiological process of IPF and its complications, and might become a novel molecular target for diagnosis and prognosis of IPF patients.

MiRNA-hub gene regulatory network and TF-hub gene regulatory network can be regarded as key to the understanding of pathogensisof IPF and might also lead to new therapeutic approaches. Studies have shown that PARD6B [570], BMI1 [498], CAV1 [559], TRIB3 [509], TTN (titin) [501], PITX1 [63], TRIM29 [805], OTX1 [804], NFYA (nuclear transcription factor Y subunit alpha) [806], TFAP2A [807], JUN (Jun proto-oncogene, AP-1 transcription factor subunit) [808], STAT1 [809], TP53 [810] and NFKB1 [811] plays an important role in **lung cancer.** Studies have shown that NEK7 [206], CAV1 [210], TRIB3 [182], STAT1 [812], TP53 [813] and NFKB1 [814] are closely related to the development of the, diabetes mellitus. Some paper reported that NEK7 [337], BMI1 [297], CAV1 [341], TRIB3 [304], TTN (titin) [300], hsa-mir-19b-3p [815], hsa-mir-941 [816], NFYA (nuclear transcription factor Y subunit alpha) [817], STAT1 [818], MEF2A [819], TP53 [820] and NFKB1 [821] altered expression is associated with development and progression of heart failure. Previous studies have demonstrated that NEK7 [431], BMI1 [410], CAV1 [433], TRIB3 [414], SRF (serum response factor) [822], STAT1 [823] and TP53 [824] are an independent prognostic factors in IPF and might be associated with IPF pathogenesis. NEK7 [656], CAV1 [658], TSGA10 [665], STAT1 [825], TP53 [826] and NFKB1 [827] are associated with systemic lupus erythematosus. NEK7 [686] and CAV1 [689] were an important participant in pneumonia. NEK7 [718], CAV1 [720], TRIB3 [707], hsa-mir-8057 [828], hsa-mir-1537-5p [829], STAT1 [830] and TP53 [832] were reported to induce viral respiratory diseases. NEK7 [449], TRIB3 [452], CAV1 [463], NFYA (nuclear transcription factor Y subunit alpha) [817], STAT1 [832] and TP53 [833] were significantly altered expression in pulmonary hypertension. CAV1 [148], TRIB3 [120], STAT1 [834], TP53 [835] and NFKB1 [836] are crucial for obesity pathogenesis. CAV1 [258], hsa-mir-19b-3p [837], SRF (serum response factor) [838] and TP53 [839] might play a central role in the development of chronic obstructive pulmonary disease. CAV1 [391], STAT1 [840] and NFKB1 [841] might be potential targets for the treatment of airway inflammation. CAV1 [626], TFAP2A [842], TP53 [843] and NFKB1 [844] have been identified as a new potential therapeutic target against systemic sclerosis. CAV1 [767], hsa-mir-19b-3p [845], JUN (Jun proto-oncogene, AP-1 transcription factor subunit) [846], STAT1 [847], TP53 [848] and NFKB1 [849] are reported to promote rheumatoid arthritis. The role of CAV1 [797] in the growth and progression of scleroderma. TTN (titin) [300] and STAT1 [850] plays an important role in the development of sarcoidosis. Documented altered expression of STAT1 [851] in dermatomyositis. Novel biomarkers include MAPT, GPR156, CCDC146, IQUB, MORN3, hsa-mir-6830-5p, hsa-mir-362-3p, hsa-mir-15b, hsa-mir-766-5p, hsa-mir-302a-3p, hsa-mir-4524a-3p, EN1 and ELK4 were found to be competitive endogenous RNAs whose expression intensity might predict the prognosis of IPF and its complications.

In summary, the current data provide a comprehensive bioinformatics analysis of DEGs that might be related to the progression of IPF and its complications. We have identified 958 candidate DEGs with NGS dataset and integrated bioinformatics analyses. A variety of novel genes and pathways might be involved in the molecular pathogenesis of IPF and its complications.

## Acknowledgement

I Katherine Monaghan, CSL Limited, Research and Development, Melbourne, Victoria, Australia, very much, the author who deposited their NGS dataset GSE213001, into the public GEO database.

## Ethical approval

This article does not contain any studies with human participants or animals performed by any of the authors.

## Informed consent

No informed consent because this study does not contain human or animals participants.

## Availability of data and materials

The datasets supporting the conclusions of this article are available in the GEO (Gene Expression Omnibus) (https://www.ncbi.nlm.nih.gov/geo/) repository. [(GSE213001) https://www.ncbi.nlm.nih.gov/geo/query/acc.cgi?acc=GSE213001]

## Consent for publication

Not applicable.

## Author Contributions

1. B. V. - Writing original draft, and review and editing
2. C. V. - Software and investigation

